# Behaviors of individual microtubules and microtubule populations relative to critical concentrations: Dynamic instability occurs when critical concentrations are driven apart by nucleotide hydrolysis

**DOI:** 10.1101/260646

**Authors:** Erin M. Jonasson, Ava J. Mauro, Chunlei Li, Ellen C. Norby, Shant M. Mahserejian, Jared P. Scripture, Ivan V. Gregoretti, Mark S. Alber, Holly V. Goodson

**Author notes:** Author for correspondence: Holly Goodson, Department of Chemistry and Biochemistry, 251 Nieuwland Science Hall, Notre Dame, IN 46556. Co-first authors (Erin M. Jonasson, Ava J. Mauro).

## Abstract

The concept of critical concentration (CC) is central to understanding behaviors of microtubules and other cytoskeletal polymers. Traditionally, these polymers are understood to have one CC, measured multiple ways and assumed to be the subunit concentration necessary for polymer assembly. However, this framework does not incorporate dynamic instability (DI), and there is work indicating that microtubules have two CCs. We use our previously established simulations to confirm that microtubules have (at least) two experimentally relevant CCs and to clarify the behaviors of individuals and populations relative to the CCs. At free subunit concentrations above the lower CC (CC_IndGrow_), growth phases of individual filaments can occur *transiently*; above the higher CC (CC_PopGrow_), the population’s polymer mass will increase *persistently*. Our results demonstrate that most experimental CC measurements correspond to CC_PopGrow_, meaning “typical” DI occurs below the concentration traditionally considered necessary for polymer assembly. We report that [free tubulin] at steady state does not equal CC_PopGrow_, but instead approaches CC_PopGrow_ asymptotically as [total tubulin] increases and depends on the number of stable microtubule seeds. We show that the degree of separation between CC_IndGrow_ and CC_PopGrow_ depends on the rate of nucleotide hydrolysis. This clarified framework helps explain and unify many experimental observations.

## INTRODUCTION

The concept of critical concentration (CC) is fundamental to experimental studies of biological polymers, including microtubules (MTs) and actin, because it is used to determine the amount of subunit needed to obtain polymer and to interpret the effects of polymer assembly regulators. In the standard framework for predicting the behavior of biological polymers, there is one critical concentration of subunits at which polymer assembly commences (e.g., (Alberts et al., 2015; Mirigian et al., 2013)). However, as indicated by other work (Hill and Chen, 1984; Walker et al., 1988), this framework fails to account for the dynamic instability (DI) displayed by microtubules and other DI polymers (e.g., PhuZ, ParM) (Mitchison and Kirschner, 1984a; Garner et al., 2004; Erb et al., 2014). One purpose of the work presented here is to examine the many experimental and theoretical definitions of CC in order to show how the definitions relate to each other. Another purpose is to clarify how the behaviors of individual dynamically unstable filaments and their populations relate to each other and to the experimental measurements of CC. To address these problems, we computationally modeled systems of dynamic microtubules with one of the two ends of each MT fixed (e.g., as occurs for MTs growing from centrosomes) and performed analyses that are directly comparable to those used in experiments. A significant advantage of computational modeling for this work is that it allows simultaneous examination of the behaviors of individual subunits, individual microtubules, and the population’s bulk polymer mass.

### Traditional understanding of Critical Concentration ( CC) based on equilibrium polymers

Traditionally, “the critical concentration” is understood to be the concentration of subunits needed for polymer assembly to occur (CC_PolAssem_, measured by Q1 in **Figure 1A,D**); equivalently, the CC is defined as the concentration of free subunits left in solution once polymer assembly has reached a steady-state level (CC_SubSoln_, measured by Q2 in **Figure 1A,D**). This set of ideas is based on early empirical observations with actin (Oosawa et al., 1959). These observations were initially given a theoretical framework by Oosawa and colleagues, who explained the behavior of actin by developing a theory for the equilibrium assembly behavior of helical polymers (Oosawa and Kasai, 1962; Oosawa, 1970). This equilibrium theory was extended to tubulin by Johnson and Borisy (Johnson and Borisy, 1975).

**Figure 1:**
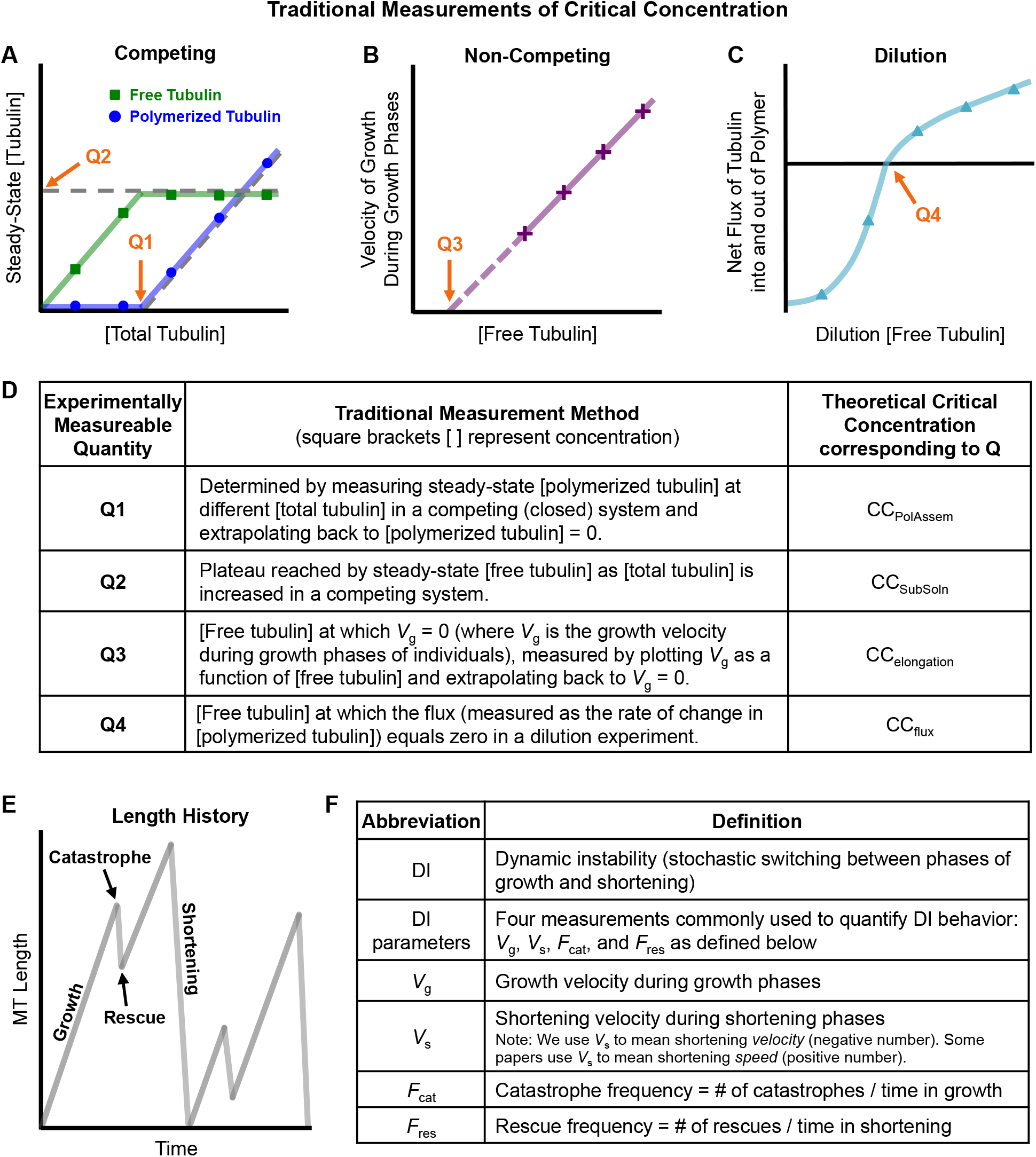
Classical understanding of microtubule (MT) polymer assembly behavior. See **Table 1** for additional description of the critical concentration measurements depicted here. [Free tubulin] is the concentration of tubulin dimers in solution, [polymerized tubulin] is the concentration of tubulin dimers in polymerized form, and [total tubulin] = [free tubulin] + [polymerized tubulin], ***(A)*** In a competing (closed) system, [total tubulin] is held constant over time and MTs compete for tubulin. As typically presented in textbooks, the critical concentration (CC) can be measured in a competing system by observing either the concentration of total tubulin at which MT polymer appears (**Q1**) or the concentration of free tubulin left in solution once the amount of polymer has reached steady state (**Q2**). ***(B)*** In a non-competing (open) system, [free tubulin] is held constant over time. In such a system, critical concentration is considered to be the minimum concentration of tubulin necessary for MT polymers to grow, which is estimated by measuring the growth rate of individual filaments (*V*_g_) and extrapolating back to *V*_g_ = 0 (**Q3**). ***(C)*** In dilution experiments, MTs are grown under competing conditions until the system reaches polymer-mass steady state, and then diluted into various [free tubulin]. The initial rate of change in [polymerized tubulin] is measured. Here, critical concentration is the concentration of dilution [free tubulin] at which the rate of change in [polymerized tubulin] is zero, (i.e., the dilution [free tubulin] at which the net flux of tubulin into and out of MT polymer is zero) (**Q4**). ***(D)*** Summary table of the definitions of the experimentally measureable quantities **Q1-4** depicted in panels **A-C**. ***(E)*** Individual MTs exhibit a behavior called dynamic instability (DI), in which the individuals undergo phases of growth and shortening separated by approximately random transitions termed catastrophe and rescue. ***(F)*** Table of definitions of DI parameters (four measurements commonly used to quantify DI behavior).

**Table 1:**
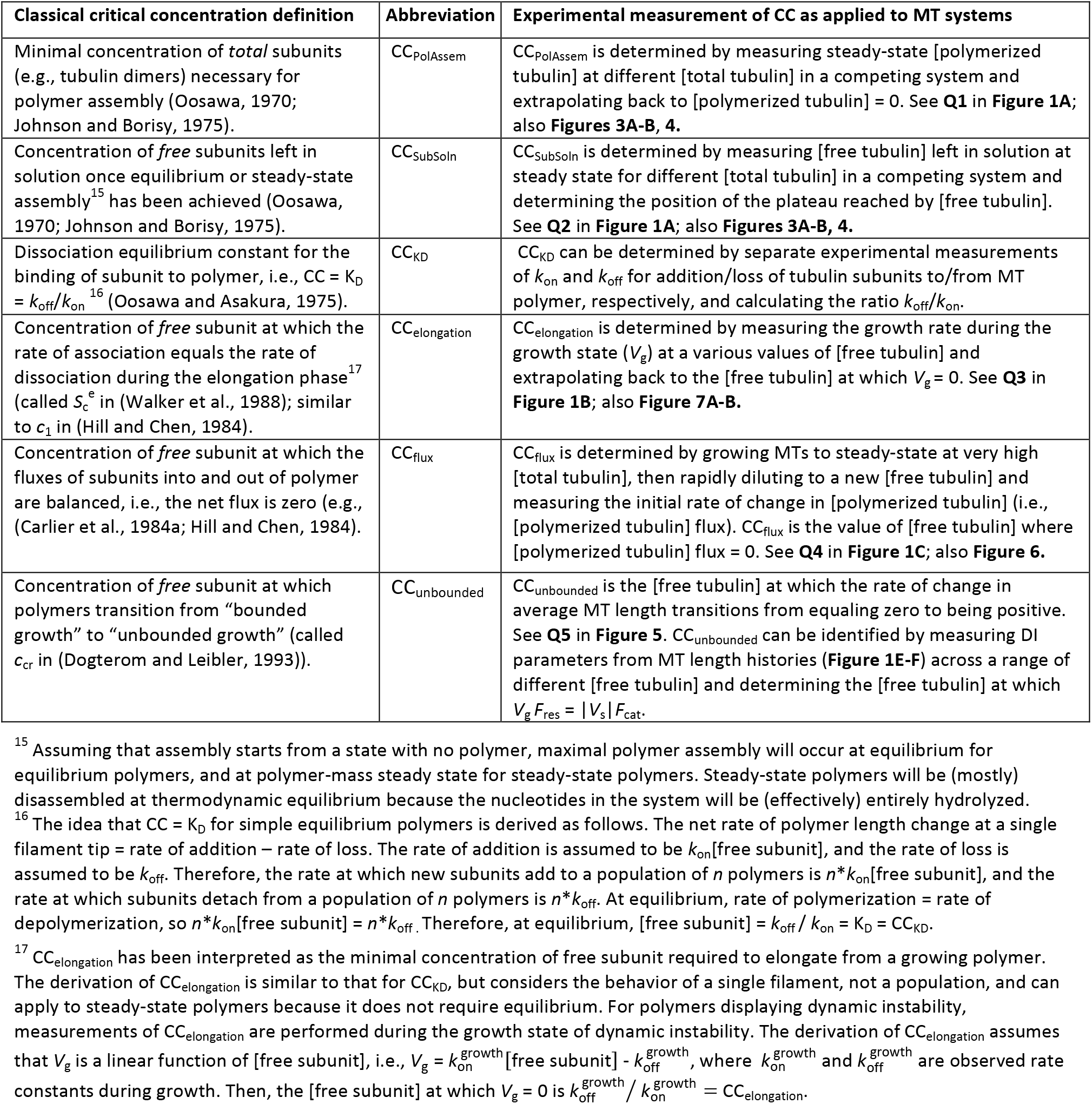
Traditional critical concentration (CC) definitions used in the literature. These definitions of critical concentration (CC) are interchangeable for equilibrium polymers, but have not all been compared in a single analysis for DI polymers. For each CC definition, we have assigned a specific abbreviation and provide an example of an early publication where that definition was used. The terms CC_PolAssem_, CC_SubSoln_, etc. refer to theoretical values (concepts), and Q1, Q2, etc. refer to experimentally measurable quantities (i.e., values obtained through experimental approaches as indicated in the figures). All definitions except CC_KD_ can be applied to both equilibrium and steady-state polymers (CC_KD_ assumes the system is at equilibrium and therefore can be applied to only equilibrium polymers). The traditional framework in **Table 1** will be revised in the Results Section (see **Table 3** for a summary).

For equilibrium polymers, the CC is commonly defined as *k*_off_/*k*_on_ = *K*_D_, where *k*_on_ and *k*_off_ are the rate constants for attachment/detachment of a subunit to/from a filament tip; polymer will then undergo net assembly when *k*_on_*[free subunit] is greater than *k*_off_ (**Table 1**). The idea that polymer assembly commences at the critical concentration is now used routinely to design and interpret experiments involving cytoskeletal polymers (e.g. (Amayed et al., 2002; Buey et al., 2005; Wieczorek et al., 2015; Díaz-Celis et al., 2017; Schummel et al., 2017; Concha-Marambio et al., 2017)), and it is a standard topic in cell biology textbooks (e.g., (Alberts et al., 2015; Lodish et al., 2016)). Over time, a set of experimental measurements and definitions of critical concentration have emerged (**Table 1, Figure 1**), all of which would be equivalent for an equilibrium polymer.

### Nucleotide hydrolysis allows microtubules to exhibit dynamic instability

Microtubules (composed of subunits called tubulin dimers) are steady-state polymers, not equilibrium polymers, because they require a constant input of energy in the form of GTP (guanosine triphosphate) nucleotides to maintain a (highly) polymerized state. Microtubules exhibit a behavior known as dynamic instability (DI), in which they stochastically switch between phases of growth and shortening via transitions known as catastrophe and rescue (**Figure 1E**) (Mitchison and Kirschner, 1984a; Walker et al., 1988). The DI behavior of MTs is driven by GTP hydrolysis (conversion of GTP-tubulin to GDP-tubulin): tubulin subunits containing GTP assemble into MTs, while tubulin subunits containing GDP do not (the *k*_on_ and *k*_off_ values for GTP-tubulin differ from those for GDP-tubulin). In contrast, tubulin subunits containing non- or slowly-hydrolyzable GTP analogs (e.g., GMPCPP) assemble into stable MTs that do not display DI (Hyman et al., 1992). Though some details about the mechanism of DI remain unclear, the consensus explanation for DI behavior is that growing MTs have a cap of GTP-tubulin subunits (the “GTP cap”) that stabilizes the underlying GDP-tubulin lattice. The MTs switch to rapid disassembly (i.e., undergo catastrophe) when they lose their stabilizing cap, exposing the unstable GDP-tubulin lattice below. When MTs regain their cap, they undergo rescue (transition from shortening to growth) (reviewed in (Goodson and Jonasson, 2018)).

On the surface, it may seem reasonable to apply the traditional critical concentration framework as outlined above (see also **Table 1**) to understanding DI polymers like microtubules because this framework is founded on theory (albeit equilibrium polymer theory) and appears to be consistent with many experimental results (Howard, 2001). A problem with this approach is that it leaves open various questions regarding how dynamic instability and energy utilization fit into the traditional CC framework. For example, how does the DI behavior of an individual filament in **Figure 1B,E** relate to the population-level behavior in **Figure 1A,C**? Is there one experimentally relevant critical concentration (as assumed from equilibrium polymer theory) or more than one? If more than one, how many? More broadly, why do some steady-state polymers (e.g., microtubules) display dynamic instability, while others (e.g., actin) do not?

As one might imagine, these questions have been studied previously, but ambiguity in understanding critical concentration still exists. A brief summary of some key previous efforts on CC for MTs is as follows:

- In the 1980s, Hill and colleagues investigated some of the questions outlined above and worked to develop a theory of steady-state polymer assembly. Their conclusions included the idea that growth of microtubules is governed by two distinct critical concentrations: a lower CC at which “the mean subunit flux per polymer” during “phase 1” (growth phase) equals zero and an upper CC at which “the mean net subunit flux per polymer” is zero (similar to **Figure 1C**) (e.g., (Hill and Chen, 1984), elaborated on in (Hill, 1987)). However, the published work did not clarify for readers the biological significance of these two CCs nor how they relate to the behaviors of individual filaments and their populations.
- Later in the 1980s, Walker et al. used video microscopy to analyze in detail the behavior of individual MTs undergoing dynamic instability. They demonstrated that microtubules observed *in vitro* have a “critical concentration for elongation” (CC_elongation_), which they described as the concentration at which the rate of tubulin association 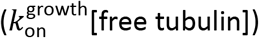 is equal to the rate of dissociation 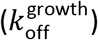 during the elongation phase (Walker et al., 1988) (**Figure 1B; Table 1** and its footnotes). Consequently, at tubulin concentrations below CC_elongation_, there is no elongation. Later in this same paper, the authors discussed the existence of a higher critical concentration above which a population of polymers will undergo “net assembly”. Thus, the analysis in this manuscript clearly indicates that microtubules have two critical concentrations. However, this conclusion is not stated explicitly, and the manuscript does not address the question of how either of the two Walker et al. CCs relates to the two CCs predicted by Hill.
- In the 1990s, Dogterom et al. and Fygenson et al. used a combination of modeling (Dogterom and Leibler, 1993) and experiments (Fygenson et al., 1994) to show that there is a “critical value of monomer density, c = c_cr_”, above which microtubule growth is “unbounded” (i.e., the average length increases indefinitely and does not level off with time) (Dogterom and Leibler, 1993; Fygenson et al., 1994; Dogterom et al., 1995). Hereafter, we refer to this c_cr_ as CC_unbounded_. Dogterom et al. also provided equations (similar to those proposed initially by (Hill and Chen, 1984) and (Walker et al., 1988)) that can be used to relate CC_unbounded_, which is a population-level characteristic, to the dynamic instability parameters (**Figure 1E-F**), which describe individual-level behaviors. One of the many significant outcomes of these papers was that they encouraged readers to think about how small changes to DI parameters (e.g., as caused by regulatory changes to MT binding proteins) could change the behavior of a system of MTs, especially in a cellular context. However, the implications of these articles for understanding critical concentrations more broadly remained poorly appreciated because they did not explicitly relate CC_unbounded_ to the more classical CC definitions and measurements in **Table 1** or to those discussed by (Hill and Chen, 1984) and (Walker et al., 1988).

Thus, although dynamic instability has been studied for more than 30 years, confusion remains about how the traditionally equivalent definitions of critical concentration and the interpretation of CC measurements should be adjusted to account for dynamic instability. Remarkably, the literature as yet still lacks a clear discussion of how the CC_elongation_ and CC_unbounded_ mentioned above relate to each other, to the CCs predicted by Hill, or to the classical experimental measurements of CC depicted in **Figure 1A**. How many distinct CCs are produced by the different experimentally measurable quantities (Q values, **Figure 1** and **Table 1**), what is the practical significance of each, and which measurements yield which CC? How do any of these values relate to behaviors at the scales of subunits, individual microtubules, and the bulk polymer mass of populations of microtubules? How does dynamic instability behavior relate to the separation between distinct CCs? Undoubtedly, many researchers have an intuitive understanding of the answers to at least some of these questions. However, the observation that even recent literature contains many references to “the” CC for microtubule assembly (e.g., (Wieczorek et al., 2015; Alfaro-Aco and Petry, 2015; Hussmann et al., 2016; Schummel et al., 2017) indicates that this problem deserves attention. While these issues are interesting from a basic science perspective, they also have significant practical relevance: proper design and interpretation of experiments that involve perturbing microtubule dynamics (e.g., characterization of MT-directed drugs or proteins) requires an unambiguous understanding of critical concentrations and how they are measured (e.g., (Verdier-Pinard et al., 2000; Bonfils et al., 2007; Hussmann et al., 2016; Cytoskeleton Inc.).

### Computer simulations as an approach to addressing these questions

To investigate the concept of critical concentration as it applies to dynamically unstable polymers, we have used computational modeling. Computational models are ideal for addressing this type of problem because the biochemistry of the reactions can be explicitly controlled and “experiments” can be performed quickly and easily. Furthermore, it is possible to simultaneously follow the behavior of the system at all relevant scales: addition/loss of individual subunits to/from the free end, dynamic instability of individual filaments, and any changes in polymer mass of the population of filaments. In comparison, it is challenging to address these questions using physical systems because experiments have thus far been limited technically to measurements at one (or at most two) of these scales at a time.

As described more below (Results subsection “Computational Models” and **Figure 2**), the rules and conditions controlling our simulations correspond to those that would be set by the intrinsic properties of the biological system (e.g., kinetic rate constants) or by the experimenter (e.g., total tubulin concentration). Typical experimental results (DI parameters, concentrations of free and polymerized tubulin) are emergent properties of the system of biochemical reactions, just as they would be in a physical experiment.

**Figure 2:**
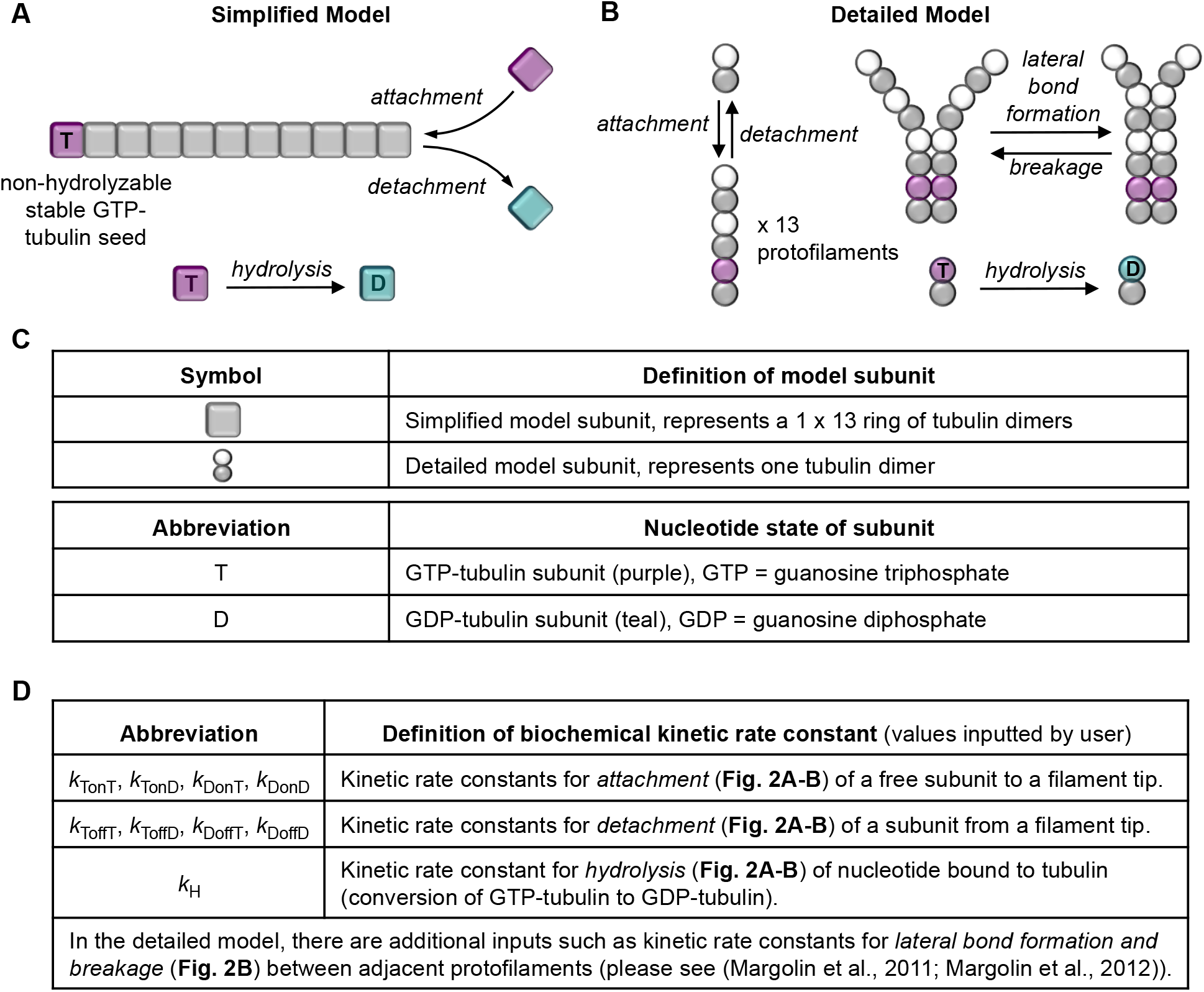
Processes that occur in the computational models. ***(A)*** In the <u>simplified model</u>, microtubules are approximated as simple linear filaments that can undergo three processes: subunit addition, loss, and hydrolysis. Addition and loss can occur only at the tip. Hydrolysis can occur anywhere in the filament where there is a GTP-subunit. ***(B)*** In the <u>detailed model</u>, there are 13 protofilaments, which each undergo the same processes as in the simplified model but also undergo lateral bonding and breaking between adjacent protofilaments. (***C***) Information about the subunits in the models. In both models, the kinetic rate constants (panel **D**) controlling these processes are inputted by the user, and the MTs grow off of a user-defined constant number of stable MT seeds (composed of non-hydrolyzable GTP-tubulin). The standard dynamic instability parameters (*V*_g_, *V*_S_, *F*_cat_, *F*_res_; see **Figure 1E-F**) are emergent properties of the input rate constants, [free tubulin], and other aspects of the environment such as the number of stable seeds. For more information about the models and their parameter sets, see **Box 1**, Methods, Supplemental Methods, and (Gregoretti et al., 2006; Margolin et al., 2011; Margolin et al., 2012).

### Summary of Conclusions

Using these systems of simulated microtubules, we show that classical interpretations of experiments such as those in **Figure 1** can be misleading in terms of understanding the behavior of individual MTs. In particular, we use the simulations to illustrate the fact that dynamically unstable polymers like microtubules do have (at least) two major experimentally distinguishable critical concentrations, as originally proposed by Hill and colleagues (summarized in (Hill, 1987)). We clarify how the CCs relate to behaviors of individual MTs and populations of MTs. At [free tubulin] above the lower CC, extended growth phases of *individual* filaments *can* occur *transiently*. At [free tubulin] above the higher CC, the polymer mass of a large *population will* increase *steadily*, even while individual filaments in the population potentially still exhibit dynamic instability. We show that the lower critical concentration corresponds to CC_elongation_ as measured by Walker et al. (**Table 1**; (Walker et al., 1988)), which can be described as the free tubulin concentration above which individual MTs are able to elongate during the growth phase. This CC can be measured by experimental quantity Q3 in **Figure 1B**. The higher CC corresponds to CC_unbounded_ as identified by Dogterom et al., i.e., the concentration of free tubulin above which “unbounded growth” occurs (**Table 1**; (Dogterom and Leibler, 1993; Fygenson et al., 1994; Dogterom et al., 1995). This upper CC can be measured by Q1, Q2, and Q4 in **Figure 1A,C**. To clearly distinguish these two CCs and avoid confusing either with a situation where a physical boundary is involved, we suggest calling them CC_IndGrow_ and CC_PopGrow_, respectively.^1^ In addition to these two experimentally accessible CCs, there are two more CCs (perhaps not experimentally accessible) that correspond to the K_D_ for the GTP and GDP forms of tubulin subunits. We suggest calling these CC_KD_GTP_ and CC_KD_GDP_, respectively. While our studies focus on microtubules, we suggest that these critical concentration definitions and interpretations can apply to steady-state polymers more generally but are especially significant for those that exhibit dynamic instability.

We show that most experiments intended to measure “the CC” actually measure CC_PopGrow_ (i.e., the higher CC). This conclusion means that “typical” microtubule dynamic instability (where MTs grow and depolymerize back to the seed) is limited to concentrations *below* what has traditionally been considered “the” CC needed for polymer assembly. Furthermore, we show that in competing systems (i.e., closed systems where MTs compete for a limited total number of tubulin subunits), the concentration of free tubulin at steady state ([free tubulin]_SteadyState_) does not *equal* CC_PopGrow_ as would be expected from traditional interpretations of classic CC experiments (**Figure 1A**). Instead, [free tubulin]_SteadyState_ asymptotically *approaches* CC_PopGrow_ as [total tubulin] increases. In addition, we demonstrate that the degree of separation between CC_IndGrow_ and CC_PopGrow_ depends on the GTP hydrolysis rate constant (*k*_H_). We also show that CC_IndGrow_ can differ from CC_KD_GTP_, contrary to previous assumptions that growing MTs always have GTP-tubulin at their tips (topmost subunits) (e.g, (Bowne-Anderson et al., 2015)). Finally, we demonstrate that dynamic instability itself can produce results (e.g., sigmoidal seed occupancy plots) previously interpreted as evidence that growth from stable seeds requires a nucleation step.

## RESULTS

### Computational Models

In this work, we used both a “simplified” model of MT dynamics, in which MTs are modeled as simple linear polymers (Gregoretti et al., 2006), and a “detailed” model, where microtubules are composed of 13 protofilaments, with lateral and longitudinal bonds between subunits (tubulin dimers) modeled explicitly (Margolin et al., 2011; Margolin et al., 2012) (**Figure 2**). The simulations were designed to be intuitively understandable to researchers familiar with biochemical aspects of cytoskeletal polymers. Consequently, the rules governing the simulations correspond directly to biochemical reaction kinetics. Key elements of these models are described in **Box 1**.

We utilize both the simplified and detailed computational models because each has particular strengths for addressing problems related to microtubule dynamics. The simplified simulation has fewer kinetic parameters, all of which are directly comparable to parameters in typical analytical models (i.e., mathematical equations). Thus, the simplified simulation is useful for testing analytical model predictions relating biochemical properties to individual filament level and bulk population level behaviors. In contrast, the increased resolution of the detailed model is important for testing the generality and relevance of conclusions derived from the simplified model.

In addition, the inputted kinetic rate constants in the two models were tuned to produce dynamic instability behavior that is *quantitatively* different between the two models, so it follows that the specific numerical values for critical concentrations extracted from these two simulations will be different. However, as discussed more below, the behavioral changes that occur at each CC are *qualitatively* similar in the two models. Thus, these two models enable us to determine which conclusions are general and to avoid making conclusions that are specific to particular parameter sets or polymer types.

##### Box 1: Key elements of the two computational models (simplified and detailed) used in this study

- Subunit addition/loss and GTP hydrolysis (both models) and lateral bond formation/breaking (detailed model only) are modeled as stochastic events that occur according to kinetic rate equations based on the biochemistry of these processes (**Figure 2**) (Gregoretti et al., 2006; Margolin et al., 2012).
- The user-defined (adjustable) parameters correspond to the following: (a) the biochemistry of the proteins being studied (i.e., kinetic rate constants for the reactions listed above); and (b) attributes of the environment that would be set by either the experimenter or the cell (e.g., the concentration of tubulin in the system, whether the system is competing (closed) or non-competing (open), the number of stable seeds, and the system volume).
- As in physical experiments, emergent properties of the simulated systems include the dynamic instability parameters (*V*_g_, *V*_s_, *F*_cat_, *F*_res_, see **Figure 1E-F**) and the concentrations of free and polymerized tubulin at steady state. In particular, transitions between growth and shortening (catastrophe and rescue) are spontaneous processes that occur when the stabilizing GTP cap happens to be lost or regained as a result of the biochemical reactions described above.
- Because microtubules in cells and in many *in vitro* experiments grow from stable seeds (nucleation sites such as centrosomes, axonemes, or GMPCPP seeds), our simulations assume that one end of each MT is fixed (as would be the case for growth from centrosomes), and that all addition and loss occur at the free end. In our simulations, the seeds are composed of non-hydrolyzable GTP-tubulin. Except where otherwise noted, the number of stable seeds was set to 100 in the simplified model and 40 in the detailed model.
- Both simulations spontaneously undergo the full range of dynamic instability behaviors (including rescue), and they can simulate systems of dynamic microtubules for hours of simulated time (Gregoretti et al., 2006; Margolin et al., 2012).
- The behaviors of the evolving systems of dynamic microtubules can be followed at the scales of subunits, individual filaments, or populations of filaments.
- The kinetic rate constants used as input parameters for the detailed model were previously tuned to approximate the DI parameters of mammalian brain MTs *in vitro* (Margolin et al., 2012). The simplified model parameters used here are modified from those of (Gregoretti et al., 2006) and were chosen for use here because they produce DI behavior that is quantitatively different from that of the detailed model.

The sum of these attributes make these simulations ideal for studying the relationships between the concentration of tubulin, the behaviors of individual MTs, and behaviors of systems of dynamic MTs. See the Methods, Supplementary Information, and (Gregoretti et al., 2006; Margolin et al., 2012) for additional details including input parameters.

### Approach to understanding the relationship between microtubule behaviors and critical concentrations

To clarify the concept of critical concentration as it applies to microtubules, we examined which of the commonly used critical concentration definitions (outlined in **Table 1**) are meaningful when studying microtubules, and for the set that are meaningful, which are equivalent. The term critical concentration can have a specific thermodynamic meaning as the solute concentration at which a phase change occurs. Here we define the term operationally, as the concentration at which a behavioral change occurs.

To determine how the various CC definitions relate to each other and to dynamic instability, we used the simulations to simultaneously examine the behaviors of individual MTs and populations of MTs. More specifically, we ran sets of simulations for both the simplified and detailed models at various tubulin concentrations in both competing systems (closed systems with constant [total tubulin], as might happen in a test tube) and non-competing systems (open systems with constant [free tubulin], similar to what might happen in a microscope flow cell). This approach mimics various experiments (**Table 2**) that are classically used to measure microtubule critical concentration (**Table 1**). We then assessed and compared the behaviors of the individual microtubules (e.g., DI parameters), population-level properties (e.g., [free tubulin] at steady state), and critical concentrations as determined by the traditional definitions (**Table 1**).

**Table 2:**
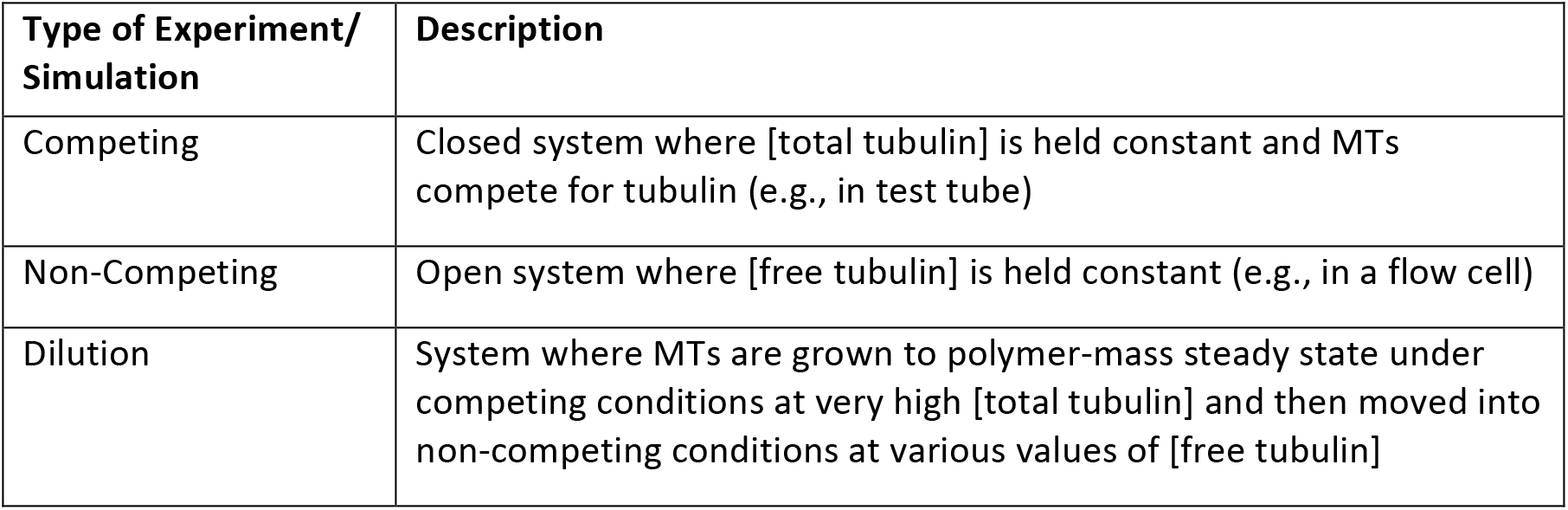
Types of Experiments/Simulations.

For the work presented here, it is important to recognize that the relevant observations are the *behaviors* of the systems at different scales and the concurrence (or disagreement) between the values of CC that result from various definitions or measurement approaches; the specific numerical CC values observed are simply outcomes of the particular input kinetic rate constants used and so are not by themselves significant. This situation is analogous to physical MTs, where DI parameters and CC values depend on the protein sequences, temperatures, and buffer conditions used (e.g., (Williams et al., 1985; Gildersleeve et al., 1992; Fygenson et al., 1994; Hussmann et al., 2016; Schummel et al., 2017)).

We use the terms Q1, Q2, etc. to refer to specific experimentally measurable quantities (i.e., values obtained through experimental approaches as indicated in the figures), and the terms CC_KD_, CC_PolAssem_, CC_SubSoln_, etc. to refer to theoretical values (concepts) that may or may not correspond to particular experimentally measurable quantities and may or may not be equivalent. **Table 1** summarizes traditional critical concentration definitions and measurements used in the literature. **Table 3** summarizes our clarifications of critical concentration definitions and additional Q value measurements based on the results that will be presented in this work.

### Addressing the idea that “the” critical concentration is *k*_off_/*k*_on_ (CC_KD_)

The idea that “the CC” is the *K*_D_ for addition of subunits to polymer (i.e., CC = *k*_off_/*k*_on_ = *K*_D_; CC = CC_KD_; **Table 1**) is a serious oversimplification when applied to microtubules. Though this formula is frequently stated in textbooks, it is well-recognized that this relationship cannot be applied in a straightforward way to populations of dynamic microtubules, or to steady-state polymers more generally (Alberts et al., 2015). More specifically, experimentally observed critical concentrations for systems of dynamic MTs (however measured) cannot be equated to simple *k*_off_/*k*_on_ = *K*_D_ values because the GTP and GDP forms of tubulin have significantly different values of *k*_off_/*k*_on_. For example, the critical concentration for GMPCPP (GTP-like) tubulin has been reported to be less than 1 μM (Hyman et al., 1992), while that for GDP-tubulin is very high, perhaps immeasurably so (Howard, 2001).

Exactly how the measured CC value(s) for a system of dynamic microtubules relate to the *K*_D_ values for GTP- and GDP-tubulin has not previously been established. However, intuition suggests that any CCs must lie between the respective *K*_D_ values for GTP- and GDP-tubulin (Howard, 2001). Consistent with this idea, experimentally reported values for mammalian brain tubulin CC typically lie between ~1 and ~20 μM (Verdier-Pinard et al., 2000; Bonfils et al., 2007; Mirigian et al., 2013; Wieczorek et al., 2015). Note that while the idea that CC = *K*_D_ cannot apply in a simple way to a system of dynamic microtubules, it can apply to tubulin polymers in the absence of hydrolysis, where assembly is an equilibrium phenomenon. Examples include systems containing only GDP-tubulin (when polymerized with certain drugs) or tubulin bound to non-/slowly-hydrolyzable GTP analogs (e.g., GTP-γS, GMPCPP) (Hyman et al., 1992; Díaz et al., 1993; Buey et al., 2005).

### DI polymers grow at concentrations below standard experimental quantities commonly thought to measure the critical concentration for polymer assembly

A typical way to measure “the critical concentration” for microtubule assembly is to determine the [total tubulin] at which polymer assembles in a competing (closed) experiment such as that portrayed in **Figure 1A**, where Q1 measures the CC for polymer assembly (CC_PolAssem_) (see e.g., (Mirigian et al., 2013)). An alternative approach treated as equivalent is to measure the concentration of free tubulin left in solution once steady-state polymer assembly has occurred (**Figure 1A**, Q2), traditionally considered to yield CC_SubSoln_ (Mirigian et al., 2013). In other words, the expectation is that Q1 ≈ Q2, and that these experimentally obtained quantities provide equivalent ways to measure the critical concentration for polymer assembly, where CC_PolAssem_ = CC_SubSoln_ (**Table 1**).

We tested these predictions by performing simulations of competing systems where individual MTs growing from stable seeds compete for a limited pool of tubulin (i.e., [total tubulin] is constant). This situation is analogous to a test-tube experiment in which microtubules grow from pre-formed MT seeds, and both [polymerized tubulin] and [free tubulin] are measured after the system has reached polymer-mass steady state (**Figure S1A-D**).^2^ At first glance, the behavior of the systems of simulated microtubules might seem consistent with that expected from common understanding (**Figure 1A**): significant polymer assembly was first observed at [total tubulin] ≈ Q1, and Q1 ≈ Q2 (**Figure 3A-B**).

**Figure 3:**
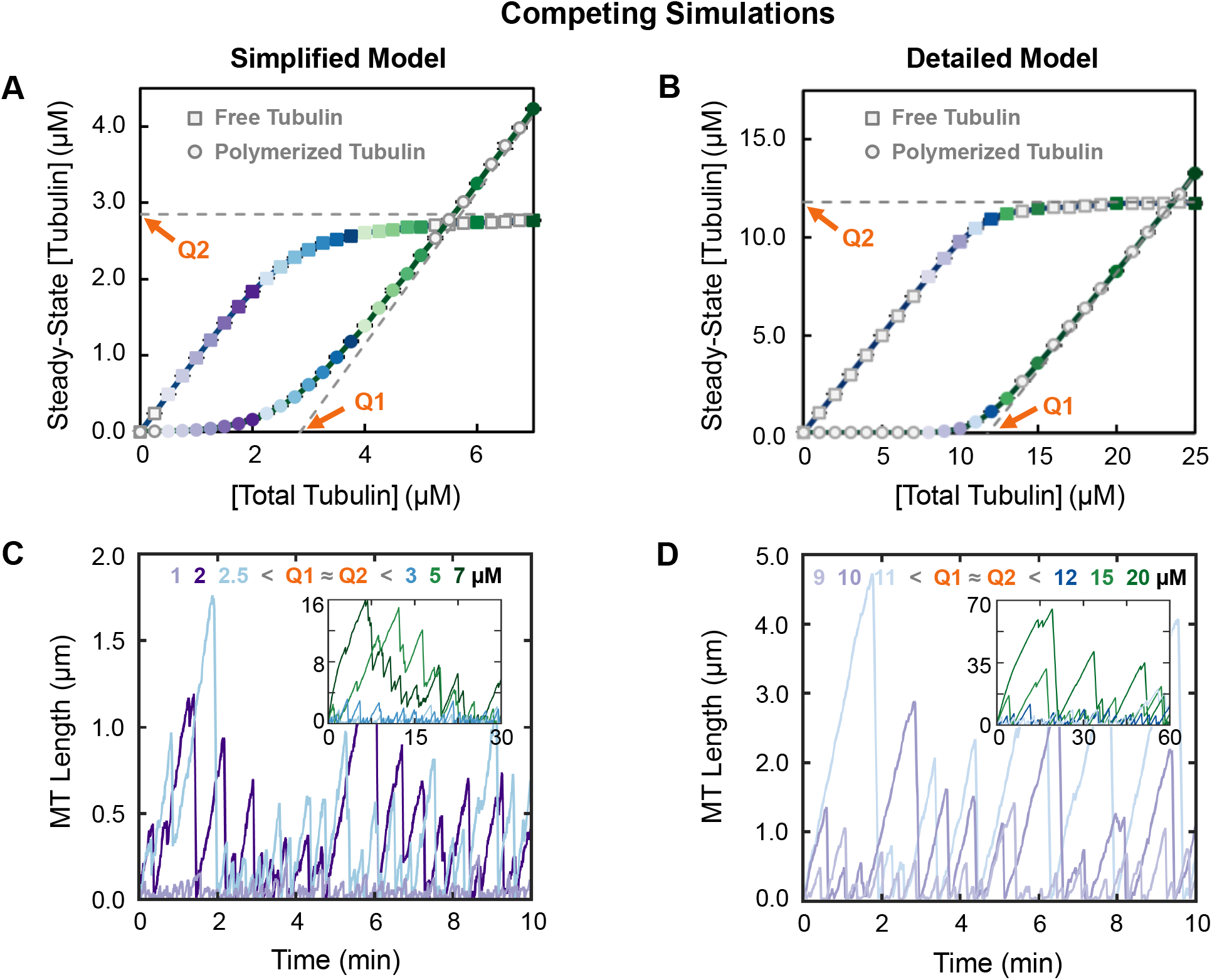
Behavior of microtubules (populations and individuals) under conditions of constant total tubulin. Left panels: simplified model; right panels: detailed model; colors of data points reflect the concentrations of *total* tubulin. (***A,B***) Classical critical concentration measurements (compare to **Figure 1A**). Systems of competing MTs at total tubulin concentrations as indicated on the horizontal axis were each allowed to reach polymer-mass steady state (shown in **Figure S1A-D**). Then the steady-state concentrations of free (squares) and polymerized (circles) tubulin were plotted as functions of [total tubulin]. (***C,D***) Representative length history plots for individual MTs from the simulations used in panels **A-B.** The value of [total tubulin] for each length history is indicated in the color keys at the top of panels **C-D**. ***Interpretation***: Classically, **Q1** estimates CC_PolAssem_, and **Q2** estimates CC_SubSoln_. However, as can be seen in panel **C-D,** MTs grow in both simulations at [total tubulin] below **Q1 ≈ Q2** (~2.85 μM in the simplified model and ~11.8 μM in the detailed model). Consistent with this observation, the main text provides justification for the idea that CC as estimated by **Q1 ≈ Q2** instead measures CC_PopGrow_, the CC for persistent population growth. Note that the difference in the values of **Q1 ≈ Q2** between the two simulations is expected from the fact that the inputted kinetic parameters for the simulations were chosen to produce quantitatively different DI measurements in order to provide a test of the generality of conclusions about qualitative behaviors; the results show that the behaviors are indeed qualitatively similar between the two simulations. For additional data related to these simulations (e.g., plots of [free tubulin] and [polymerized tubulin] as functions of time), see **Figure S1**. ***Methods***: Data points in panels **A-B** represent the mean +/− one standard deviation of the values obtained in three independent runs of the simulations. The values from each of three runs are averages over 15 to 30 minutes for the simplified model (panel **3A**) and over 30 to 60 minutes for the detailed model (panel **3B**). These time periods were chosen so that [free tubulin] and [polymerized tubulin] have reached their steady-state values (**Figure S1A-D**).

However, closer examination of these data shows that there is no sharp transition at either Q1 or Q2 (**Figure 3A-B**), as traditionally expected (**Figure 1A**). Significantly, small but non-zero amounts of polymer exist at [total tubulin] below reasonable estimates for Q1 (**Figure 3A-B**, **S1E-F**). In addition, the steady-state concentration of free tubulin ([free tubulin]_SteadyState_) is not constant with respect to [total tubulin] for [total tubulin] > Q1 as is often assumed. Instead, [free tubulin]_SteadyState_ approaches an asymptote represented by Q2 (**Figure 3A-B**). Nonetheless, Q1 is still approximately equal to Q2.^3^ Consistent with these observations, examination of individual MTs in these simulations shows MTs growing and exhibiting dynamic instability at [total tubulin] below Q1 ≈ Q2 (**Figure 3C-D**; compare to **Figure 3A-B**).

These data (**Figure 3**) suggest that one of the most commonly accepted predictions of traditional critical concentration understanding is invalid when applied to systems of dynamic microtubules: instead of both Q1 and Q2 providing an experimental measure of the minimum concentration of tubulin needed for polymer assembly (CC_PolAssem_), *neither* does, since MTs exhibiting dynamic instability appear at concentrations below Q1 ≈ Q2. Correspondingly, the results in **Figure 3A-B** indicate that the critical concentration called CC_SubSoln_ would be more accurately defined as the asymptote approached by the [free tubulin]_SteadyState_ as [total tubulin] increases, not the value of [free tubulin]_SteadyState_ itself (**Figure 1A**).

### The number of stable MT seeds impacts the sharpness of the transition at Q1 and Q2

Why is the transition at Q1 and Q2 in **Figure 3A-B** more gradual than the theoretical transition as depicted in **Figure 1A**? Previous results of our simplified model (Gregoretti et al., 2006) and other models (e.g., (Vorobjev and Maly, 2008; Mourão et al., 2011)) indicate that [free tubulin]_SteadyState_ depends on the number of stable MT seeds. Therefore, we investigated how changing the number of stable MT seeds affects the shape of the curves in classical CC plots. Examination of the results (**Figure 4A-B**, zoom-ins in **Figure 4C-D**) shows that changing the number of MT seeds does change the sharpness of the transition at Q1 and Q2. More specifically, when the number of MT seeds is small, a relatively sharp transition is seen at both Q1 and Q2 in graphs of steady-state [free tubulin] and [polymerized tubulin]; little if any bulk polymer is observed at [total tubulin] below Q1 (**Figure 4**, fewer seeds, darkest curves; similar to **Figure 1A**). In contrast, when the number of MT seeds is high, measurable amounts of polymer appear at concentrations well below Q1, and consequently [free tubulin]_SteadyState_ approaches the Q2 asymptote more gradually (**Figure 4**, more seeds, lightest curves). Moreover, the data for various numbers of seeds all approach the same asymptotes (grey dashed lines, **Figure 4**). These observations indicate that the number of MT seeds does not impact the value of Q1 ≈ Q2, but does affect how sharply steady-state [free tubulin] approaches the Q2 asymptote.

**Figure 4:**
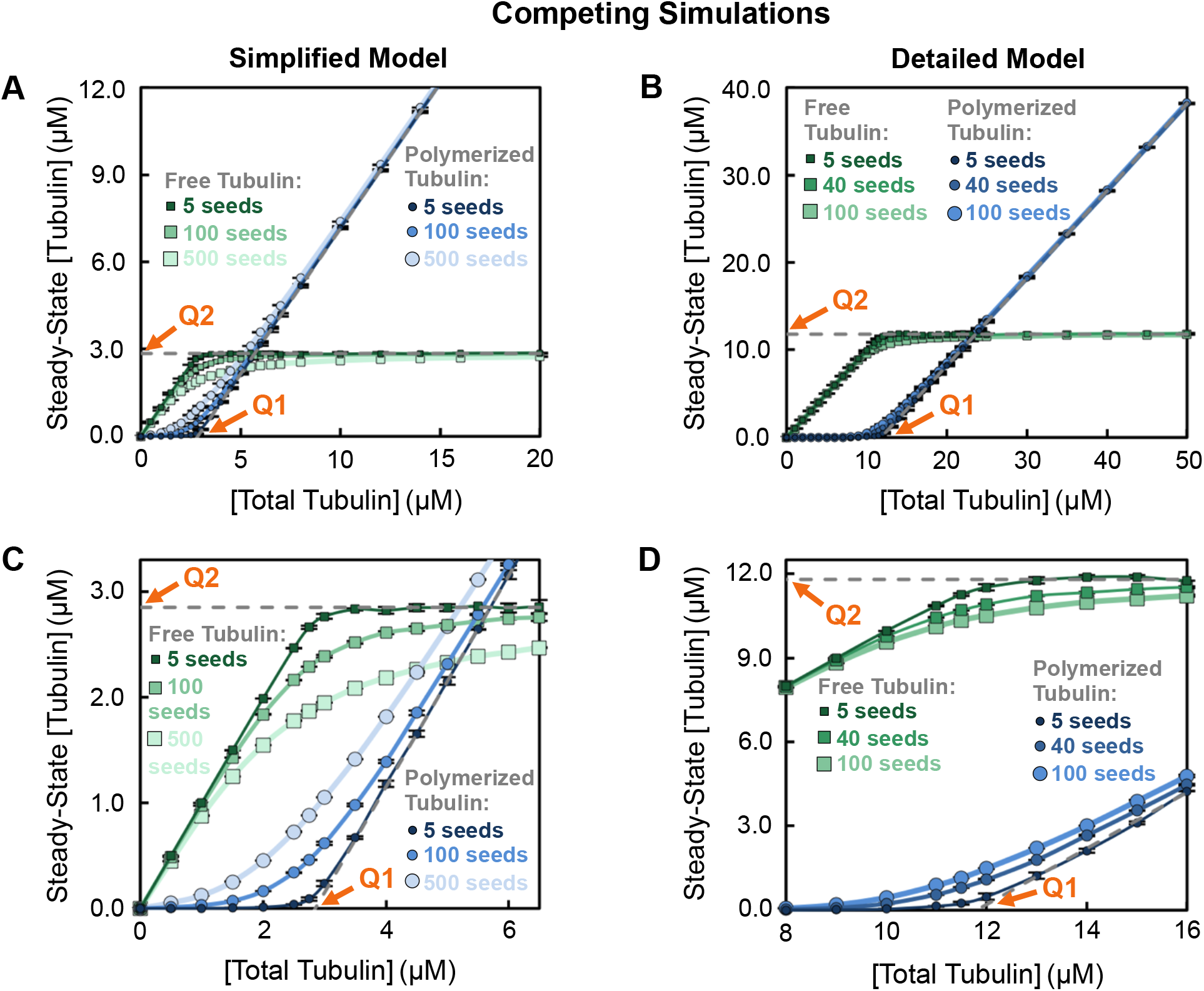
Impact of changing the number of microtubule seeds. Steady-state concentrations of free (squares) and polymerized (circles) tubulin in a competing system as in **Figure 3A-B**. ***(A,C)*** Simplified model with MTs growing from 5, 100, or 500 stable MT seeds (data for 100 seeds re-plotted from **Figure 3A**). ***(B,D)*** Detailed model with MTs growing from 5, 40, or 100 stable MT seeds (data for 40 seeds re-plotted from **Figure 3B**). Panels 4C-D show zoom-ins of the data plotted in panels **4A-B**, respectively. The darker curves with smaller symbols correspond to fewer seeds and the lighter curves with larger symbols correspond to more seeds. ***Interpretation***: These data show that changing the number of stable MT seeds alters the approach to the asymptotes determining Q1 and Q2 (dashed grey lines re-plotted here from **Figure 3A-B**), but does not change the value of **Q1 ≈ Q2**. ***Methods***: Data points represent the mean +/− one standard deviation of the values obtained in three independent runs of the simulations. Similar to **Figure 3**, [free tubulin] and [polymerized tubulin] from each run were averaged over a period of time after polymer-mass steady state was reached. The time to reach this steady state depends on the number of stable MT seeds (see **Figure S2**). For the simplified model, the averages of [free tubulin] and [polymerized tubulin] were taken from 120 to 150 minutes for 5 MT seeds and from 15 to 30 minutes for 100 and 500 MT seeds. For the detailed model, the averages were taken from 100 to 150 minutes for 5 MT seeds and from 30 to 60 minutes for 40 and 100 MT seeds. We were able to use a higher number of seeds in the simplified model than in the detailed model because it is more computationally efficient.

The observations thus far raise a question: Since CC_SubSoln_ is not the minimum tubulin concentration needed for polymer assembly (CC_PolAssem_), what is the significance of Q1 ≈ Q2 ≈ CC_SubSoln_ for microtubule behavior?

### A critical concentration for persistent growth of MT populations (CC_PopGrow_)

To investigate the significance of Q2, i.e., the asymptote approached by [free tubulin]_SteadyState_ as [total tubulin] is increased (**Figures 3A-B, 4**), we examined the dependence of MT behavior on the concentration of free tubulin in non-competing simulations. For these studies, we fixed [free tubulin] at various values instead of allowing polymer growth to deplete the free tubulin over time. This set of conditions is analogous to a laboratory experiment involving MTs polymerizing from stable seeds in a constantly replenishing pool of free tubulin at a known concentration, such as might exist in a flow cell.

As described above, Q1 and Q2 from competing systems do not yield the critical concentration for polymer assembly (CC_PolAssem_) as expected from traditional understanding. Instead, comparison with the non-competing simulations (**Figure 5**) shows that Q1 and Q2 correspond to a different CC, which can be described as the [free tubulin] above which individual MTs will exhibit *net growth* over long periods of time (**Figure 5A-B**). Equivalently, this CC can be described as the [free tubulin] above which the polymer mass of a large population of MTs will grow persistently (we use this terminology based on the experimentally-observed “persistent growth” in (Komarova et al., 2002)). As discussed more below, this CC is the same as that previously identified by Dogterom et al. as the CC at which the transition from “bounded growth” to “unbounded growth”^4^ occurs (Dogterom and Leibler, 1993; Fygenson et al., 1994; Dogterom et al., 1995), by Walker et al. as the CC for “net assembly” (Walker et al., 1988), and by Hill et al. as the CC where net subunit flux equals zero (Hill and Chen, 1984). To avoid implying that a physical boundary is involved, we suggest identifying this CC as the critical concentration for persistent microtubule population growth (CC_PopGrow_). CC_PopGrow_ can be measured by Q5a, the [free tubulin] at which the net rate of change in average MT length (**Figure 5C-D**, left axes) or in polymer mass (**Figure 5C-D**, right axes) transitions from equaling zero to being positive. Additional approaches to measuring CC_PopGrow_ are discussed later.

**Figure 5:**
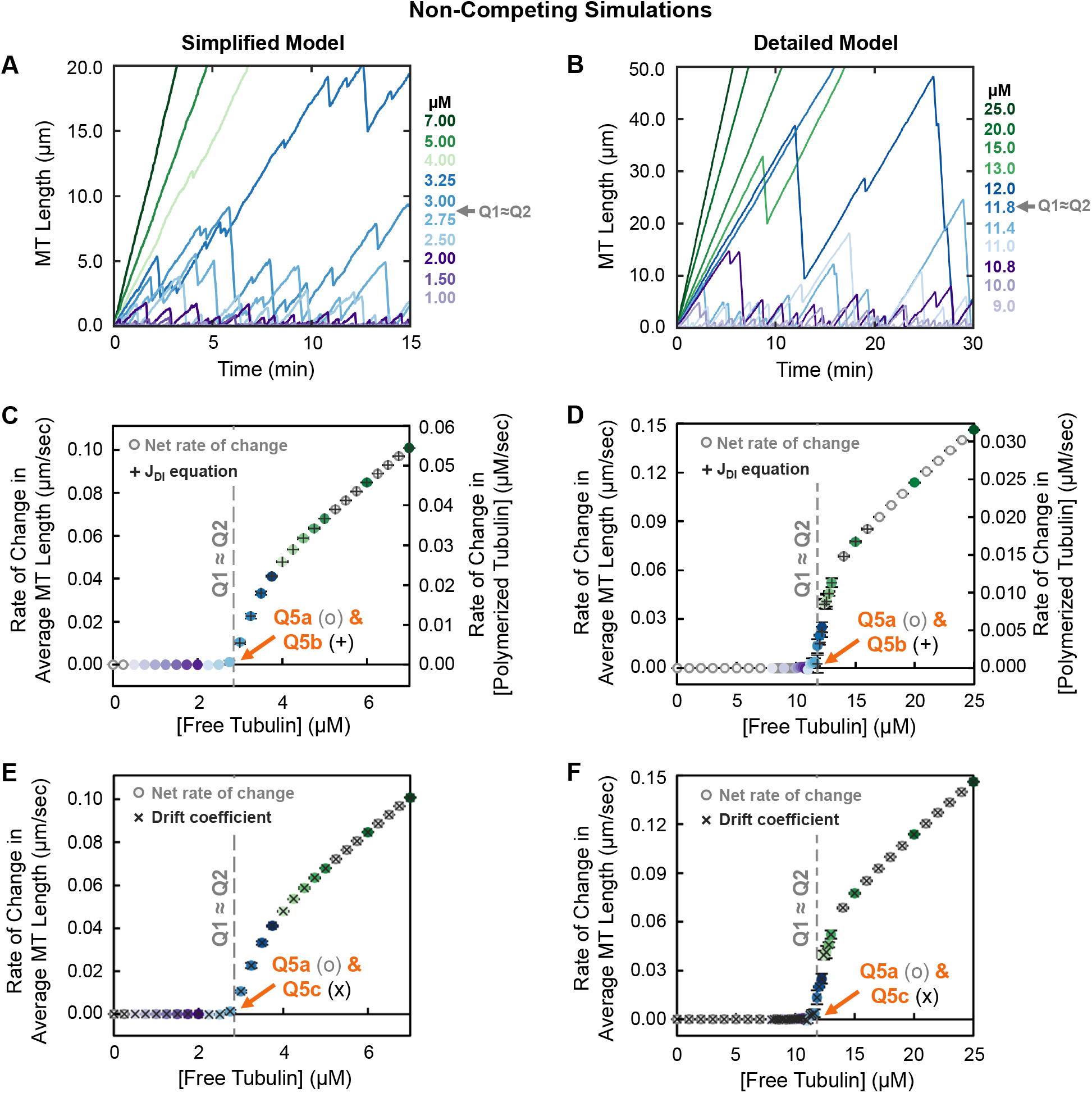
Behavior of microtubules (individuals and populations) under conditions of constant free tubulin. Left panels: simplified model; right panels: detailed model; colors of data points reflect the concentrations of *free* tubulin. ***(A,B)*** Representative length history plots for one individual MT at each indicated constant free tubulin concentration. ***(C,D)*** Steady-state net rate of change (o symbols) in average MT length (left axis) or in concentration of polymerized tubulin (right axis) for the free tubulin concentrations shown. **Q5a** indicates the concentration at which this rate becomes positive. This panel also shows the theoretical rate of change in average MT length (+ symbols) as calculated from the extracted DI measurements using the equation *J*_DI_ = (*V*_g_ *F*_res_ – |*V*_s_|*F*_cat_)/(*F*_cat_ + *F*_res_) in the [free tubulin] range where *J*_DI_ > 0 (Equation 1 in the “unbounded growth” regime) (Hill and Chen, 1984; Walker et al., 1988; Verde et al., 1992; Dogterom and Leibler, 1993). **Q5b** is the concentration at which J_D_| becomes positive. ***(E,F)*** Drift coefficient (Komarova et al., 2002) of MT populations as a function of [free tubulin] (x symbols). **Q5c** is the concentration above which drift is positive. For ease of comparison, the rate of change in average MT length (o symbols) from panels **C** and **D** is re-plotted in panels **E** and **F** respectively. For additional data related to these simulations, see **Figure S3**. ***Interpretation***: The results show that **Q5a ≈ Q5b ≈ Q5c**, hereafter referred to as **Q5**. At concentrations *below* **Q5**, populations of MTs reach a polymer-mass steady state where the average MT length is constant over time (the rate of change in average MT length or polymer mass is approximately zero; panels **C-D**), and the system of MTs exhibits zero drift (panels **E-F**). At free tubulin concentrations *above* **Q5**, populations of MTs reach a polymer-growth steady state where the average MT length and polymer mass increase over time at constant average rates that depend on [free tubulin] (panels C-D), and system of MTs exhibits positive drift (panels **E-F**). The average MT length as a function of time is shown in **Figure S3A-B**. Note that the concentration range below **Q5** corresponds to the “bounded” regime as discussed by Dogterom et al., while that above **Q5** corresponds to the “unbounded” regime (Dogterom and Leibler, 1993). The overall conclusions of the data in this figure are that (i) MTs exhibit net growth (as averaged over time or over individuals in a population) at [free tubulin] above the value **Q5** (**Q5a ≈ Q5b ≈ Q5c**); (ii) Q5 is similar to the value Q1 ≈ Q2 (grey dashed line) as determined in **Figure 3A-B**. Thus, Q1, Q2, and **Q5** all provide measurements of the same critical concentration, defined as CC_PopGrow_ in the main text. ***Methods***: All population data points (panels **C-F**) represent the mean +/− one standard deviation of the values obtained in three independent runs of the simulations. In panels **C-D**, the net rate of change was calculated from 15 to 30 minutes. In panels E-F, the drift coefficient was calculated using a method based on Komarova *et al.* (Komarova et al., 2002) (Supplemental Methods).

#### How MT behaviors relate to CC_PopGrow_

Examination of **Figure 5** shows that MT polymerization behavior in non-competing conditions (i.e., where [free tubulin] is constant) can be divided into two regimes:

**Polymer-mass steady state:** At concentrations of free tubulin below CC_PopGrow_ (measured by Q5a), both the average MT length and [polymerized tubulin] within a *population* reach steady-state values that increase with [free tubulin] but are constant with time (**Figures 5C-D**, **S3A-B**). *Individual* microtubules in these systems exhibit what might be called “typical” dynamic instability: they undergo periods of growth and shortening, but they eventually and repeatedly depolymerize back to the stable MT seed (**Figure 5A-B**).
**Polymer-growth steady state**: At CC_PopGrow_, the *populations* of dynamic MTs undergo a major change in behavior: they begin to grow persistently. More specifically, when [free tubulin] is above label Q5a in **Figure 5C-D**, there is no polymer-mass steady state where [polymerized tubulin] is constant over time (**Figure S3A-B**). Instead, the system of MTs arrives at a different type of steady state where [polymerized tubulin] increases at a constant rate (**Figures 5C-D, S3A-B**). *Individual* MTs within these populations still exhibit dynamic instability (except perhaps at very high [free tubulin]), but they exhibit unbounded growth (Dogterom and Leibler, 1993) (also described as net assembly (Walker et al., 1988)) if their behavior is assessed over sufficient time (**Figure 5A-B**).

Significantly, for both simulations, Q5a (**Figure 5C-D**) lies at approximately the value of Q1 ≈ Q2 (**Figures 3A-B**). This observation indicates that steady-state [free tubulin] in competing systems asymptotically approaches the same [free tubulin] at which microtubules begin to exhibit net growth (i.e., unbounded growth) in non-competing systems. In other words, these data show that CC_SubSoln_ ≈ CC_PopGrow_. This conclusion means that classical methods for measuring “the CC for polymer assembly” do not yield the CC at which individual DI polymers *appear* (as traditionally expected), but instead yield the CC at which populations of polymers grow *persistently*.

### Other experimental methods for measuring CC_PopGrow_

As noted above, Dogterom and colleagues previously predicted the existence of a critical free tubulin concentration CC_unbounded_, at which MTs will transition from exhibiting “bounded growth” to exhibiting “unbounded growth”^4^ (Dogterom and Leibler, 1993; Dogterom et al., 1995). This prediction was experimentally verified by Fygenson et al. (Fygenson et al., 1994).

Dogterom and colleagues (Verde et al., 1992; Dogterom and Leibler, 1993) used an equation for the rate of change in average MT length as a function of the DI parameters to characterize bounded and unbounded growth (note that J is a typical abbreviation for flux):

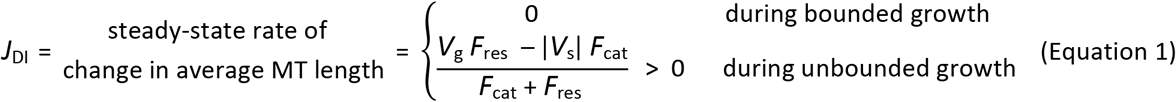

Dogterom et al. identified CC_unbounded_ as the [free tubulin] at which *V*_g_ *F*_res_ = | *V*_s_ | *F*_cat_ (see label Q5b in **Figure 5C-D**). Significantly, CC_unbounded_ as predicted by Q5b from this equation evaluated with our DI parameter measurements matches Q5a (compare + symbols to o symbols in **Figure 5C-D**). Hence, CC_PopGrow_ corresponds to CC_unbounded_, and polymer-mass steady state and polymer-growth steady state correspond to “bounded growth” and “unbounded growth” respectively. Thus, CC_PopGrow_ can be measured classically, by determining Q1 or Q2, but it can also be determined by measuring DI parameters for individual MTs at different [free tubulin] and inputting them into the equation for rate of change in average MT length (Equation 1). Having said this, determining DI parameters across a range of concentrations requires extended (> tens of minutes) analysis of many individual MTs and so is laborious and time consuming.

An alternative approach to measuring CC_PopGrow_ that may be more tractable experimentally is to use video microscopy to simultaneously analyze the behavior of many individual MTs within a population according to the drift paradigm of Borisy and colleagues (Vorobjev et al., 1997; Vorobjev et al., 1999; Komarova et al., 2002).^5^ The drift coefficient is the mean rate of change in position of the MT ends (for plus or minus ends separately), also described as the mean velocity of displacement of the MT ends. In cases where one end is fixed, as in our simulations, the drift coefficient is equivalent to the rate of change in average MT length. Here we used a method based on (Komarova et al., 2002), which calculates the drift coefficient from the displacements of MT ends over small time steps, e.g., between consecutive frames of a movie (see Supplemental Methods for additional information). As can be seen in **Figures 5E-F** (x symbols) and S3G-H (all symbols), a population of MTs at steady-state exhibits zero drift at [free tubulin] below Q5c (i.e., in this state, the average length of MTs in the population is constant with time) but exhibits positive drift at [free tubulin] above Q5c (i.e., the average MT length increases with time; the population grows persistently) (Komarova et al., 2002). As one might intuitively predict, Q5a ≈ Q5b ≈ Q5c (**Figure 5C-F**).

The evident similarity between the different measurements in **Figure 5C-F** suggests that Dogterom’s equation using DI parameters (+ symbols in **Figure 5C-D**) (Verde et al., 1992; Dogterom and Leibler, 1993) and the equation of Komarova et al. using short-term displacements (x symbols in **Figure 5E-F**) (Komarova et al., 2002) equations are simply two different representations of the same relationship. Indeed, both yield the rate of change in average MT length as functions of experimentally observed growth and depolymerization behaviors. Additionally, various forms of this equation were presented earlier by Hill and colleagues (Hill and Chen, 1984; Hill, 1987) and Walker et al. (Walker et al., 1988), and have since been used in other work (e.g., (Bicout, 1997; Gliksman et al., 1992; Maly, 2002; Vorobjev and Maly, 2008; Mourão et al., 2011; Mahrooghy et al., 2015; Aparna et al., 2017)). Thus, experimentalists should be able to use whichever analysis method is tractable and appropriate for their experimental system.

#### Measuring CC_PopGrow_ using population dilution experiments

Next we tested if CC_PopGrow_ is the same as the CC obtained from the population dilution experiments used in early studies of steady-state polymers (e.g., (Carlier et al., 1984b; Carlier et al., 1984a); see Q4 in **Table 1** and **Figure 1C**). These experiments measure the rate of change in [polymerized tubulin], which is also described the flux (typically abbreviated as J) of tubulin into or out of polymer. This measurement is performed after a population of MTs at steady state is diluted into a large pool of free tubulin at a new concentration.^6^ The measured data from the dilution experiments are then use to produce J(c) plots, where J is plotted as a function of subunit concentration “c” (**Figure 6A-B**). In these plots, “the CC” is identified as the dilution [free tubulin] at which J = 0 (i.e., where the plotted curve crosses the horizontal axis, Q4). At this concentration, individual MTs undergo periods of growth and shortening, but the population-level fluxes into and out of polymer are balanced (i.e., net growth is zero). We refer to the CC measured via J(c) plots as CC_flux_ (**Table 1**). CC_flux_ corresponds to one of the CCs that was identified by Hill and colleagues, variously named *c*_o_ in (Hill and Chen, 1984; Chen and Hill, 1985b) and *a*_α_ in (Hill, 1987).

**Figure 6:**
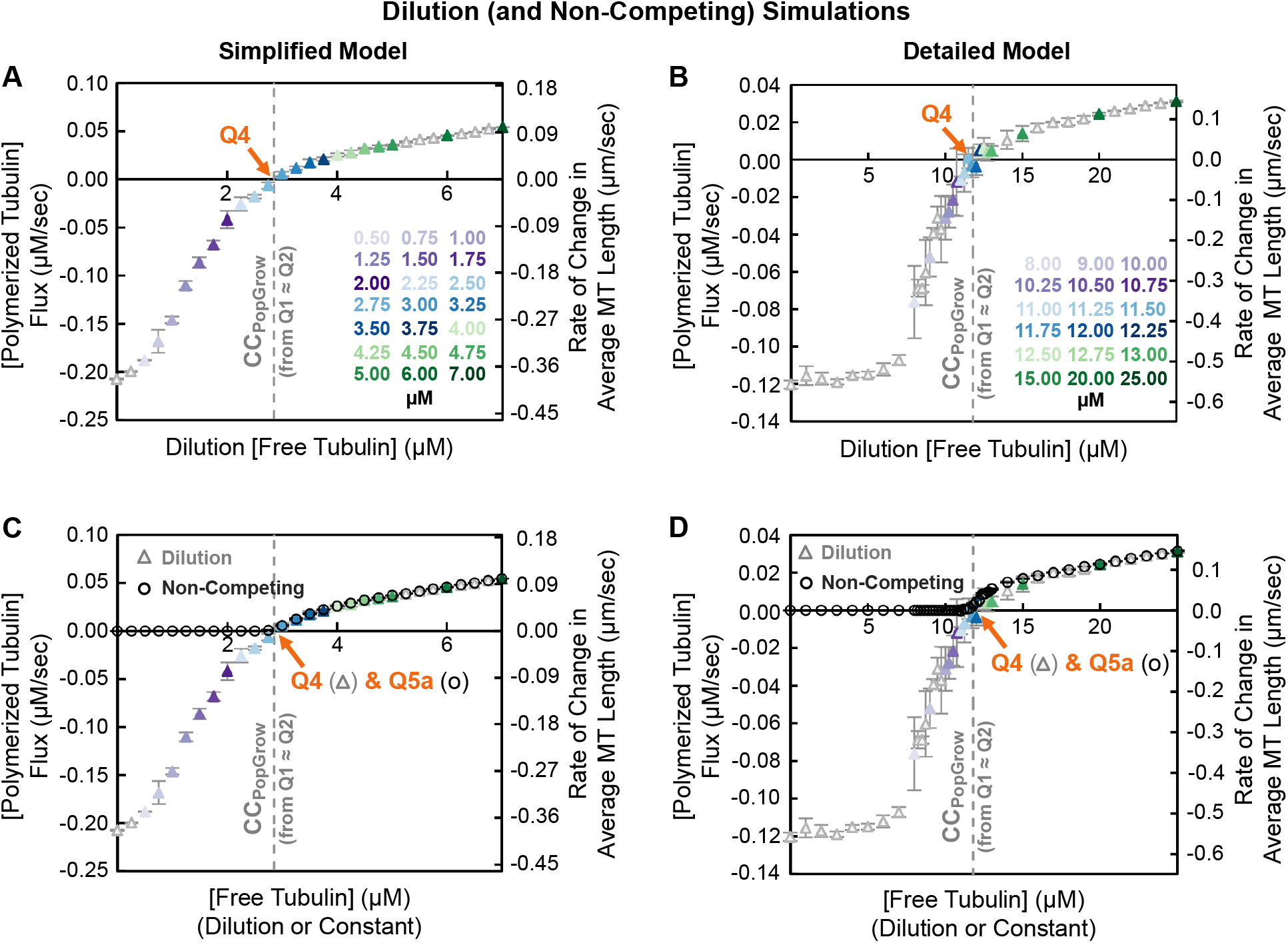
Flux of tubulin subunits into and out of MT polymer as a function of dilution [free tubulin]. (i.e., a J(c) plot as in (Carlier et al., 1984) and **Figure 1C**). Left panels: simplified model; right panels: detailed model. ***(A,B)*** In the dilution simulations, competing systems of MTs at high [total tubulin] were allowed to polymerize until they reached polymer-mass steady state. The MTs were then transferred into (“diluted into”) the free tubulin concentrations shown on the horizontal axis. After a brief delay, the initial flux (rate of change in [polymerized tubulin] (left axis) or in average MT length (right axis)) was measured, similar to (Carlier et al., 1984). ***(C,D)*** Data replotted to show that the J(c) curves from the dilution simulations in panels **A-B** (triangle symbols) and the net rate of change in average MT length from the constant [free tubulin] simulations in **Figure 5C-D** (circle symbols) overlay with each other for [free tubulin] above CC_PopGrow_ · ***Interpretation***: These data show that CC as determined by **Q4** from J(c) plots is approximately the same value as Q1 ≈ Q2 (grey dashed line), and thus **Q4** also provides a measurement of CC_PoPGrow_ ***Methods***: Competing systems of MTs at 22μM total tubulin were allowed to reach polymer-mass steady state. Then, at minute 10 of the simulation in the simplified model and at minute 20 of the simulation in the detailed model, the MTs were transferred into the free tubulin concentrations shown on the horizontal axis. After a 5 second delay, the flux was measured over a 10 second period (see **Figure S4** for plots of [free tubulin] and [polymerized tubulin] as functions of time). Note that the delay after dilution was necessary in the original experiments because of instrument dead time, but it is important for obtaining accurate J(c) measurements because it allows the cap length to respond to the new [free tubulin] (Duellberg et al., 2016; Bowne-Anderson et al., 2013). For accurate measurements, the pre-dilution MTs must be sufficiently long that none completely depolymerize during the 15-second period after the dilution. Data points for different concentrations of dilution [free tubulin] (see color key) represent the mean +/− one standard deviation of the values obtained in three independent runs of the simulations.

Significantly, the value of CC_flux_ as measured by Q4 in the dilution simulations corresponds to CC_PopGrow_ (grey dashed line, **Figure 6A-B**) as measured by Q1 ≈ Q2 in the competing simulations (**Figures 3A-B**) and by Q5abc in the non-competing simulations (**Figure 5C-F**). Note also that for dilution [free tubulin] above CC_PopGrow_, the J(c) curve obtained from the dilution simulations is superimposable with the net rate of change in average MT length obtained from the constant [free tubulin] simulations (**Figure 6C-D**). This observation might seem surprising given the differences in the experimental approaches; however, it makes sense because in each case the measurement is performed during a time period when [free tubulin] is constant and the rate of change has reached its steady-state value for each [free tubulin] (**Figures S3A-B** and **S4E-F**).

Thus, all of the experimental approaches for measuring critical concentration discussed thus far yield the critical concentration for persistent population growth (CC_PopGrow_). This conclusion leaves us with an unresolved question: What is the significance of the remaining common experimental CC measurement Q3 (obtained from experiments measuring growth velocity during growth phases for individual MTs as a function of [free tubulin], see **Figure 1B, Table 1**)?

### A critical concentration for transient elongation (growth) of individual filaments (CC_elongation_ = CC_IndGrow_)

Q3 (**Figure 1B**) has previously been used as a measure of the “critical concentration for elongation” (CC_elongation_) (Walker et al., 1988). According to standard models, CC_elongation_ is the free subunit concentration where the rate of subunit addition to an individual filament in the growth phase exactly matches the rate of subunit loss from that individual filament, meaning that individual filaments would be expected to grow at subunit concentrations above Q3 ≈ CC_elongation_ (see **Table 1** and its footnotes).

To determine the value of Q3 in our simulations, we used the standard approach for MTs as outlined in **Table 1** (experiments in (Walker et al., 1988); see also theory in (Hill and Chen, 1984; Hill, 1987)). We plotted the growth velocity (*V*_g_) of individual filaments observed during the growth phase of dynamic instability as a function of [free tubulin], and extrapolated a linear fit back to the [free tubulin] at which to *V*_g_ is zero.^7^ In addition to performing these measurements on the constant [free tubulin] simulations (**Figure 7A-B**), we also used the growth phases that occurred in the dilution experiments to obtain a measurement of CC_elongation_ (Q6 in **Figure 7C-D**). Comparing these measurements of CC_elongation_ in **Figure 7A-D** to the data in **Figures 3-6** shows that in both simulations CC_elongation_ (as determined by Q3 ≈ Q6) is well below CC_PopGrow_ as measured by any of the other approaches (Q1 ≈ Q2 ≈ Q4 ≈ Q5abc).^8^

**Figure 7:**
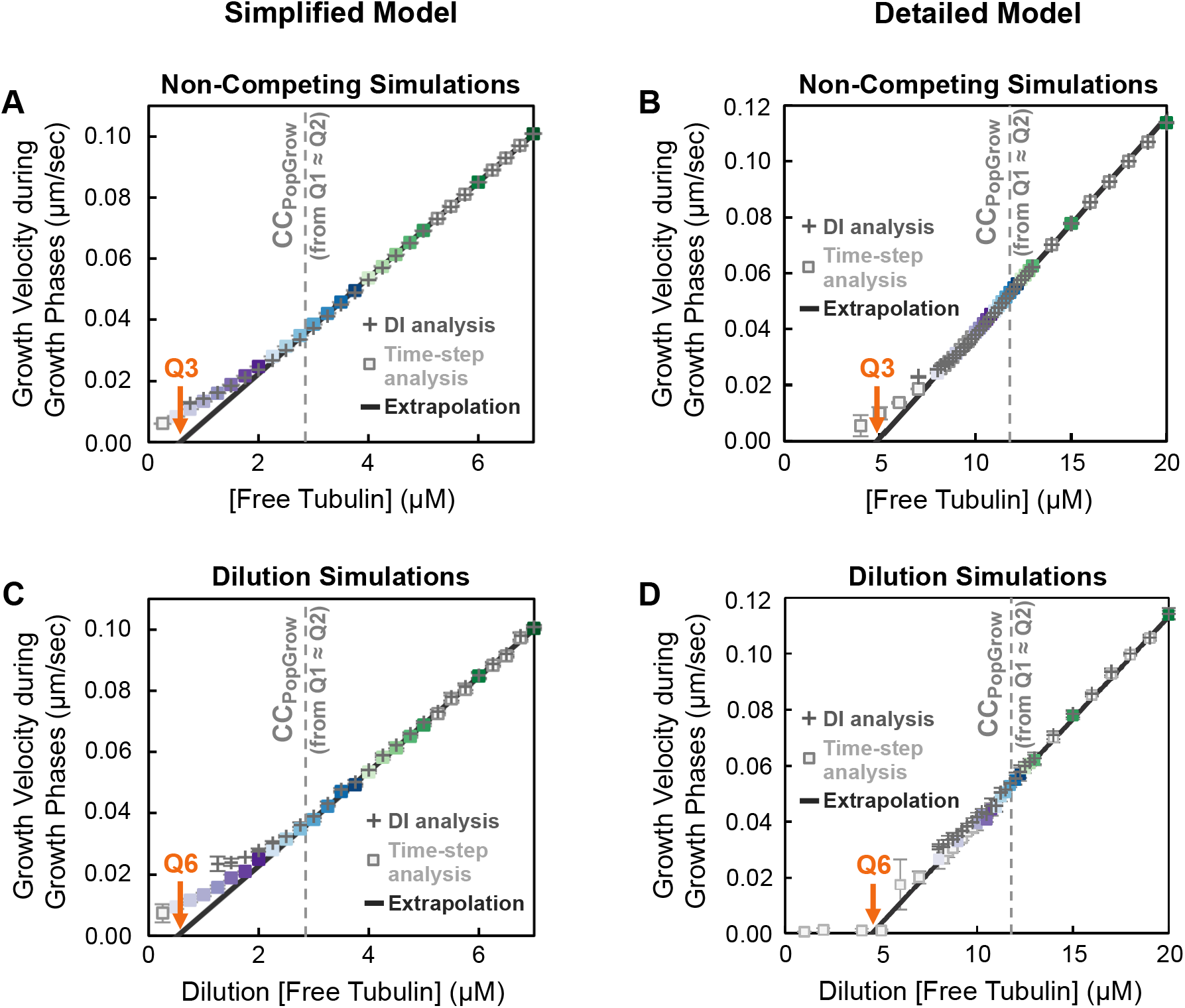
Growth velocity of individual MTs during the growth state as a function of [free tubulin]. Left panels: simplified model; right panels: detailed model; colors of data points reflect the concentrations of *free* tubulin. ***(A-D)*** Growth velocity (*V*_g_) measured using growth phases from either the constant [free tubulin] simulations of **Figure 5** (panels **7A-B**) or the dilution simulations of **Figure 6A-B** (panels **7C-D**). Each of panels **A-D** shows *V*_g_ as measured by a standard Dl-based analysis method (+ symbols) and a time-step based method (square symbols). Regression lines (solid black) were fitted to the linear range of these data, and extrapolated back to *V*_g_ = 0 to obtain **Q3** for the constant [free tubulin] simulations (panels **A-B**) and **Q6** for the dilution simulations (panels **C-D**). ***Interpretation***: These data show that the CC as measured by **Q3** is approximately equal to that measured by **Q6** and is different from CC_PopGrow_ (grey dashed line) as measured by Q1 ≈ Q2 (≈ Q4 ≈ Q5) from **Figures 3-6.** The main text provides justification for the idea that **Q3** and **Q6** estimate CC_IndGrow_, the CC for extended, but transient, growth phases of individual filaments. ***Methods***: In panels **A-B** (constant [free tubulin]), the *V*_g_ measurements were taken during the time period from minute 15 to minute 30 of the simulations (chosen so that the system has reached either polymer-mass or polymer-growth steady state). In panels **C-D** (dilution simulations), the *V*_g_ data were acquired from 5 to 15 seconds after the dilution, i.e., the J(c) measurement period described in **Figure 6**. For the Dl-based analyses (panels **A-D,** + symbols), we used a custom MATLAB program to identify and quantify growth phases by finding peaks in the length history data. The time-step based method (panels **A-D**, square symbols) divides each length history into 2-second intervals and identifies intervals during which there is a positive change in the MT length. See Supplemental Methods for more information about both methods. Regression lines were fitted to the time-step measurements of *V*_g_ for [free tubulin] in ranges where the *V*_g_ data are approximately linear as a function of [free tubulin]: from 3 to 7 μM for the simplified model (panels **A,C**) and from 7 to 15 μM for the detailed model (panels **B,D**). All data points represent the mean +/− one standard deviation of the values obtained in three independent runs of the simulations.

This observation demonstrates that Q3 ≈ Q6 provides information about MT behavior not provided by the other measurements. Specifically, since Q3 and Q6 are determined from measurements of the growth velocity of individuals during the growth phase of dynamic instability, Q3 and Q6 provide estimates of the [free tubulin] above which *individual* filaments *can* grow *transiently* (i.e., to extend during the growth phase of dynamic instability). Whether growth phases *will* occur also depends on a variety of other factors, such as rescue frequency and frequency of initiating growth from seeds. Consistent with the identification of the upper CC as CC_PopGrow_ for population growth, we suggest referring to CC_elongation_ as CC_IndGrow_ for individual filament growth.

Comparison of **Figure 7** with of **Figures 5-6** leads to additional conclusions with practical significance for measuring the CCs. **Figure 8A-B** shows that the *V*_g_ data from individual growth phases in **Figure 7A-B** and the net rate of change in average MT length data from populations in **Figure 5C-D** (or equivalently the population drift coefficient in **Figure 5E-F**) overlay each other when [free tubulin] is sufficiently high (i.e., far enough above CC_PopGrow_ that catastrophe is rare). This makes sense because when catastrophe is unlikely, almost all MTs will be growing, so measurements of individuals and populations should give approximately the same results. Thus, linear extrapolation from the net rate of change in average MT length data at high [free tubulin] to obtain Q7 as shown in **Figure 8C-D** yields approximately the same value for CC_IndGrow_ as Q3 ≈ Q6. Additionally, since the net rate of change in average MT length data from the constant [free tubulin] experiments and J(c) from the dilution experiments match each other at high [free tubulin] (**Figure 6C-D**), the Q7 extrapolation can also be performed on the J(c) data to measure CC_IndGrow_. Thus, both constant [free tubulin] experiments and dilution experiments can be used to obtain not only CC_PopGrow_ (via Q4 ≈ Q5abc) but also CC_IndGrow_ (via Q3 ≈ Q6 ≈ Q7).

**Figure 8:**
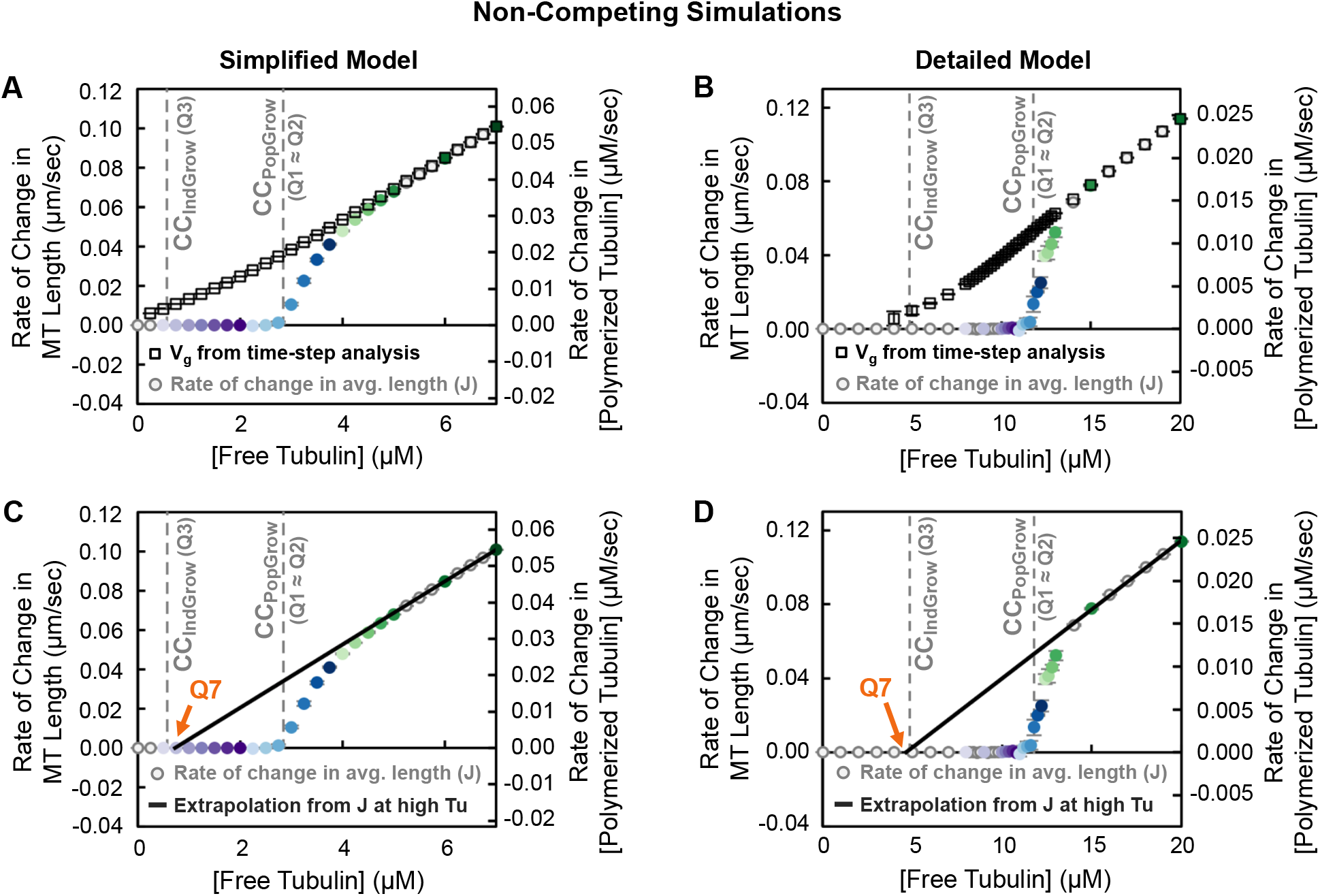
An alternative method for measuring CC_IndQrow_. Left panels: simplified model; right panels: detailed model. In all panels, the grey dashed lines represent CC_IndGrow_ as measured by Q3 (**Figure 7A-B**) and CC_PopGrow_ as measured by Q1 ≈ Q2 (**Figure 3A-B**). ***(A,B)*** Overlay of *V*_g_ from the time-step analysis of growing individual MTs (square symbols; re-plotted from **Figure 7A-B**) and the net rate of change in average MT length of the MT population (circle symbols; re-plotted from **Figure 5C-D**), both from the constant [free tubulin] simulations. ***Interpretation of panels A,B***: These data show that at high [free tubulin], the net rate of change in average MT length approaches the *V*_g_ of individual MTs. These two data sets converge because at sufficiently high [free tubulin] individual MTs are growing (nearly) all the time, as seen in the length histories (**Figure 5A-B**). Thus, CC_IndGrow_ which was obtained from *V*_g_ in **Figure 7**, should also be obtainable by extrapolating from the net rate of change data. ***(C,D)*** Extrapolation to obtain Q7 from the net rate of change in average MT length. ***Interpretation of panels C,D***: In each of the models, the value of **Q7** is approximately equal to Q3 ≈ Q6 (**Figure 7**). Thus, **Q7** provides another way of an estimating of CC_IndGrow_. ***Methods***: Regression lines were fitted to the net rate of change in average MT length for [free tubulin] in ranges where the net rate of change data is approximately linear as a function of [free tubulin]: from 6 to 7 μM for the simplified model (panel **C**) and from 14 to 20 μM for the detailed model (panel **D**). **Q7** is the x-intercept of the regression line. Note that the [free tubulin] ranges used for determination of Q7 are higher than those used for **Q3** and **Q6** because the **Q7** extrapolation requires conditions where all MTs in the population are growing (no depolymerization phases).

### CC_IndGrow_ is not CC_PolAssem_

The information above leads to the straightforward conclusion that CC_IndGrow_ represents a lower limit for microtubule growth. One might be tempted to use this idea to predict that CC_IndGrow_ is the concentration of free tubulin at which polymer appears (i.e., that CC_IndGrow_ = CC_PolAssem_). However, this prediction fails. Contrary to traditional expectation, there is *no total or free tubulin concentration at which polymer assembly commences abruptly*. Instead, the amount of polymer initially increases in a slow and nonlinear way with respect to [free tubulin], increasing more rapidly only as [free tubulin] approaches CC_PopGrow_ (**Figure S3A-F**). The same conclusion is reached whether examining polymer mass (**Figure S3A-B**), average MT length (**Figure S3A-F**), or maximal MT length (**Figure S3C-F**).^9^

These observations indicate that microtubules (and DI polymers more broadly) do not have a critical concentration for polymer appearance (CC_PolAssem_) as traditionally understood. CC_IndGrow_ is the tubulin concentration above which extended growth phases *can* occur, but significant amounts of polymer do not accumulate in experiments with bulk polymer until [free tubulin] nears or exceeds CC_PopGrow_ (**Figures 3, S1-S3**).

These behaviors might seem counterintuitive, but they can be explained by the following reasoning. First, when [free tubulin] is just above CC_IndGrow_, the growth velocity during the growth phase is low (*V*_g_ = 0 at Q3) and the frequency of catastrophe (*F*_cat_) is high. Under these conditions, individual microtubules will be both short (**Figure 5A-B, S3A-F**) and short-lived (**Figure 5A-B**), and thus difficult to detect. As [free tubulin] rises, MTs will experience growth phases that last longer (because *F*_cat_ drops) and also have faster growth velocity (**Figure 7**). The combined impact of these two effects creates a nonlinear relationship between [free tubulin] and [polymerized tubulin] or equivalently the average MT length observed at steady state; it similarly creates a nonlinear relationship between [free tubulin] and maximal MT length as observed within a period of time (**Figure S3C-F**).

### Measurement of CC_IndGrow_ by Q3, Q6, or Q7 is approximate

CC_IndGrow_ and CC_PopGrow_ are intrinsic properties of a system (i.e., a particular protein sequence in a particular environment), whereas the experimental measurements (Q values) are subject to measurement errors and are therefore approximate. The measurements of CC_IndGrow_ by Q3, Q6, or Q7 can be particularly sensitive to measurement errors and noise because they are based on extrapolations.

More specifically, since Q3 and Q6 are determined by extrapolations from regression lines fitted to plots of *V*_g_ versus [free tubulin], small changes in the *V*_g_ data (e.g., from noise) can be amplified when extrapolating to the *V*_g_ = 0 intercept. Additionally, in the simulation results, nonlinearities are observed in the *V*_g_ versus [free tubulin] plots in both simulations. In the presence of noise and/or nonlinearities, the values of Q3 and Q6 will depend on the [free tubulin] range where the regression lines are fitted to the *V*_g_ plots.

The deviation from linearity in the simulation plots is explained in part by measurement bias: at the lowest [free tubulin], there are few growing MTs, all of which are short (**Figures 5A-B, S3C-F**). The measured *V*_g_ data is biased towards those MTs that happened to grow fast enough and long enough to be detected (see maximum MT length data in **Figure S3C-F**). This indicates that the lowest concentrations should not be used in the linear extrapolation to identify Q3 or Q6. To our knowledge, such deviations from linearity at low concentrations have not been detected experimentally. However, the simulations generate much more data and at smaller length thresholds than is possible with typical experiments. Because measurement bias could also be a problem in physical systems, we speculate that similar effects may eventually be seen experimentally.

Given the nonlinearities and the measurement bias described above, one might be concerned that detection thresholds would affect the measured value of CC_IndGrow_. We therefore compared two different analysis methods for determining *V*_g_ (**Figure 7**). Specifically, for the DI analysis method (**Figure 7**, + symbols), we set a threshold of 25 subunits (200 nm) of length change to detect growth or shortening phases (we set this threshold to be comparable to typical length detection limits in light microscopy experiments). In contrast, for the time-step method (**Figure 7**, square symbols), we did not impose a threshold on the length change during each time step (see Supplemental Methods). The *V*_g_ results from the two methods agree well with each other in the [free tubulin] range used to determine CC_IndGrow_ (i.e., the range where *V*_g_ is approximately linear). Thus, when implementing *V*_g_ analysis to estimate CC_IndGrow_, the regression lines should be fitted to the linear region to avoid the effect of detection thresholds. If the regression lines are not fitted in the tubulin range where V_g_ is linear, then Q3 and Q6 will be less accurate approximations of CC_IndGrow_.

Depending on the specific system, Q7 may be a less accurate approximation than Q3 or Q6. Q7 is obtained from the rate of change in average MT length at free tubulin concentrations that are sufficiently high that (almost) all MTs are growing (i.e., where *V*_g_ and the rate of change in average MT length overlap with each other, **Figure 8**). Since the Q7 extrapolation is performed from higher concentrations than the Q3 or Q6 extrapolations, measurement errors or noise in the data can be further amplified. Moreover, *V*_g_ and the rate of change in average MT length may not overlap until tubulin concentrations are so high that experimental measurements may no longer be feasible (e.g., because of problems such as free nucleation).

In both the detailed model and in physical MTs, an additional factor can cause *V*_g_ to have nonlinearities as a function of [free tubulin] and therefore likely interferes with the accuracy of identifying CC_IndGrow_ via Q3, Q6, or Q7. Previous work has provided experimental and theoretical evidence that the GTP-tubulin detachment rate depends on the tubulin concentration (Gardner et al., 2011), contrary to the assumptions classically used to determine CC_elongation_. This observation has been explained by the occurrence of concentration-dependent changes in the MT tip structure (Coombes et al., 2013).

Thus, both detection issues and actual structural features can potentially make observed *V*_g_ measurements nonlinear with respect to [free tubulin]. As a result, the value obtained for CC_IndGrow_ from Q3 ≈ Q6 ≈ Q7 may depend on what [free tubulin] range is used for the linear fit. These observations mean that these values (Q3, Q6, Q7) provide at best approximate measurements of CC_IndGrow_.

### Effect of hydrolysis rate constant (*k*_H_) on CC_IndGrow_ and CC_popGrow_

The results above show that CC_IndGrow_ is obtained from measurements of individual MTs that are in the growth phase, while CC_PopGrow_ is obtained from measurements performed on populations (or on individuals over sufficient time) that include both growth and shortening phases (see also (Hill, 1987; Walker et al., 1988)). Thus, the co-existence of both growth and shortening phases (i.e., dynamic instability itself) occurs in conjunction with the separation between CC_IndGrow_ and CC_PopGrow_. Dynamic instability in turn depends on nucleotide hydrolysis, since GTP-tubulin is prone to polymerization and GDP-tubulin is prone to depolymerization. Therefore, to develop an improved understanding of the separation between CC_IndGrow_ and CC_PopGrow_ in DI polymers, we next examined the effect of the hydrolysis rate constant *k*_H_ on CC_IndGrow_ and CC_PopGrow_. To allow a straightforward comparison between the observed behaviors and the input kinetic parameters, we utilized the simplified model.

More specifically, we ran simulations in the simplified model under constant [free tubulin] conditions across a range of *k*_H_ values, while holding the other biochemical kinetic parameters constant. From these data we determined CC_IndGrow_ as measured by Q3 and CC_PopGrow_ as measured by Q5a (**Figures 9, S5**). When the hydrolysis rate constant *k*_H_ equals zero, only GTP-tubulin subunits are present. As would be expected, the behavior is that of an equilibrium polymer: no DI occurs (see length histories in **Figure S6A**), and all observed CC values correspond to the *K*_D_ for GTP-tubulin as defined by the input rate constants. In other words, when *k*_H_ is zero, CC_IndGrow_ = CC_KD_GTP_ = *k*_ToffT_/*k*_TonT_ = CC_PopGrow_ (**Figure 9A**). When *k*_H_ is greater than zero in these simulations, both GTP- and GDP-tubulin subunits contribute to polymer dynamics, concurrent with the appearance of dynamic instability (**Figure S6B-F**). As *k*_H_ increases, CC_IndGrow_ (Q3) and CC_PopGrow_ (Q5a) both increase and diverge from each other (**Figures 9, S5**), and dynamic instability occurs over a wider range of [free tubulin] (**Figure S6**).

**Figure 9:**
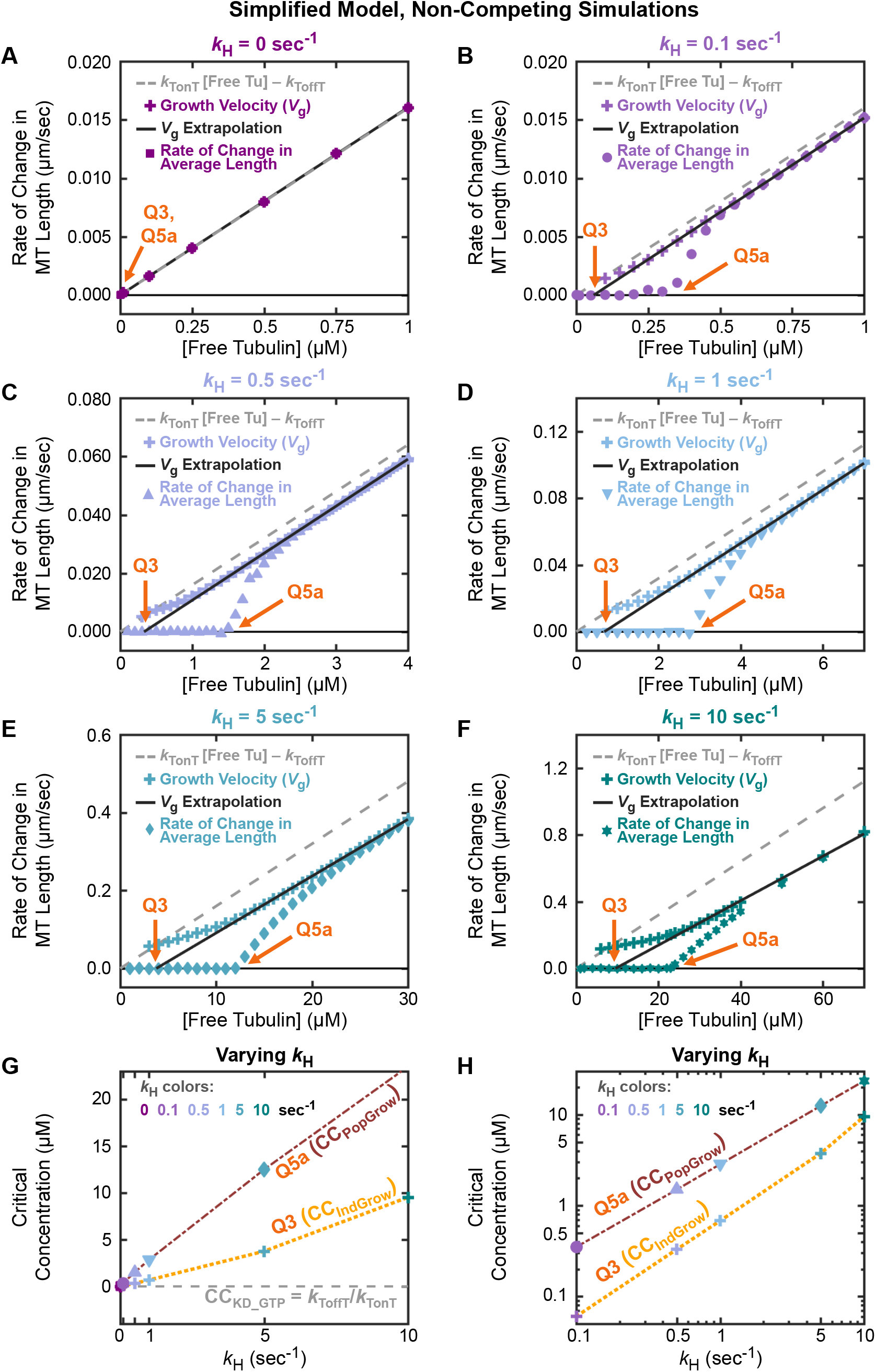
Effect of varying the rate constant for nucleotide hydrolysis (*k*_H_) in the simplified model. Non-competing simulations of the simplified model were performed for various values of *k*_H_ (all other input kinetic rate constants are the same as in the other figures). Each of panels **A-F** corresponds to a different value of *k*_H_, ranging from 0 to 10 sec^-1^, as indicated in the panel titles. ***(A-F)*** The growth velocity during growth phases (*V*_g_) (+ symbols; color coded by *k*_H_ value) and the rate of change in average MT length (color and symbol vary by *k*_H_ value) as functions of [free tubulin]. We also plot the theoretical equation for *V*_g_ that assumes that growing ends have only GTP-tubulin at the tips (grey dashed line). Note that the scales of the axes vary among panels **A-F**; for data replotted at the same scale, see **Figure S5**. ***(G,H)*** CC_IndGrow_ and CC_PopGrow_ as functions of *k*_H_, with CC_IndGrow_ and CC_PopGrow_ measured respectively by Q3 and Q5a from panels **A-F**. The axes have linear scales in panel **G** and log scales in panel **H**. The vertical separation between CC_IndGrow_ and CC_PopGrow_ at each *k*_H_ in the log-log plot (panel **H**) represents their ratio CC_PopGrow_/CC_IndGrow_. ***Interpretation***: When *k*_H_ is zero (panel **G**; see also panel **A**), C_IndGrow_ and CC_PopGrow_ are equal to each other and to CC_KD_GTP_. As *k*_H_ increases (panels **G** and **H**; see also panels **B-F**), the values of CC_IndGrow_ and CC_PopGrow_ increase, and diverge from each other and from CC_KD_GTP_. Thus, the separations between CC_KD_GTP_, CC_IndGrow_, and CC_PopGrow_ depend on *k*_H_. To see how DI behaviors relate to the CCs, see **Figure S6** for representative length history plots of individual MTs at each *k*_H_ value presented here. ***Methods***: The simulations were performed using the simplified model with 50 stable MT seeds. *V*_g_ was measured using the DI analysis method (Supplemental Methods). The steady-state rate of change in average MT length was measured from the *net* change method (see Q5a, **Table 3**). All measurements were taken from 40 to 60 minutes. Regression lines (black solid line) were fitted to the *V*_g_ data points in the [free tubulin] range above CC_PopGrow_ and then extrapolated back to *V*_g_ = 0.

### CC_IndGrow_ can differ from CC_KD_

In addition to showing that nucleotide hydrolysis drives CC_IndGrow_ (Q3) and CC_PopGrow_ (Q5a) apart from each other, the results in **Figure 9** also show that hydrolysis drives both away from CC_KD_GTP_ (x-intercept of grey dashed line in **Figure 9A-F**; grey dashed line in **Figure 9G**). In particular, while the relationship CC_KD_GTP_ = *k*_ToffT_/*k*_TonT_ is independent of *k*_H_, we observe that CC_IndGrow_ changes with *k*_H_. This could be viewed as surprising because one might expect CC_IndGrow_ to equal CC_KD_GTP_ even in the presence of DI. The reasoning behind this expectation is as follows.

First, the rate of growth of an individual MT during the growth state is assumed to change linearly with [free tubulin] according to the relationship,

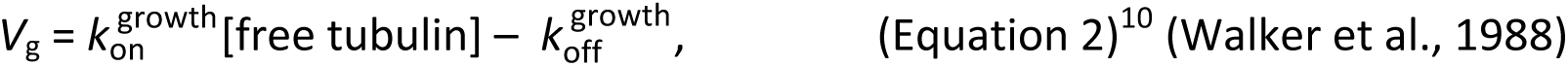

where 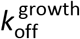 and 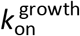 (called *k*_-1_^e+^ and *k*_2_^e+^ in (Walker et al., 1988)) are effective (observed) rate constants for addition and loss of GTP-tubulin subunits on a growing tip. By “effective” we mean that they are emergent quantities extracted from the *V*_g_ data, as opposed to directly measured kinetic rate constants. More specifically, the values of 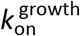 and 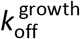 are measured from the slope and the y-intercept respectively of a regression line fitted to *V*_g_ data, given the equation above. Since CC_IndGrow_ is measured as the value of [free tubulin] at which *V*_g_ is zero, setting Equation 2 equal to zero and solving for [free tubulin] leads to the conclusion that 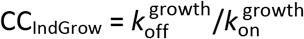. This ratio 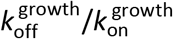 is measured as the x-intercept of the regression line (Equation 2) (Walker et al., 1988).

Second, it is commonly assumed that rapidly growing tips have only GTP-subunits at the end (e.g., (Howard, 2001; Bowne-Anderson et al., 2015)). Under this assumption, and also assuming that all unpolymerized tubulin is bound to GTP, Equation 2 becomes

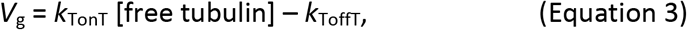

which leads to the prediction that CC_IndGrow_ = *k*_ToffT_/*k*_TonT_ = CC_KD_GTP_.

Instead, the results (**Figures 9A-F, S5A**) show that Equation 3 fits the data well only when *k*_H_ is close to zero. As *k*_H_ increases, the *V*_g_ regression line and CC_IndGrow_ diverge away from values that would be predicted from Equation 3.^11^ More specifically, when *k*_H_ is greater than zero, the effective 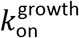 and 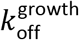 (slope and intercept of *V*_g_ in Equation 2) in the simulations diverge from *k*_TonT_ and *k*_ToffT_; in this case, *V*_g_ does not satisfy Equation 3, and CC_IndGrow_ diverges from CC_KD_GTP_. These observations indicate that GDP-subunits can influence behavior during growth phases.

#### Possible mechanisms for exposure of GDP-tubulin at growing MT tips

There are strong reasons to expect that GDP-subunits will influence growth behavior in physical MTs. The idea that growing MT tips could have GDP-tubulin subunits might seem surprising, but GDP-tubulin subunits could become exposed on the surface of a growing tip either by detachment of a surface GTP-subunit from a GDP-subunit below it or by direct hydrolysis. The first mechanism conflicts with earlier ideas that GTP-subunits rarely detach, but is consistent with recent experimental data indicating rapid exchange (attachment and detachment) of GTP-subunits on MT tips (Gardner et al., 2011; Coombes et al., 2013); see also (Margolin et al., 2012)).

The idea that GDP-tubulin cannot be exposed at MT tips during growth phases may be a remnant of vectorial hydrolysis models,^12^ where GDP-tubulin would become exposed only when the GTP cap is entirely lost (at least for single protofilament models). However, various authors have shown that vectorial hydrolysis is neither sufficient (Flyvbjerg et al., 1994; Flyvbjerg et al., 1996; Padinhateeri et al., 2012) nor necessary (Margolin et al., 2012; Padinhateeri et al., 2012) to explain MT dynamic instability behavior.

Additionally, Hill and colleagues examined both vectorial and random hydrolysis models. In the vectorial hydrolysis model, the growth velocity satisfied an equation equivalent to Equation 2 above, which assumes only GTP tips during growth (Hill, 1987). In their random hydrolysis model, the observed (emergent) slope and intercept of *V*_g_ did not equal the input rate constants for addition and loss of GTP-subunits, as explicitly pointed out in (Hill, 1987; Hill and Chen, 1984). This conclusion from Hill’s random hydrolysis model is consistent with the results of our model, which also has random hydrolysis.

The conclusion above that CC_IndGrow_ ≠ CC_KD_ also helps explain the observation from earlier in the paper that there is no concentration at which polymer assembly abruptly commences (i.e., there is no CC_PolAssem_). Instead, the amount of polymer increases slowly with increasing [free tubulin] (**Figure S3A-F**). More specifically, although the MTs typically reach experimentally detectable lengths (e.g., > 200 nm, depending on the method used) at some concentration above CC_IndGrow_ (**Figure S3A-F**), small amounts of growth can occur even below CC_IndGrow_ (**Figure S3E-F**; square symbols in **Figure 7**). When [free tubulin] is above CC_KD_GTP_, attachment to a GTP-subunit will be more favorable than detachment; thus, small amounts of growth can occur. In contrast, as noted above, CC_IndGrow_ is the [free tubulin] necessary for a microtubule to exhibit *extended* growth phases. The dependence of CC_IndGrow_ on *k*_H_ indicates that attachment must in some sense outweigh both detachment and hydrolysis of GTP-subunits in order for extended growth phases to occur.

### Dynamic instability can produce relationships previously interpreted as evidence of a nucleation process for growth from stable seeds

Previously, two experimental observations have been interpreted as evidence that growth of MTs from stable templates (e.g., centrosomes, axonemes, GMPCPP seeds) involves a nucleation process (e.g., conformational maturation or sheet closure) (Wieczorek et al., 2015; Roostalu and Surrey, 2017). First, MTs are generally not observed growing at [free tubulin] near CC_IndGrow_. Second, when the fraction of seeds occupied is plotted as function of [free tubulin], the shape of the resulting curve is sigmoidal, suggesting a cooperative process and/or a thermodynamic barrier. In this section we show that these two nucleation-associated behaviors are observed in our simulations, which is notable because neither simulation incorporates an explicit nucleation step (our seeds are composed of non-hydrolyzable GTP-tubulin). We show that both experimentally observed relationships can result from dynamic inability in combination with length detection thresholds. The behaviors of DI polymers relative to CC_IndGrow_ and CC_PopGrow_, as described above (e.g., **Figures 5A-B, S3A-F**), can therefore be helpful in understanding these relationships.

#### Failure to detect MT growth events in experiments at [free tubulin] near CC_IndGrow_ can result from physical detection limitations coupled with DI

As described above, when [free tubulin] is near CC_IndGrow_, *V*_g_ is small and *F*_cat_ is high, meaning that MTs are short (**Figure S3A-F**) and shortlived (**Figure 5A-B**); the average MT length remains small until [free tubulin] is closer to CC_PopGrow_ (**Figure S3A-F**). This behavior coupled with length detection thresholds (such as would be imposed by physical experiments) could make it difficult to detect MTs at [free tubulin] near CC_IndGrow_. To test this hypothesis, we used the simulations (which output the MT length without any detection threshold) to examine the effect of imposing length detection thresholds similar to those present in physical experiments.

Indeed, when we imposed a 200 nm detection threshold (comparable to light microscopy) on the length change needed for a growth phase to be recognized (**Figure 7**, + symbols), we saw that MT growth that was detected in the absence of this threshold (**Figure 7**, square symbols) is no longer detected. These results indicate that failure to observe MTs growing from stable seeds at [free tubulin] near CC_IndGrow_ can result from using experimental methods that have length detection limitations, providing evidence that such behavior can result from processes other than nucleation.

#### A sigmoidal P_occ_ curve is predictable from detection thresholds and MT population length distributions resulting from DI

*P*_occ_ is the proportion of stable MT templates/seeds that are occupied by a (detectable) MT (**Figure 10A-B**). Previous experimental work has shown that *P*_occ_ has a sigmoidal shape when plotted as a function of [free tubulin] (e.g., (Mitchison and Kirschner, 1984b; Walker et al., 1988; Wieczorek et al., 2015)). This shape has been interpreted as evidence that starting a new MT from a seed is harder than extending an existing MT and thus that growth from seeds involves a nucleation process (Walker et al., 1988; Fygenson et al., 1994; Wieczorek et al., 2015) (compare **Figure 11A** to **11B**). However, the *V*_g_ analysis described above led us to hypothesize that this sigmoidal *P*_occ_ shape can also result from the combination of length detection thresholds and DI.

**Figure 10:**
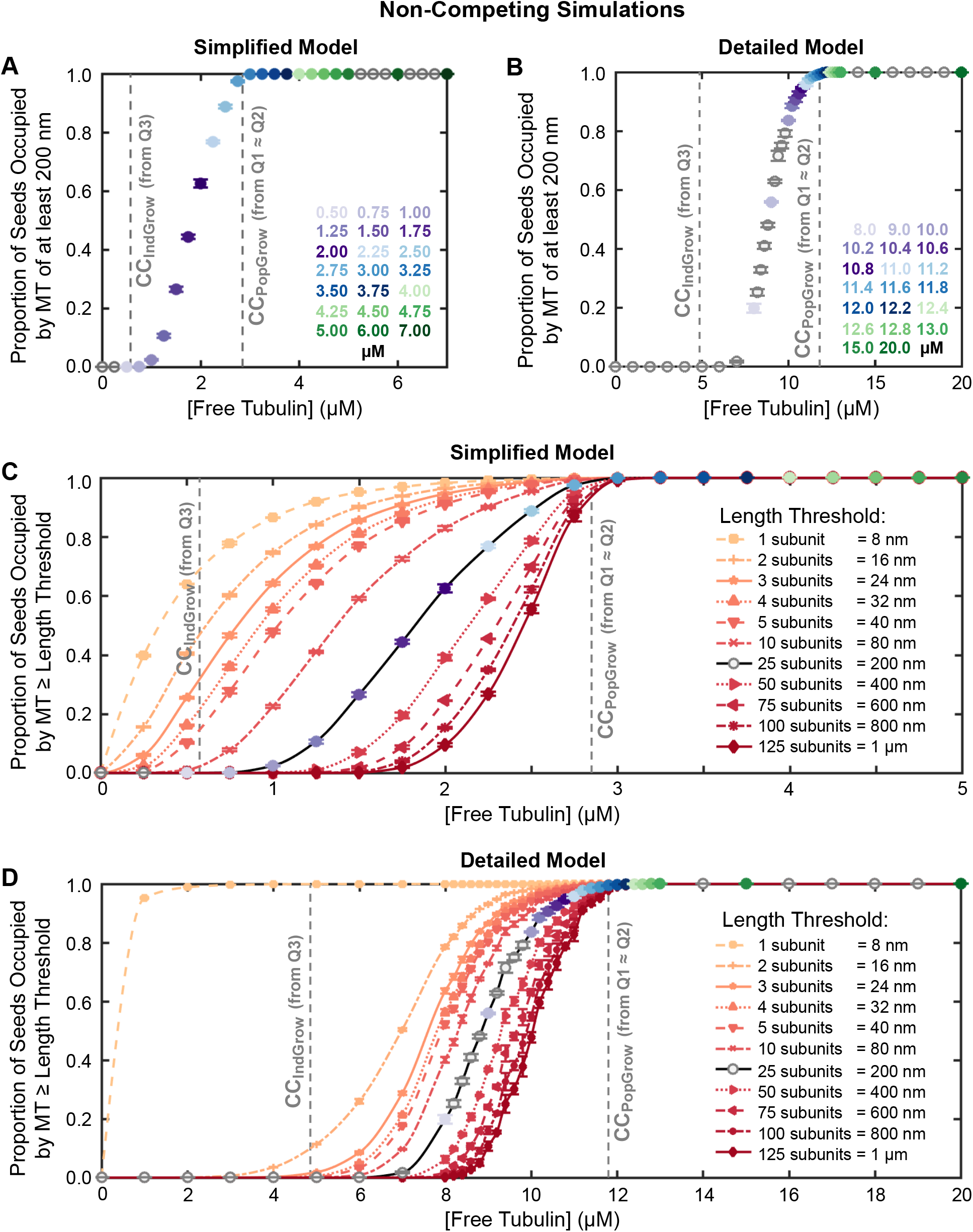
Relationship between *P*_occ_ (proportion of stable MT seeds that are occupied) and [free tubulin]. Simplified model in panels **A,C**; detailed model in panels **B,D**. The raw data analyzed in this figure are from the same non-competing (constant [free tubulin]) simulations used in **Figures 5, 6C-D, 7A-B**, and **8**. In all panels, the grey dashed lines represent CC_IndGrow_ (Q3 from **Figure 7A-B**) and CC_PopGrow_ (Q1, Q2 from **Figure 3A-B**). ***(A,B)*** Proportion of stable seeds bearing “experimentally-detectable” MTs (*P*_occ_) as a function of [free tubulin]. Here detectable MTs are those with length ≥ 25 subunits = ~200 nm (chosen because the Abbe diffraction limit for 540 nm (green) light in a 1.4 NA objective is ~200 nm). ***(C,D)*** *P*_occ_ with detection thresholds varied from 1 subunit (8 nm) to 125 subunits (1000 nm). The data with the 25 subunit threshold is re-plotted from panels **A-B**. ***Interpretation***: The data in panels A-B show that with a detection threshold similar to that in typical fluorescence microscopy experiments, little polymer is observed growing off of the GTP-tubulin seeds in either simulation until [free tubulin] is well above CC_IndGrow_. More specifically, with this 200 nm threshold, *P*_occ_ does not reach 0.5 until [free tubulin] is more than halfway from CC_IndGrow_ to CC_PopGrow_. Note that the lowest value of [free tubulin] at which 100 percent of the seeds have a detectable MT corresponds to ~CC_PopGrow_ (see also (Fygenson et al., 1994; Dogterom et al., 1995)). The data in panels C-D indicate that short MTs (with lengths below the 200 nm detection threshold from panels AB) are present at free tubulin concentrations near CC_IndGrow_. Additionally, we note that the *P*_occ_ curve of the detailed model is steeper than that of the simplified model when the same threshold is compared. We suggest that this results from the more cooperative nature of growth in the detailed (13-protofilament) model, which is an outcome of interactions between protofilaments. ***Methods***: All data points represent the mean +/− one standard deviation of the *P*_occ_ values obtained in three independent runs of the simulations. The values from each run are averages from 25 to 30 minutes, chosen so that *P*_occ_ has reached its steady-state value. MT length is measured as the number of subunits of above the seed. Note that in the detailed model, the MT length is the average of the 13 protofilament lengths and can therefore have non-integer values; see supplemental **Figure S7** for fractional thresholds below 2 subunits, which fill in the large gap between 1 and 2 subunits.

**Figure 11:**
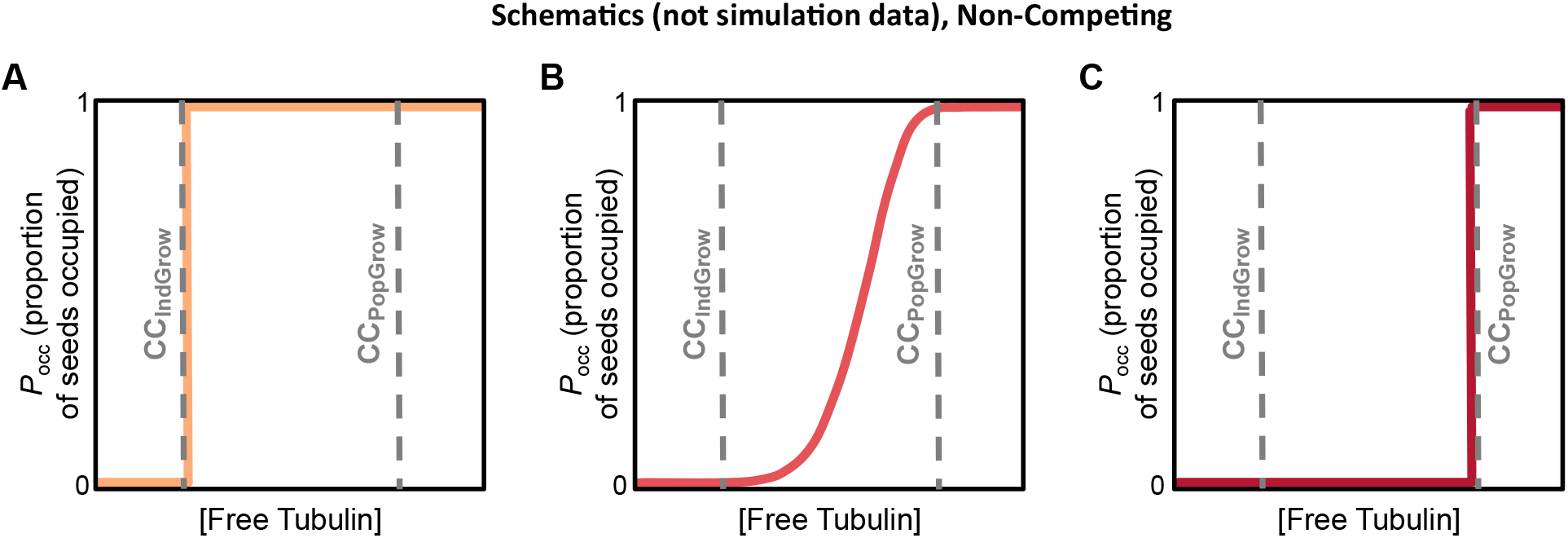
Hypothetical *P*_occ_ vs. [free tubulin] curves, where *P*_occ_ is the proportion of seeds that are occupied by MTs. It might have been expected that GTP-like seeds should start growing once [free tubulin] is above CC_IndGrow_, and that *P*_occ_ would therefore increase abruptly from 0 to 1 when [free tubulin] is at or just above CC_IndGrow_, similar to the step function in panel **A**. In contrast, sigmoidal *P*_occ_ curves, similar to panel **B**, have been observed experimentally (Mitchison and Kirschner, 1984b; Walker et al., 1988; Dogterom et al., 1995; Wieczorek et al., 2015). Obtaining a sigmoidal shape (**B**) instead of a step function (**A**) has been interpreted as evidence of a nucleation process that makes growth of MTs from stable seeds more difficult than extension from a growing end (Fygenson et al., 1994; Wieczorek et al., 2015). However, as discussed in the main text, this sigmoidal shape can be a consequence of DI in combination with experimental length detection limitations, and therefore is not necessarily evidence of a nucleation process. Note that a nucleation process that makes growth from seeds more difficult would lead to a *P*_occ_ curve that increases more rapidly from 0 to 1 and does so at [free tubulin] near CC_PopGrow_, similar to the step function in panel **C**. This behavior can be explained in the following way. When [free tubulin] is below CC_PopGrow_, MTs will repeatedly depolymerize back to the seed. When nucleation from seeds is difficult, it will take longer for a new growth phase to initiate after each complete depolymerization; seeds will therefore remain unoccupied for longer times and the proportion of seeds in the population that are occupied at any particular time be will lower. Thus, as the difficulty of nucleation increases, the shape of the *P*_occ_ curve would change from a sigmoid (as in panel **B**) to a step function at CC_PopGrow_ (as in panel **C**).

To test this hypothesis, we examined *P*_occ_ as a function of [free tubulin] with varying detection thresholds (**Figures 10C-D, S7**). The results show that at each [free tubulin] (below CC_PopGrow_), as the detection threshold is increased, the detected *P*_occ_ decreases (i.e., fewer MTs are longer than the threshold). This results in a sigmoidal shape emerging in the plots for both simulations as the length detection threshold is increased. The steepness of the sigmoid depends strongly on the detection threshold. These observations indicate that the sigmoidal shape can result simply from imposing a length detection threshold on a system (such as dynamic MTs) where some of the filaments are shorter than the detection threshold. In the presence of DI with complete depolymerizations back to the seeds (as occurs below CC_PopGrow_), MTs will necessarily be below any non-zero detection threshold for at least some amount of time.

#### The P_occ_ curve reaches 1 at [free tubulin] near CC_PopGrow_

The results in **Figures 10** and **S7** provide another observation relevant to understanding critical concentrations: in both simulations, *P*_occ_ approaches 1 as [free tubulin] approaches CC_PopGrow_ (except possibly at very small thresholds, where *P*_occ_ nears 1 at lower [free tubulin]). This result is predictable, with or without a nucleation process, because only at [free tubulin] above CC_PopGrow_ (where the population undergoes net growth) would all active seeds be occupied by MTs longer than an arbitrarily chosen length threshold. This full occupancy would occur if sufficient time is allowed, because at [free tubulin] above CC_PopGrow_, MTs will eventually become long enough to escape depolymerizing back to the seed. Thus, the idea that *P*_occ_ = 1 at [free tubulin] above CC_PopGrow_ after sufficient time may provide a practical way to identify CC_PopGrow_ experimentally (see also (Chen and Hill, 1985a; Fygenson et al., 1994; Dogterom et al., 1995)).

Taking all this information together, we propose that a combination of dynamic instability itself and the existence of detection thresholds contributes to phenomena (failure to observe growing MTs at [free tubulin] near CC_IndGrow_, **Figure 7**; and sigmoidal *P*_occ_ plots, **Figures 10, S7**) that have previously been interpreted as evidence that growth of MTs from stable seeds involves a nucleation process (Fygenson et al., 1994; Wieczorek et al., 2015). In fact, any process that makes growth from a seed more difficult than extension of a growing tip (i.e., a nucleation process such as sheet closure) would make the *P*_occ_ curve more step-like, not less so (**Figure 11**, compare panels **B** and **C**). While we cannot exclude the existence of nucleation processes such as conformational maturation or sheet closure in physical microtubules, our work suggests that neither sigmoidal *P*_occ_ curves nor absence of detectable MTs on seeds at [free tubulin] near CC_IndGrow_ are sufficient evidence to conclude that growth from templates (e.g., centrosomes, stable seeds) involves a physical nucleation process.

## DISCUSSION

### The behavior of MTs is governed by two major critical concentrations

Using the dynamic microtubules in our computational simulations, we examined the relationships between subunit concentration and polymer assembly behaviors for dynamic instability (DI) polymers. Our results show that there is no true CC_PolAssem_ as traditionally defined, meaning that there is no concentration where MTs abruptly come into existence. Instead, there are at least two major critical concentrations. There is a lower CC (CC_IndGrow_), above which *individual* filaments *can* grow *transiently*, and an upper CC (CC_PopGrow_), above which a *population* of filaments *will* grow *persistently* (**Figure 12B,D**). For [free tubulin] above CC_PopGrow_, individual MTs may still undergo dynamic instability (**Figure 12D**, blue length history), but will exhibit net growth over time (**Figure 12D**, blue and green length histories). What might be considered “typical” or “bounded” dynamic instability (where individual MTs repeatedly depolymerize back to the seeds) occurs at [free tubulin] between CC_IndGrow_ and CC_PopGrow_ (**Figure 12D**, purple length history; **Figure 12C**).

**Figure 12:**
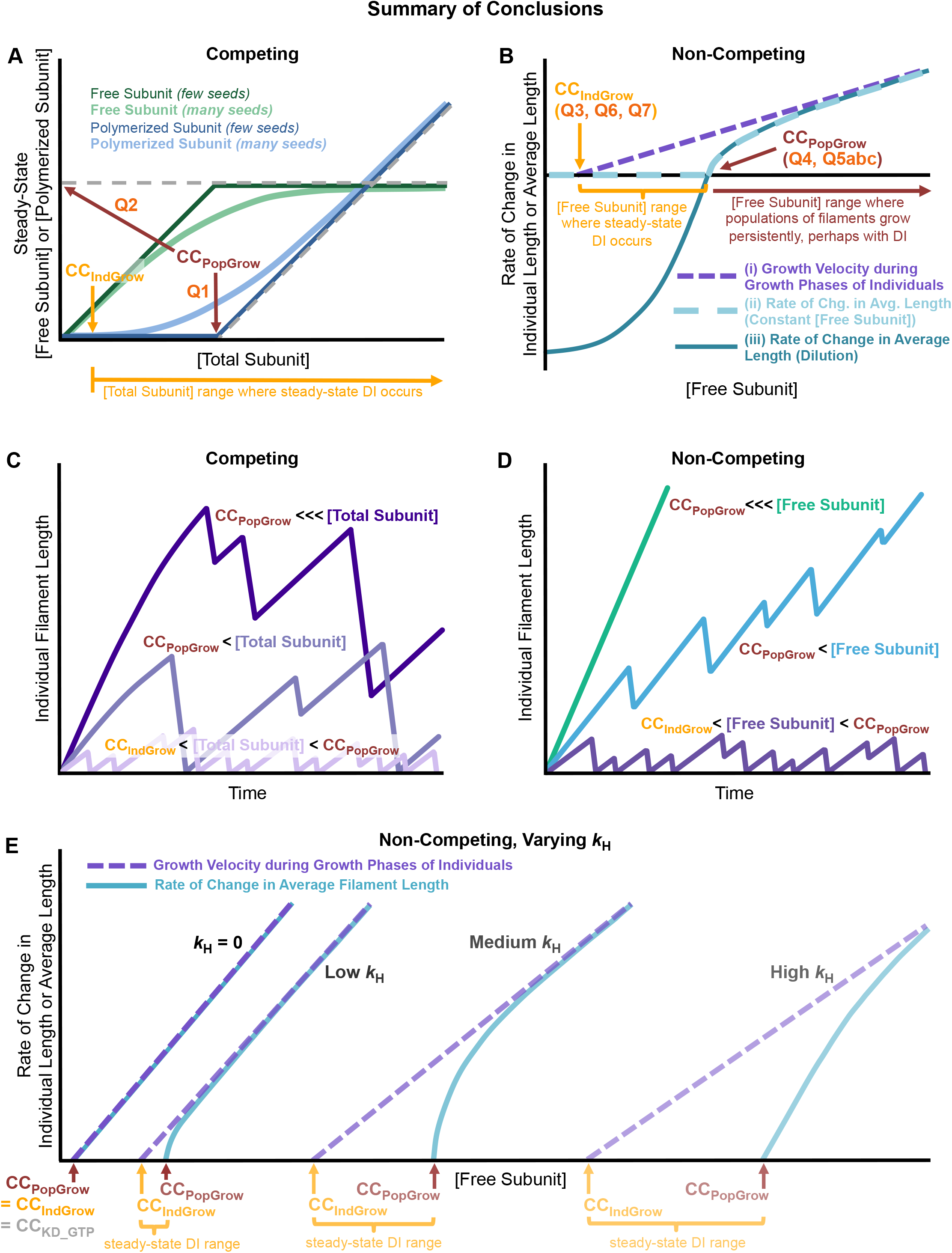
Schematic summary of the relationships between dynamic instability ( DI) behavior and critical concentrations for DI polymers. ***(A)*** Relationships between [total subunit] and [free subunit] (green) or [polymerized subunit] (blue) for a population of filaments competing for a fixed pool of subunits (constant [total subunit]) at polymer-mass steady state, similar to **Figures 3A-B, 4**). Notice that the steady-state [free subunit] in such competing systems approaches CC_PopGrow_ and that the sharpness of the approach depends on the number of seeds. In particular, for many seeds, steady-state [free subunit] is noticeably below CC_PopGrow_ even at very high [total subunit]. ***(B)*** Relationships between [free subunit] and the rate of polymerization/depolymerization under non-competing conditions (constant [free subunit]) for either individual filaments during growth phases or populations of filaments. More specifically, the panel shows: (i) the growth velocity (*V*_g_) of individual filaments during the growth phase (purple dashed line; similar to **Figure 7**); (ii) the net rate of change in average filament length in a population of filaments as assessed from experiments performed with [free subunit] held constant for the entire time of the experiment (light turquoise dashed curve; similar to **Figure 5C-F**); and (iii) the net rate of change in average filament length in a population of filaments as assessed from dilution experiments (dark turquoise solid curve; similar to **Figure 6**). Notice that curves (ii) and (iii) are superimposed for [free subunit] > CC_PopGrow_, and that curves (ii) and (iii) approach curve (i) for [free subunit] >>> CC_PopGrow_. ***(C-D)*** Length histories of individual filaments in competing systems (panel **C**) and non-competing systems (panel **D**). Note that when [free subunit] is below CC_PopGrow_ (purple length history in panel **D** and all length histories after polymer-mass steady state in panel **C**), individual filaments display steady-state dynamic instability in which they eventually and repeatedly depolymerize back to the seed; furthermore, the average filament length and the polymer mass reach finite steady-state values given sufficient time (see polymer-mass steady state in **Figures S1C-D** and **S3A-B**). When [free subunit] is above CC_PopGrow_ (panel **D**), individual filaments display net growth over time, while still undergoing dynamic instability (sky blue, panel **D**) except perhaps at very high concentrations (sea green, panel **D**). The text underneath the horizontal axis in panels **A-B** relates the dynamic instability behavior of individual filaments in panels **C-D** to the indicated ranges of [free subunit] and the population behaviors in panels **A-B**. ***(E)*** Effect of changing *k*_H_, the rate constant for nucleotide hydrolysis (similar to **Figure 9**). When *k*_H_ = 0 (equilibrium polymer), CC_KD_GTP_ = CC_IndGrow_ = CC_PopGrow_. When *k*_H_ > 0 (steady-state polymer), CC_IndGrow_ and CC_PopGrow_ are distinct from each other and from CC_KD_GTP_. As *k*_H_ increases, CC_IndGrow_ and CC_PopGrow_ both increase and the separation between them increases (at low enough *k*_H_, CC_IndGrow_ and CC_PopGrow_ would be experimentally indistinguishable); CC_KD_GTP_ does not change with *k*_H_. The [free subunit] range where steady-state DI occurs, i.e., the range between CC_PopGrow_ and CC_IndGrow_ (yellow brackets in panel **E**; compare to panels **B** and **D**), increases with *k*_H_. Note that this figure is a schematic representation of behaviors over a wide range of concentrations and is not drawn to scale.

CC_IndGrow_ is estimated by Q3, Q6, and Q7, and CC_PopGrow_ is estimated by Q1, Q2, Q4, and Q5abc (**Figure 12A-B, Table 3**). Classical critical concentration measurements (e.g., **Figure 1A** Q1 and Q2) do not yield the traditionally expected CC_PolAssem_, but instead yield CC_PopGrow_ (**Figure 12A** Q1 and Q2). Importantly, [free tubulin]_SteadyState_ in a competing system does not equal CC_PopGrow_, but approaches CC_PopGrow_ asymptotically as [total tubulin] increases and depends on the number of stable seeds (**Figure 12A**, compare dark and light green lines).

#### Bulk polymer experiments can create the illusion that CC_PopGow_ corresponds to CC_PolAssem_

The above conclusion that MTs grow transiently at [tubulin] between CC_IndGrow_ and CC_PopGrow_ might appear to conflict with experimental observations reporting that bulk polymer is detectable only above Q1 (**Figure 1A**, see e.g., (Johnson and Borisy, 1975; Mirigian et al., 2013)). As discussed above, Q1 provides a measure of CC_PopGrow_, but is traditionally expected to provide the critical concentration for polymer assembly, CC_PolAssem_. This apparent conflict between these observations and the conclusions above can be resolved by recognizing that the fraction of total subunits converted to polymer will be low until the free tubulin concentration nears CC_PopGrow_. Thus, for [total tubulin] < CC_PopGrow_, [free tubulin] will be approximately equal to [total tubulin] (**Figure 12A**, dark green line), unless there are many stable seeds (**Figure 12A**, light green line). In contrast, for [total tubulin] > CC_PopGrow_, all free tubulin in excess of CC_PopGrow_ will be converted from free to polymerized form if sufficient time is allowed (**Figure S1A-D**).^13^ This conversion will happen because the average MT filament will experience net growth until [free tubulin] falls below CC_PopGrow_ (**Figure 12C**, compare early in time to later in time). The outcome of these relationships is that in bulk polymer experiments, little if any MT polymer mass will be detected^14^ until the total tubulin concentration is above CC_PopGrow_ (**Figure 12A**, dark blue line), even though dynamic individual MT filaments can transiently exist at tubulin concentrations below CC_PopGrow_ (**Figure 3C-D**). Thus, the experimental quantities Q1 and Q2 may look like the expected critical concentration for polymer assembly (CC_PolAssem_), but they actually represent the critical concentration for persistent population growth (CC_PopGrow_).

#### P_occ_ plots can create the illusion that there is a [free tubulin] at which MT assembly commences abruptly, i.e., that CC_PolAssem_ exists

*P*_occ_ plots with length detection thresholds (such as thresholds intrinsic to microscope-based experiments) (**Figure 10A-B**) may have led to the conclusion that there is a CC_PolAssem_, at which *P*_occ_ first becomes positive. However, at low [free tubulin], MTs are short and short-lived as a result of low *V*_g_ and high *F*_cat_, as described above, and therefore can be undetectable by standard microscopy. By varying the length detection threshold imposed on simulation data (**Figure 10C-D**), it can be seen that the [free tubulin] at which *P*_occ_ first becomes positive depends on the threshold. These results, together with the polymer mass, average length, and maximal length data (**Figures S1C-F, S3A-F**) indicate that there is no concentration at which assembly of DI polymers commences abruptly.

#### Two additional CCs help define polymer behaviors

In addition to the major CCs (CC_IndGrow_ and CC_PopGrow_), there are at least two additional CCs that impact MT assembly. The first of these is CC_KD_GTP_ = *k*_ToffT_/*k*_TonT_, which corresponds to the *K_D_* for binding of a free GTP-tubulin subunit to a GTP-tubulin at a MT tip. The second additional CC is the *K*_D_ for binding of a free GDP-tubulin subunit to a GDP-tubulin at a MT tip, CC_KD_GDP_ = *k*_DoffD_/*k*_DonD_. Since CC_KD_GTP_ and CC_KD_GDP_ provide biochemical limits on the behaviors of GTP-tubulin and GDP-tubulin, any CCs must lie between these two nucleotide-specific CCs (CC_KD_GTP_ ≤ CC_IndGrow_ ≤ CC_PopGrow_ ≤ CC_KD_GDP_). CC_KD_GTP_ is the [free tubulin] above which GTP-tubulin polymers will grow in the absence of hydrolysis and provides the lower limit for short-term assembly of polymers in the presence of hydrolysis. As the hydrolysis rate constant increases, CC_IndGrow_ (the [free tubulin] above which extended growth phases can occur) can diverge from CC_KD_GTP_ (**Figure 9G**, compare yellow CC_IndGrow_ line to grey CC_KD_GTP_ line). Unlike CC_KD_GTP_, CC_KD___GDP_ is not straightforwardly measurable for MTs, because GDP-tubulin subunits alone do not polymerize into microtubules (Howard, 2001), but could be relevant to other steady-state polymers (e.g., actin).

#### Separation between the CCs is created by GTP hydrolysis

By running simulations in the simplified model at different *k*_H_ values, we show that increasing *k*_H_ causes CC_IndGrow_ and CC_PopGrow_ to diverge from each other and from CC_KD_GTP_ (**Figure 12E**). We expect that the magnitude of the separation between the various CCs will depend not on the value of *k*_H_ *per se*, nor on any individual rate constants, but rather on the relative relationships between the various rate constants. This is a topic of ongoing investigation. We speculate that the separation between the CCs has significance for understanding the difference between actin and tubulin, as discussed more below.

#### Relationship to previous work

As discussed above, the idea that MTs and other steady-state (energy-using) polymers have two major critical concentrations was first investigated in depth by Hill and colleagues, who studied the behavior of these systems using a combination of theory, computational simulations, and experiments (Hill and Chen, 1984; Carlier et al., 1984a; Hill, 1987). Their *c*_1_ (also referred to by other names including *a*_1_) corresponds to our CC_IndGrow_; their *c*_0_ (also called *a*_α_) corresponds to our CC_PopGrow_ (Hill and Chen, 1984; Hill, 1987). Moreover, Hill and Chen concluded that MTs grow at concentrations below what they referred to as the “real” CC (corresponding to our Q4 in **Figure 1C**) (Hill and Chen, 1984). However, the significance of this work for MT DI behavior was not fully incorporated into the CC literature, perhaps because it was not clear how their two CCs related to classical CC measurements (e.g., Q1 and Q2 in **Figure 1A**). Walker et al.’s seminal 1988 manuscript on dynamic instability parameters included measurements of two different critical concentrations that they termed the CC for elongation (CC_IndGrow_ in our notation) and the CC for net assembly (our CC_PopGrow_) (Walker et al., 1988). They calculated their value of the CC for net assembly from their measured DI parameters using a version of the J_DI_ equation (see equation on page 1445 of (Walker et al., 1988)). However, perhaps because the manuscript focused on CC_elongation_ and did not directly relate either of these CCs to those predicted by Hill and colleagues, the idea that MTs have two CCs still did not become widely acknowledged. Soon thereafter, the manuscripts of Dogterom et al. and Fygenson et al. were important in showing clearly and intuitively how the behavior of MTs changes at the CC for unbounded growth (our CC_PopGrow_), which they described using the J_DI_ equation shown in Equation 1 (Dogterom and Leibler, 1993; Fygenson et al., 1994; Dogterom et al., 1995). However, these authors did not relate their CC for unbounded growth to the CCs discussed by Hill or Walker et al. or to more classical CCs (**Table 1, Figure 1**).

Some of the continued confusion about critical concentration may have resulted from the fact that the published experimental work typically involved either competing conditions or non-competing conditions but not both. More specifically, classical experiments for determining “the critical concentration” (e.g., **Figure 1A**) involved competing conditions, but much of the previous work described above was performed under conditions of constant [free tubulin] (e.g., **Figure 1B-C**). Walker et al. (Walker et al., 1988) did note in their Discussion section that the concentration of free tubulin at steady state in their competing system was below their calculated CC for net assembly (i.e., CC_PopGrow_), contrary to the expectation that [free tubulin]_SteadyState_ would equal the CC for net assembly. They attributed this difference to “uncertainties inherent in [their] assumptions and measurements” (Walker et al., 1988). Instead, as shown above, the observation that [free tubulin]_SteadyState_ approaches CC_PopGrow_ without actually reaching it is a predictable aspect of dynamic instability. More specifically, [free tubulin]_SteadyState_ will be measurably below CC_PopGrow_ if [total tubulin] is not high enough relative to the value of CC_PopGrow_ and/or if the number of stable seeds is large (**Figures 3A-B, 4**).

More recently, Mourão et al. focused on systems of MTs growing under competing conditions (Mourão et al., 2011). Using stochastic simulations and mathematical analysis to study MT growth from stable seeds, they examined a quantity that they called “a baseline steady state free subunit concentration (*MD_SS_*)”, which is conceptually similar to our CC_SubSoln_ (measured by Q2). They concluded that [free tubulin]_SteadyState_ is not equal to *MD_SS_* but below it; our results are consistent with this conclusion. In particular, they demonstrated how the separation between [free tubulin]_SteadyState_ and *MD_SS_* depends on various factors including the number of stable MT seeds. The dependence of MT behavior on subunit concentration was not their primary focus, so they did not explicitly show that [free tubulin]_SteadyState_ asymptotically approaches *MD_SS_* = CC_PopGrow_ as [total tubulin] increases (**Figures 3A-B, 4**); however, they did perform simulations at three different values of [total tubulin] and their results are consistent with our conclusions. Additionally, the criterion that they used to determine the value of *MD_SS_* is that *MD_SS_* is the free tubulin concentration at which *V*_g_/|*V*_s_| = *F*_cat_/*F*_res_. We note that this equation is algebraically equivalent to |*V*_s_|*F*_cat_ = *V*_g_ *F*_res_, which was the criterion given by Dogterom et al. (Dogterom and Leibler, 1993) for identifying the CC for unbounded growth (equivalent to our CC_PopGrow_).

Thus, there has been a need for a unified understanding of how critical concentrations relate to each other and to MT behaviors at different scales. Our work fills this gap by clearly showing how the behaviors of individual MTs and populations of MTs relate to each other, to [free subunit] and [total subunit], and to a range of different experimental measurements in both competing and non-competing systems (conclusions summarized in **Figure 12** and **Table 3**). Taken together, our simulations and analyses should provide a more solid foundation for understanding the behavior of MTs and other DI polymers under varied concentrations and experimental conditions.

### Concurrence between different approaches for measuring MT behavior has practical significance

As shown in **Figure 5C-F**, there is remarkable concurrence between three seemingly disparate ways of measuring and analyzing MT behavior: (i) the net rate of change in [polymerized tubulin] (**Figure 5C-F**, o symbols), which is a bulk property obtained by assessing the mass of the population of polymers at different points in time (e.g., across 15 minutes); (ii) the J_DI_ equation (**Figure 5C-D**, + symbols), which uses DI parameters extracted from individual filament length history plots obtained over tens of minutes; (iii) the drift coefficient (**Figures 5E-F**, x symbols; **S3G-H**, all symbols) as measured from observing individual MTs in a population of MTs for short periods of time (e.g., 2-second time steps across as little as one minute). These approaches differ in attributes including physical scale, temporal scale, and experimental design. While the similarity of the data produced by these different approaches may initially be surprising, it can be shown that these measurements should yield the same values because the equations underlying them are also algebraically equivalent if certain assumptions are met (review article in preparation). In addition to yielding measurements of CC_PopGrow_ (Q5abc, **Figure 5**), these three experimental approaches can also provide approximate measurements of CC_IndGrow_ (Q7, **Figure 8**). The agreement between the results of these measurements indicates that the experimentally more tractable time-step approach (Komarova et al., 2002) (see Supplemental Methods) can be used to measure both CC_IndGrow_ and CC_PopGrow_ and should be used more frequently to quantitatively assess MT assembly behavior in the future.

### Biological significance of having two major critical concentrations

The understanding of critical concentration as presented above should help resolve apparently contradictory results in the microtubule literature. In particular, our results indicate that reported measurements of “the” critical concentration for MT polymerization vary at least in part because some experiments measure CC_IndGrow_ (e.g. (Walker et al., 1988; Wieczorek et al., 2015)), while others measure CC_PopGrow_ (e.g., (Carlier et al., 1984a; Dogterom et al., 1995; Mirigian et al., 2013)). This clarification should help in design and interpretation of experiments involving critical concentration, especially those investigating the effects of MT binding proteins (e.g. (Amayed et al., 2002; Wieczorek et al., 2015; Hussmann et al., 2016)), osmolytes (e.g, (Schummel et al., 2017)) or drugs (e.g. (Buey et al., 2005; Verma et al., 2016)).

Additionally, these ideas can be applied to help clarify the behavior of MTs *in vivo*. MTs in many interphase cell types grow persistently (perhaps with catastrophe and rescue, but with net positive drift) until they reach the cell edge, where they undergo repeated cycles of catastrophe and rescue with rare complete depolymerizations (Komarova et al., 2002). We showed previously that this persistent growth is a predictable outcome of having enough tubulin in a confined space: if sufficient tubulin is present, the MTs grow long enough to contact the cell boundary, which causes catastrophe; this drives the [free tubulin] above its natural steady-state value, which reduces catastrophe, enhances rescue, and induces the persistent growth behavior (Gregoretti et al., 2006). In light of the current results, we can now phrase this previous work more succinctly: persistent growth of MTs in interphase cells occurs when catastrophes induced by the cell boundary drive [free tubulin] above CC_PopGrow_. In contrast, at mitosis, when the MTs are more numerous and thus shorter, [free tubulin] remains below CC_PopGrow_. See also (Dogterom et al., 1995; Gregoretti et al., 2006; Vorobjev and Maly, 2008; Mourão et al., 2011) for relevant data and discussions.

Furthermore, it is important to emphasize that CC_IndGrow_ and CC_PopGrow_ are fundamental attributes of a specific type of tubulin in a particular environment, similar to the way a *K_D_* characterizes a protein-protein interaction or a *K*_M_ characterizes an enzyme-substrate reaction. Thus, we suggest using CC_IndGrow_ (as measured by Q3, Q6, or Q7) and CC_PopGrow_ (especially as measured by Q5c from the time-step drift coefficient approach) in addition to using dynamic instability parameters as a way to characterize tubulin (or other proteins that form polymers) and the activities of proteins that alter polymer assembly (see also the discussion in (Komarova et al., 2002)).

### Relevance for other steady-state polymers

Though the studies presented here were formulated specifically for MTs, we suggest that they can be applied to any nucleated, steady-state polymers that display dynamic instability, and perhaps to steady-state polymers more broadly. In particular, we propose that the key characteristic that distinguishes dynamically unstable steady-state polymers (e.g. mammalian MTs) from other steady-state polymers (e.g., mammalian actin) is as follows: for DI polymers, CC_IndGrow_ and CC_PopGrow_ are separable values driven apart by hydrolysis, but for other polymers, they are either identical (as is true for equilibrium polymers) or so close as to be nearly superimposed (e.g., mammalian actin). The values of CC_IndGrow_ and CC_PopGrow_ are emergent properties of the kinetic rate constants, which in turn are intrinsic properties of the protein sequence of the subunits (which comes from the gene sequence) and post-translation modifications (which come from the cell type and cellular signaling). Whether or not dynamic instability is physiologically relevant for a given polymer type in a specific cellular environment will depend on how the values of CC_IndGrow_ and CC_PopGrow_ relate to the cellular subunit concentration.

## METHODS

### Simulations

#### Simplified Model (Figure 2A)

As discussed in the main text, the simplified model of stochastic microtubule dynamics was described previously (Gregoretti et al., 2006), but the implementation used here was updated significantly. First, the code was rewritten in Java so that it could be more easily implemented on personal computers. Second, the time between events is now sampled using an exact version of the Gillespie algorithm (Gillespie, 1976), instead of an approximate version with a fixed time step. This change improves the accuracy with which the simulation carries out the underlying biochemical model with user-inputted rate constants. Third, the simulation was adjusted so that each simulated subunit now corresponds to an 8 nm MT ring (1 × 13 dimers) instead of a 20 nm MT brick (2.5 × 10 dimers) as in (Gregoretti et al., 2006). Also, the simulations in (Gregoretti et al., 2006) had a cell edge, which limited the MT lengths; the simulations presented here have no physical constraints on the MT lengths. The change in subunit size and the lack of physical boundary in the present simulation mean that the numerical values of the DI parameters and Q measurements (**Figures 3-8**, left panels) are not directly comparable between this implementation and our earlier publication (Gregoretti et al., 2006). However, the general behavior of the simulation is the same. The input parameters used here are as follows:

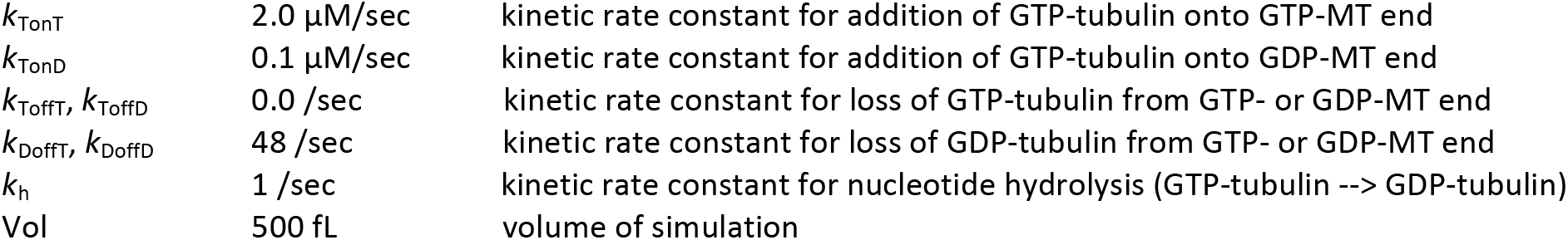

|  |  |  |
| --- | --- | --- |
| $k_{\text{TonT}}$ | 2.0 $\mu\text{M}/\text{sec}$ | kinetic rate constant for addition of GTP-tubulin onto GTP-MT end |
| $k_{\text{TonD}}$ | 0.1 $\mu\text{M}/\text{sec}$ | kinetic rate constant for addition of GTP-tubulin onto GDP-MT end |
| $k_{\text{ToffT}}, k_{\text{ToffD}}$ | 0.0 /sec | kinetic rate constant for loss of GTP-tubulin from GTP- or GDP-MT end |
| $k_{\text{DoffT}}, k_{\text{DoffD}}$ | 48 /sec | kinetic rate constant for loss of GDP-tubulin from GTP- or GDP-MT end |
| $k_h$ | 1 /sec | kinetic rate constant for nucleotide hydrolysis (GTP-tubulin → GDP-tubulin) |
| Vol | 500 fL | volume of simulation |

Unless otherwise indicated, each of the simplified model simulations was run with MTs growing from 100 stable seeds composed of non-hydrolyzable GTP-tubulin.

#### Detailed Model (Figure 2B)

The detailed model of stochastic microtubule dynamics with parameters tuned to approximate *in vitro* dynamic instability of mammalian brain MTs was first developed in (Margolin et al., 2011; Margolin et al., 2012) and later utilized in (Gupta et al., 2013; Li et al. 2014; Duan et al., 2017). The core simulation is the same as that in these prior publications, but this version has minor modifications including the addition of a dilution function to enable production of J(c) plots such as those in **Figure 6**. Please refer to (Margolin et al., 2012) for detailed information on the model, its parameter set C, and how its behavior compares to that of *in vitro* dynamic instability. Unless otherwise indicated, each of the detailed model simulations was run with MTs growing from 40 stable seeds composed of non-hydrolyzable GTP-tubulin in a volume of 500 fL.

The numerical values of the DI parameters for both models as measured by our automated DI analysis tool (described in the Supplemental Methods) are provided in the Supplemental Excel files. The values for the detailed model are similar to those that we published previously for this model (Margolin et al., 2012; Duan et al., 2017).

### Analysis

CC_PopGrow_ is estimated by determining Q1, Q2, Q4, or Q5 (**Figures 3-6**). CC_IndGrow_ is estimated (perhaps poorly) by determining Q3, Q6, or Q7 (**Figures 7-8**). See **Table 3B** for information on how to perform each of the Q measurements. The figure legends provide details about applying the measurements to the simulation data, including information about the time periods during which measurements were performed. The time periods were chosen to ensure that the variable being measured (e.g., rate of change in average length) has reached its steady-state value. For most of the measurements, this occurs when the simulated system has reached either polymer-*mass* steady state (non-competing systems with [free tubulin] < CC_PopGrow_, **Figure S3A-B**; and competing systems, **Figures S1A-D**) or polymer-growth steady state (non-competing systems with [free tubulin] > CC_PopGrow_, **Figure S3A-B**). In the Supplemental Methods, we describe the time-step analysis method (based on (Komarova et al., 2002)) used to measure drift and *V*_g_, as well as our DI analysis method used to measure *V*_g_, *V*_s_, *F*_cat_, and *F*_res_.

**Table 3A:** Revised understanding of critical concentration for dynamic instability polymers. Note that for steady-state polymers (including DI polymers), CC_KD_GTP_ ≤ CC_IndGrow_ ≤ CC_PopGrow_ ≤ CC_KD___GDP_, but for equilibrium polymers, CC_KD_ = CC_IndGrow_ = CC_Popgrow_.

| Critical Concentration | Representative Figures | Critical Concentration Description | Equivalent to (see Table 1)<sup>18</sup> | Measured by (see Table 3B) |
| --- | --- | --- | --- | --- |
| $CC_{PopGrow}$ | 1A,C, 3-6 | CC above which the polymer mass of a *population* will increase *persistently*, and individual filaments will undergo net growth over time | $CC_{SubSoln}$,<sup>19</sup><br>$CC_{flux}$,<sup>20</sup><br>$CC_{unbounded}$ | Q1, Q2, Q4, Q5 |
| $CC_{IndGrow}$ | 1B, 7, 8 | CC above which *individual* filaments *can* exhibit *transient*, but extended, growth phases | $CC_{elongation}$ | Q3, Q6, Q7 |
| $CC_{KD\_GTP}$ | 9 | Equilibrium dissociation constant for binding of a free GTP-subunit to a GTP-subunit at a polymer tip | | Any of the Q values above *under conditions where GTP is **not** hydrolyzed* |
| $CC_{KD\_GDP}$ | | Equilibrium dissociation constant for binding of a free GDP-subunit to a GDP-subunit at a polymer tip | | GDP-tubulin alone does not form MTs, so $CC_{KD\_GDP}$ is not straightforwardly measured |

<sup>18</sup> $CC_{PolAssem}$ is not listed here because there is no threshold concentration at which polymers abruptly appear. Instead, the measurement classically expected to yield $CC_{PolAssem}$ (see Q1 in Table 3B) actually yields $CC_{PopGrow}$ .
<sup>19</sup> Note that $CC_{SubSoln}$ is classically defined as the value of $[free\ tubulin]_{SteadyState}$ in a competing system whenever $[total\ tubulin]$ is above " $CC_{PolAssem}$ " (Table 1, Figure 1A). However, $CC_{SubSoln}$ is more accurately defined as the *asymptote approached* by $[free\ tubulin]_{SteadyState}$ as $[total\ tubulin]$ is increased (Q2 in Figures 3A-B, 4).
<sup>20</sup> It should be stressed that $CC_{flux}$ is the $[free\ tubulin]$ at which the *population-level* fluxes of tubulin into and out of polymer are balanced, while *individuals* may grow and shorten when $[free\ tubulin] = CC_{flux}$ .

**Table 3B:**
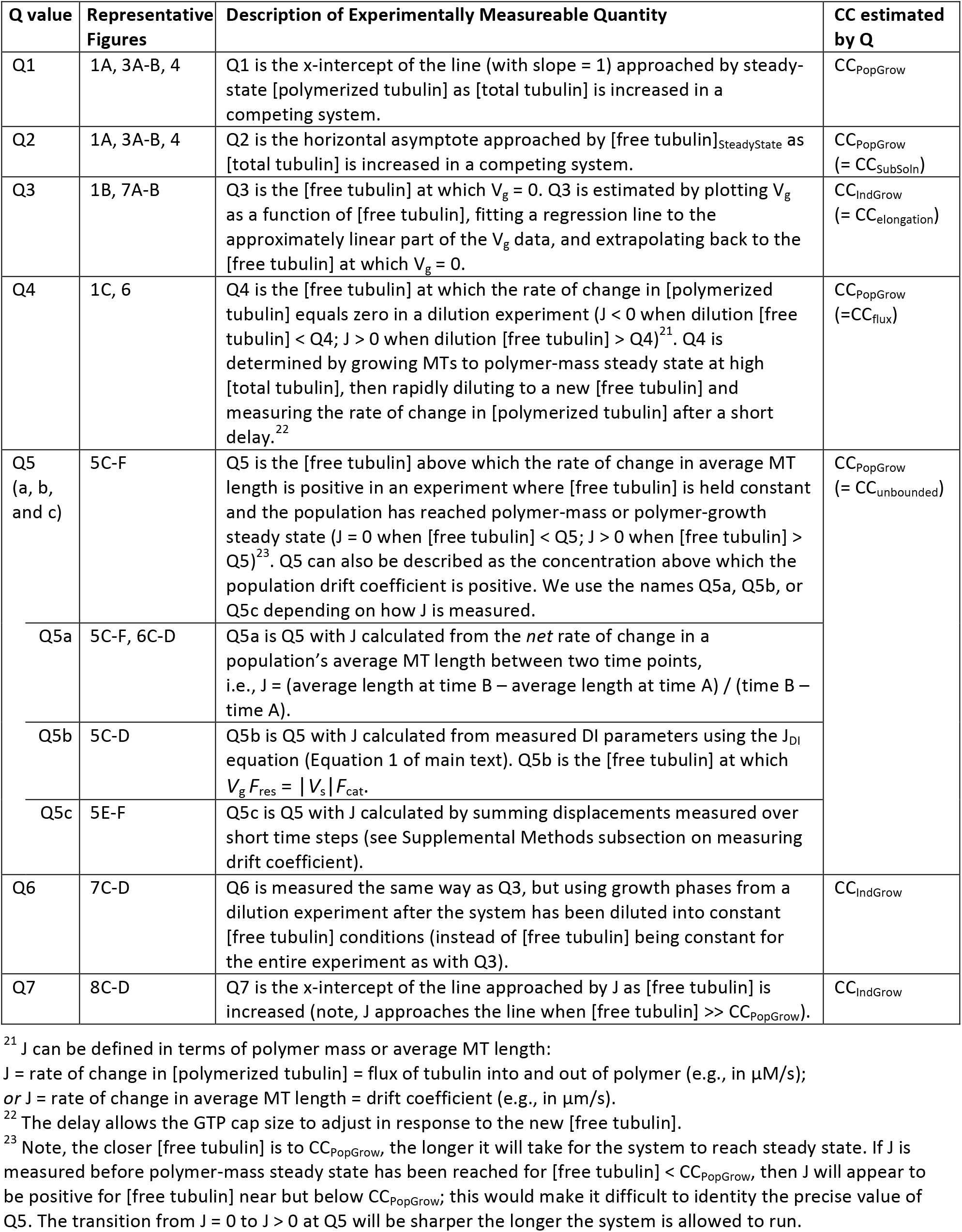
Summary of experimentally measureable quantities (Q values) used to estimate CCs. See **Table 3A** for descriptions of the CCs.

### Code Availability

The simulation codes (written in Java) and analysis codes (written in MATLAB) are available upon request.

## ACKNOWLEDGMENTS

This work was supported by NSF grants MCB-1244593 to HVG and MSA, MCB-1817966 to HVG, and MCB-1817632 to EMJ. Portions of the work were also supported by funding from the University of Massachusetts Amherst (AJM) and by a fellowship from the Dolores Zohrab Liebmann Fund (SMM). We thank the members of the Chicago Cytoskeleton community for their insightful discussions, and members of the Goodson laboratory for assistance in editing the manuscript.

## Footnotes

1 This manuscript focuses on systems composed of MTs with one end free and the other end anchored, such as would exist for MTs growing from centrosomes. In other cases, microtubules can have two free ends (plus and minus). For each of CC_IndGrow_ and CC_PopGrow_, the numerical value at the plus end could potentially differ from the value at the minus end.

2 Polymer-mass steady state describes a situation where the polymer mass has reached a plateau and no longer changes with time (other than small fluctuations around the steady-state value) (**Figure S1A-D**). Systems of dynamic microtubules can also have other steady states (e.g., polymer-length steady state)(see also (Mourão et al., 2011)).

3 Since the transitions are not sharp, it can be difficult to determine the exact values of Q1 and Q2. Depending on how the measurements are performed, the values of Q1 and Q2 might appear different from each other. However, Q1 = Q2 does hold if the measurements are performed as follows: Q2 is the value of the horizontal asymptote that [free tubulin]_SteadyState_ approaches as [total tubulin] increases; Q1 is the [polymerized tubulin] = 0 intercept of the line with slope 1 that [polymerized tubulin] approaches as [total tubulin] increases (**Figure 3A-B**). Note that Q1 would be exactly equal to Q2 in a system with no measurement error, no noise, and no non-functional tubulin, but for a physical experiment these factors can interfere with the measurements.

4 Note: Here a “bounded” system refers to one that has a constant steady-state polymer mass or average MT length; “unbounded” refers to a system where the polymer mass or average MT length exhibits net growth over time (Dogterom and Leibler, 1993; Dogterom et al., 1995). This situation should not be confused with one in which the system of MTs experiences a physical boundary (e.g., MTs in cells).

5 For a mathematical explanation of how microtubule behavior can be approximated by a drift-diffusion process, see (Maly, 2002; Vorobjev and Maly, 2008).

6 In the physical experiments, there was normally a delay of a few seconds after the dilution and before the data were recorded (Carlier et al., 1984a). Analysis of our simulated J(c) experiments incorporates a similar delay. This delay may have been necessary for technical reasons in the physical experiments, but it also serves a purpose in allowing the stabilizing GTP cap to redistribute to its steady-state size (Duellberg et al., 2016). See **Figure S4** for plots of [free tubulin] and [polymerized tubulin] as functions of time in the dilution simulations.

7 This *V*_g_ versus [free tubulin] relationship is expected to be linear on the basis of the assumption that growth occurs according to the equation *V*_g_ = *k*_TonT_ [free tubulin] – *k*_ToffT_, where the first term corresponds to the rate at which GTP-tubulin attaches to a GTP tip, and the second term corresponds to the rate at which GTP-tubulin detaches from a GTP tip (Bowne-Anderson et al., 2015). We return to this relationship later in the main text.

8 While the numerical values of the CCs in the simplified model should be regarded as arbitrary, the detailed model CC_IndGrow_ and CC_PopGrow_ numerical values closely match those obtained by Walker et al. (Walker et al., 1988). This is notable because we tuned the detailed model parameters to match Walker’s DI parameters at [free tubulin] = 10μM but did not tune to the CC values.

9 Note that Hill and Chen concluded that even equilibrium polymers have some assembly below the CC, but that the average length is very small until [free subunit] is “extremely close” to the CC (Hill and Chen, 1984).

10 The symbol = may be more appropriate than = because this equation assumes (i) that *V*_g_ increases linearly with [free tubulin] and (ii) that the detachment rate is independent of [free tubulin]. Our *V*_g_ results presented above indicate that assumption (i) may be inaccurate. See (Gardner et al., 2011) for evidence against assumption (ii).

11 Recall that the kinetic rate constants (e.g., *k*_TonT_, *k*_H_) in our simulations are inputted by the user. In contrast, the values of *V*_g_ and CC_IndGrow_ are emergent properties of the system.

12 In vectorial hydrolysis models, hydrolysis occurs only at the interface between the GDP-tubulin lattice and the GTP-tubulin cap (e.g., (Carlier et al., 1987; Hill, 1987)). In contrast, in random hydrolysis models, any internal GTP-subunit can hydrolyze (terminal GTP-subunits may also hydrolyze depending on the model).

13 More precisely, as indicated by the earlier discussion of **Figure 3A-B**, all subunits in excess of the steady-state [free tubulin] will be converted to polymer; the steady-state [free tubulin] is necessarily below but perhaps close to CC_PopGrow_.

14 The amount of polymer *present* depends on the kinetic rate constants of the particular system and the number of stable seeds (**Figure 4**). The amount of polymer *detected* depends on the amount of polymer actually present and on what the experimental setup can detect.

