## Supplemental figures and methods for "Behaviors of individual microtubules and microtubule populations relative to critical concentrations: Dynamic instability occurs when critical concentrations are driven apart by nucleotide hydrolysis"

### SUPPLEMENTAL MATERIAL

#### Table of Contents

Supplemental Figures with Legends . . . . 2 – 14

Supplemental Methods . . . . . . . . . . . . . 15 – 17

#### SUPPLEMENTAL FIGURES AND LEGENDS

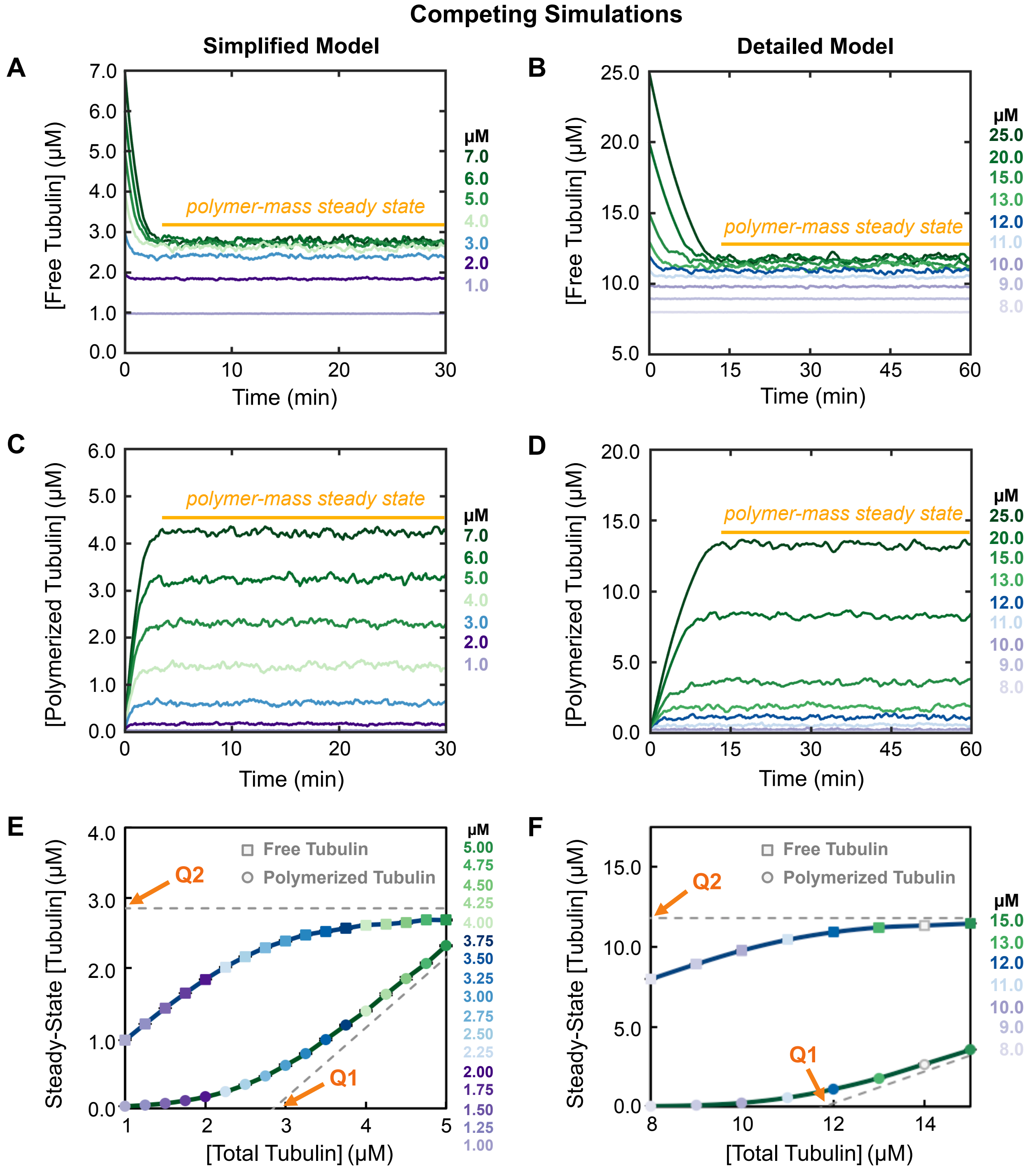


**Figure S1: Additional data relevant to Figure 3: the behavior of competing systems of MTs** (i.e., systems where [*total* tubulin] is constant). Left panels: simplified model; right panels: detailed model; colors reflect the concentrations of *total* tubulin (see color keys). ***(A-D)*** The emergent [free tubulin] (panels ***A-B***) and [polymerized tubulin] (panels ***C-D***) as functions of time from one of the three replicates of the simulation runs used in **Figure 3A-B**. ***Interpretation of panels A-D*:** Polymer-mass steady state (as shown by the horizontal line) is when [free tubulin]

and [polymerized tubulin] have reached values that stay approximately constant over time. Although there are small fluctuations around the steady-state values, there is no longer a net change over sufficient time. ***(E,F)*** Magnified regions of **Figure 3A-B**. ***Interpretation of panels E,F*****:** Detectable polymer exists at [total tubulin] below **Q1** for both the simplified and detailed simulations. Note that it can be hard to estimate the position of either **Q1** or **Q2** from the magnified region of the datasets shown here. The positions of **Q1** and **Q2** indicated here are from the full datasets shown in **Figure 3A-B**. ***Methods***: Data points in panels E-F represent the mean +/- one standard deviation of the values obtained in three independent runs of the simulations.

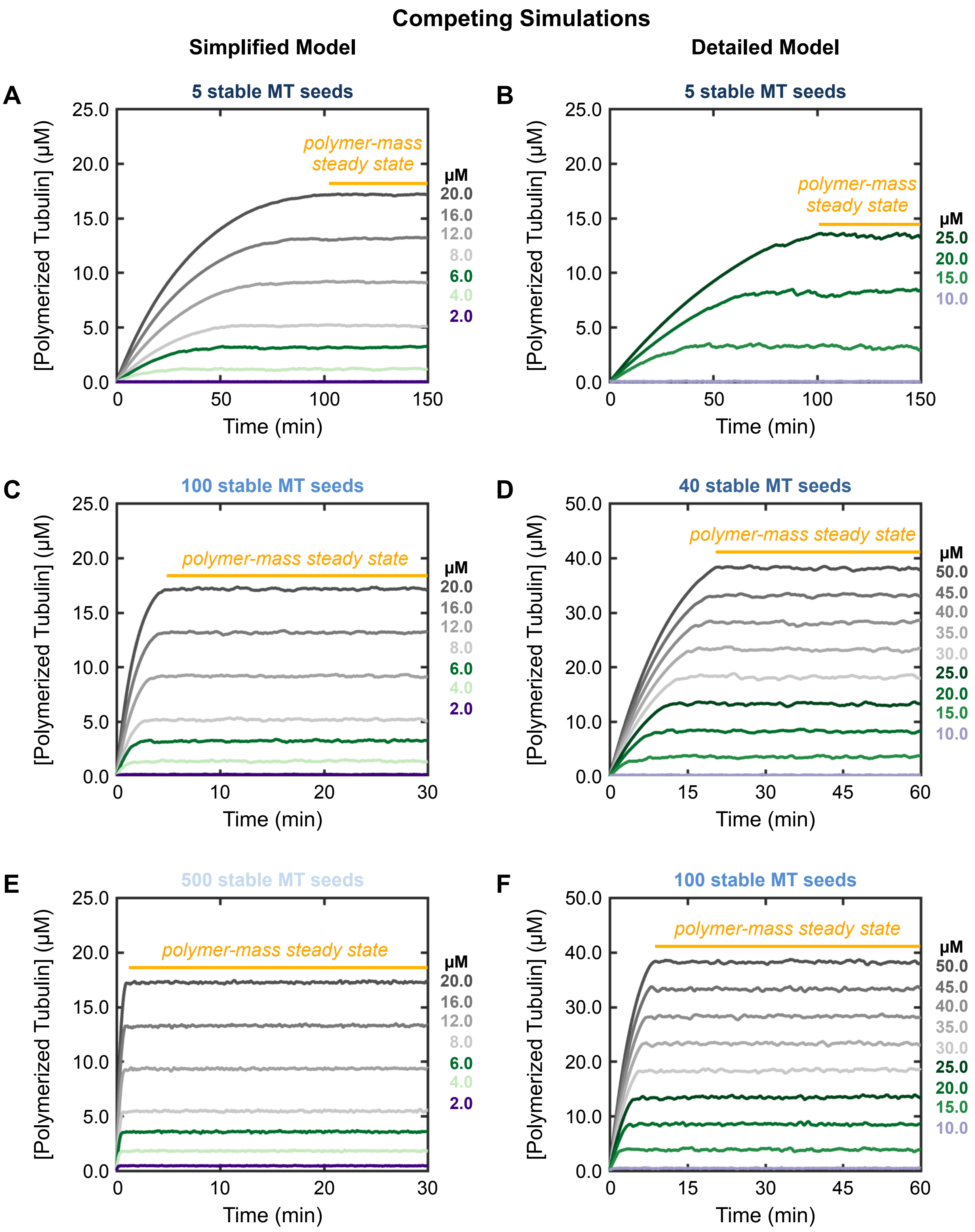


**Figure S2: Additional data relevant to Figure 4: varying the number of stable MT seeds in competing systems.** Left panels: simplified model; right panels: detailed model. Colors reflect the concentrations of *total* tubulin (see color keys). **(*A-F*)** [Polymerized tubulin] versus time, analogous to **Figure S1C-D**. Simplified model: 5 seeds (panel **A**), 100 seeds (panel **C**), 500 seeds (panel **E**). Detailed model: 5 seeds (panel **B**), 40 seeds (panel **D**), 100 seeds (panel **F**). Each plot is from one of the three replicates of the simulation runs used in **Figure 4**. ***Interpretation*:** In both models, the time to reach polymer-mass steady state is longer when there are fewer seeds or when [total tubulin] is higher.

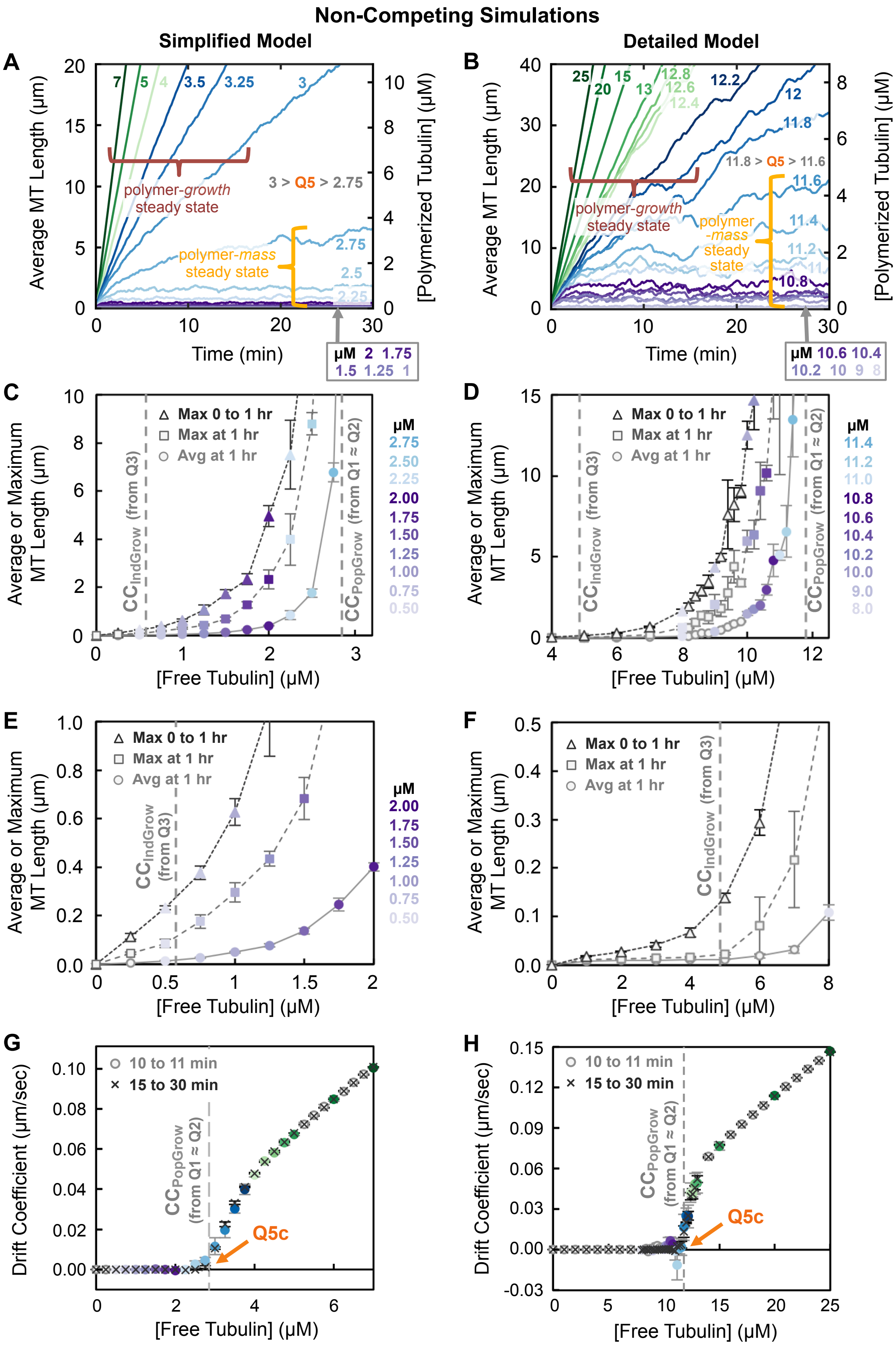


**Figure S3** ***(legend on next page)***

**Figure S3: Additional data relevant to Figure 5: the behavior of non-competing systems of MTs** (i.e., systems where [*free* tubulin] is constant). Left panels: simplified model; right panels: detailed model. Colors reflect the concentrations of *free* tubulin. ***(A,B)*** Average MT length (left axis) and [polymerized tubulin] (right axis) as functions of time from one of the three replicates of the simulation runs used in **Figure 5C-F**. ***Interpretation of panels A,B*****:** At [free tubulin] below $CC_{PopGrow}$ as measured by **Q5**, the system reaches polymer-mass steady state, where the average MT length or [polymerized tubulin] have plateaued. In contrast, at [free tubulin] above $CC_{PopGrow}$ (**Q5**), there is no polymer-mass steady state, but instead a polymer-growth steady state, where average MT length or [polymerized tubulin] increase at constant rates over time. ***(C-F)*** Average MT length of the population at 1 hour (circles). Maximum MT length of the population either at 1 hour (squares) or between 0 to 1 hour (triangles). ***Methods for panels C-F*****:** The length here is the length above the seed and the measurements include all 100 stable MT seeds in the simplified model and all 40 stable MT seed in the detailed model, so each empty seed contributes a value of 0 to the average. Panels **E-F** show zoom-ins of the data plotted in panels **C-D**. Data points represent the mean +/- one standard deviation of the values obtained in three independent runs of the simulations. ***(G,H)*** Examination of the effect of changing the total observation time when calculating the steady-state drift coefficients using the time-step analysis method (see Supplemental Methods). The plots show a comparison of the drift coefficients for MT populations observed over a 1-minute interval (o symbol) or over a 15-minute interval (x symbols, re-plotted from **Figure 5E-F**) as a function of [free tubulin]. ***Interpretation of panels G,H*****:** These data show that varying the total observation time has an impact on the noise (drift coefficients measured over the 15-minute interval show less noise, as seen by smaller error bars, than those from the 1-minute interval), but the values themselves are not affected. ***Methods for panels G,H*****:** To determine the *steady-state* value of the drift coefficient, the measurements should be performed after the system has reached the appropriate steady state (polymer-mass steady state for [free tubulin] below $CC_{PopGrow}$, and polymer-growth steady state for [free tubulin] above $CC_{PopGrow}$). Data points represent the mean +/- one standard deviation of the values obtained in three independent runs of the simulations.

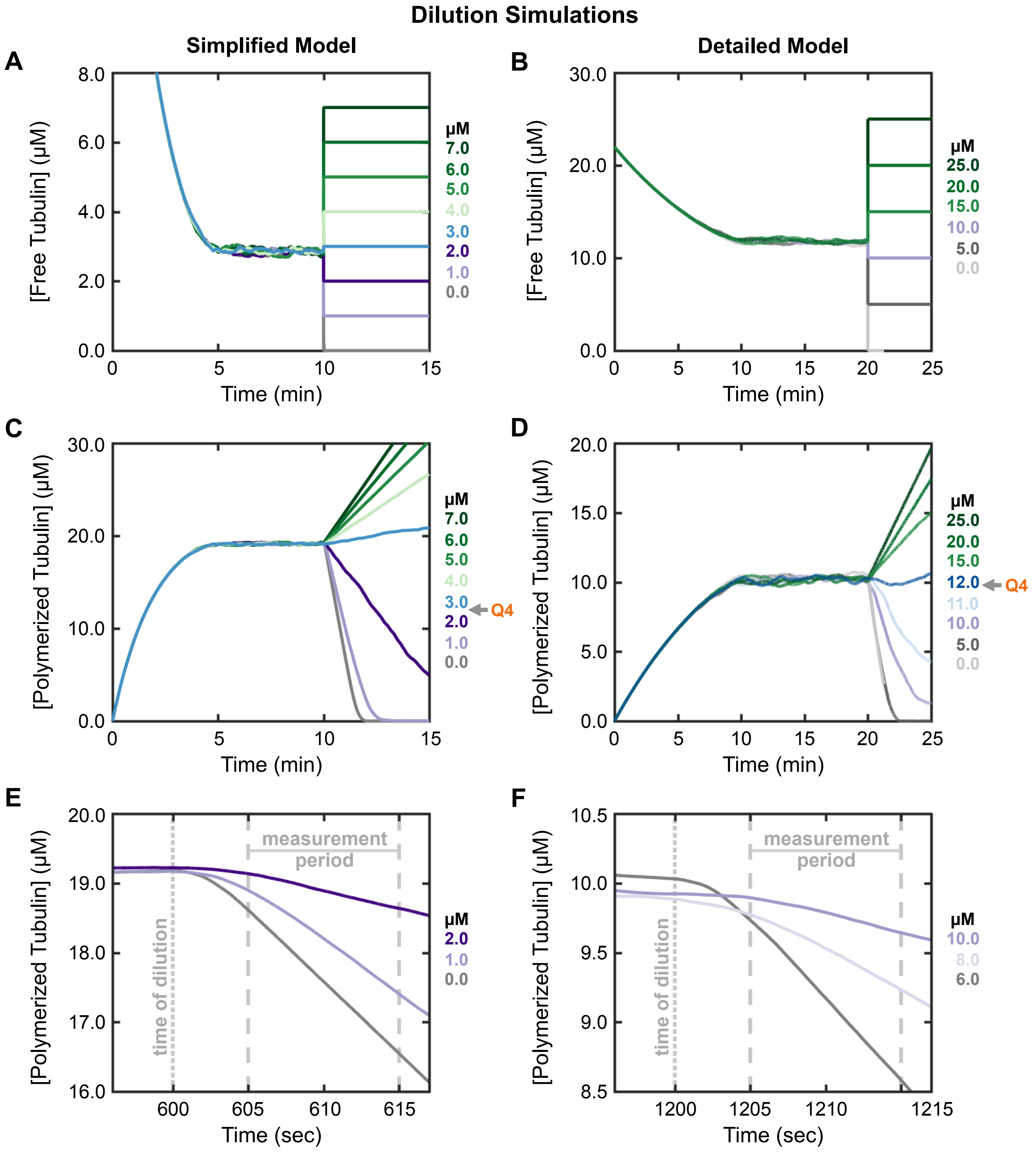


**Figure S4: Additional data relevant to Figure 6: measuring the rate of change in [polymerized tubulin], i.e., flux of tubulin into and out of polymer, in the dilution simulations.** Left panels: simplified model; right panels: detailed model. Colors reflect the *dilution [free tubulin]*, i.e., the concentration of free tubulin after the dilution. **(*A,B*)** Plots of [free tubulin] versus time for selected values of dilution [free tubulin]. **(*C,D*)** Plots of [polymerized tubulin] versus time for selected values of dilution [free tubulin]. **(*E,F*)** Plots of [polymerized tubulin] versus time for selected values of dilution [free tubulin] during a time period from shortly before the dilution through the flux measurement period from **Figure 6** (605 to 615 seconds in the simplified model; 1205 to 1215 seconds in the detailed model). ***Interpretation***: These data show that the delay after dilution allows the rate of change in [polymerized tubulin] to reach its steady-state value (as expected because the cap length needs time to respond to the new [free tubulin] (Duellberg et al., 2016; Bowne-Anderson et al., 2013)). If the measurements were performed without the delay, then the magnitude of the initial rates would be underestimated.

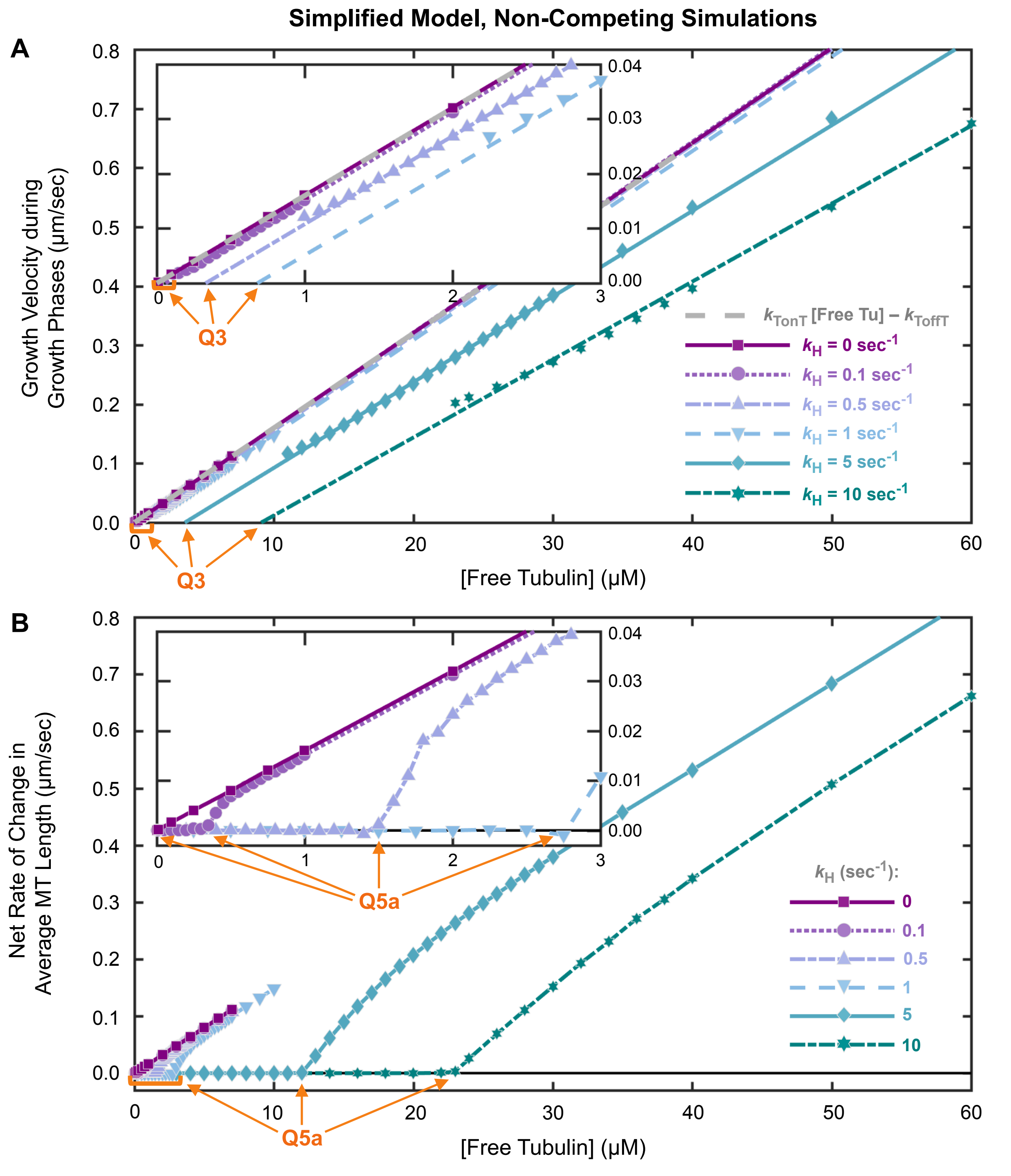


**Figure S5: Additional plots relevant to Figure 9: analysis of varying the rate constant for nucleotide hydrolysis ($k_H$) in the simplified model (non-competing simulations).** For comparison across the varying values of $k_H$: $V_g$ from each of **panels 9A-F** is re-plotted here in panel **S5A**; net rate of change in average MT length from each of **panels 9A-F** is re-plotted here in panel **S5B**. The insets show zoom-ins at low [free tubulin]. ***Interpretation***: Q3 (panel **A**) and Q5a (panel **B**) each increase as $k_H$ is increased. ***Methods:*** The regression lines (re-plotted here from **Figure 9**) were fitted in the [free tubulin] range from $CC_{PopGrow}$ to the highest [free tubulin] shown in the **Figure 9** panel for each $k_H$ value. The $V_g$ data points re-plotted here are shown only in the [free tubulin] range were $V_g$ is approximately linear as a function of [free tubulin]; for some of the $k_H$ values, this is a wider range than the range used for fitting the regression line.

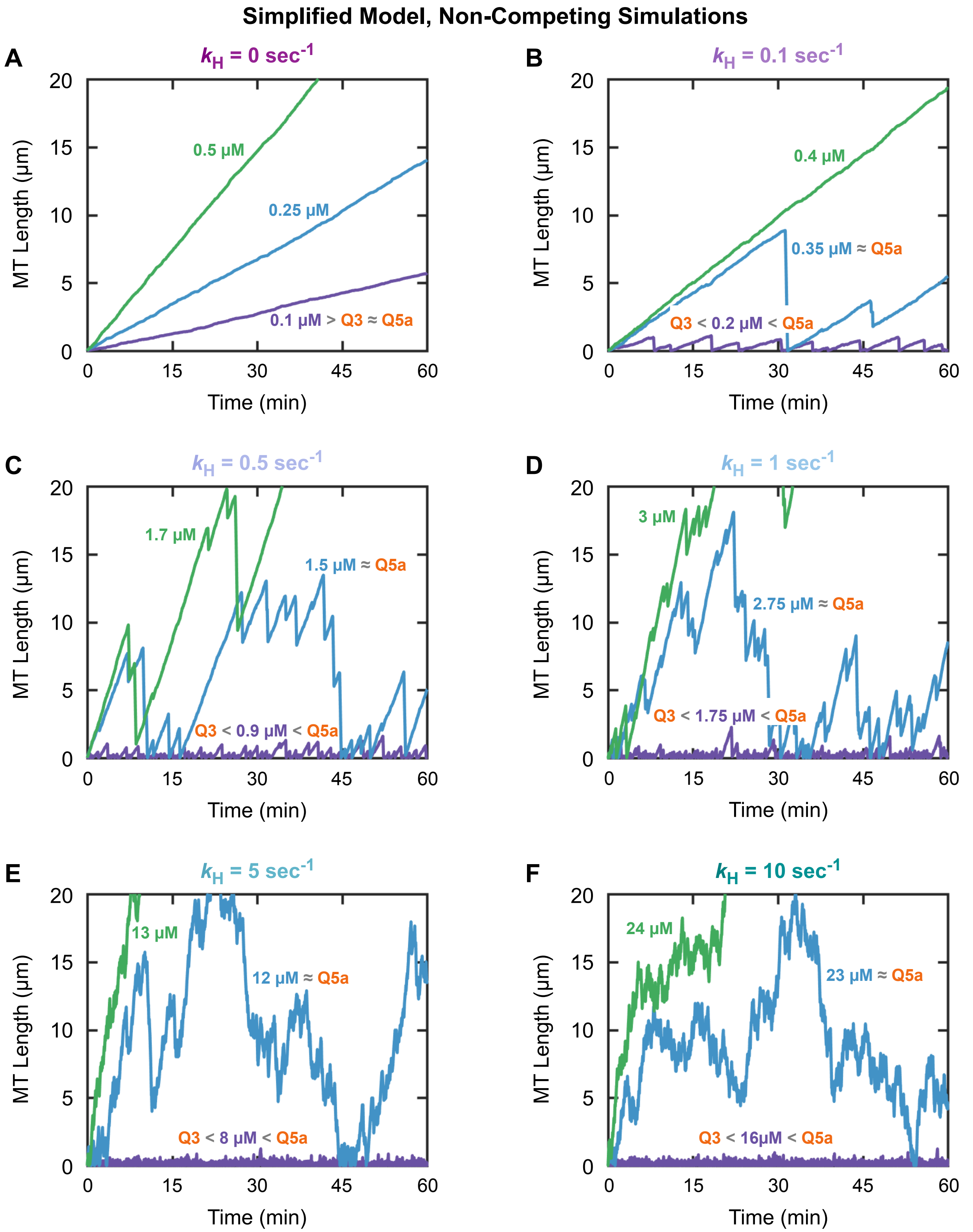


**Figure S6: Additional data relevant to Figure 9: individual MT length histories from the simplified model with varying values of the rate constant for nucleotide hydrolysis ($k_H$).** Each of panels **A-F** corresponds to a different value of $k_H$ (see panel titles). ***(A)*** For $k_H$ = 0, length histories are plotted for individual MTs at three different values of [free tubulin] above the CC ($CC_{IndGrow}$ = $CC_{PopGrow}$ when $k_H$ = 0). ***(B-F)*** For each value of $k_H$ > 0, lengths histories are plotted for individual MTs at three different values of [free tubulin]: approximately halfway between $CC_{IndGrow}$ and $CC_{PopGrow}$ (purple); near $CC_{PopGrow}$ (blue); and slightly above $CC_{PopGrow}$ (green). ***Interpretation*****:** In panel **A**, where $k_H$ = 0 (equilibrium polymer), no dynamic instability (DI) is observed. In panels **B-F**, when $k_H$ > 0 (steady-state polymer), the filaments exhibit DI. Note that for low values of $k_H$ (e.g., panel **B**), DI occurs only within a narrow range of [free tubulin]. As $k_H$ increases, DI occurs over a wider range of [free tubulin] and is therefore more likely to be observed in experiments. For intermediate $k_H$, (e.g., panels **C-D**), classical DI is evident. For very high $k_H$ (e.g., panels **E-F**), the MTs transition frequently between growth and shortening in contrast to the DI with clearer periods of extended growth and shortening observed at lower $k_H$.

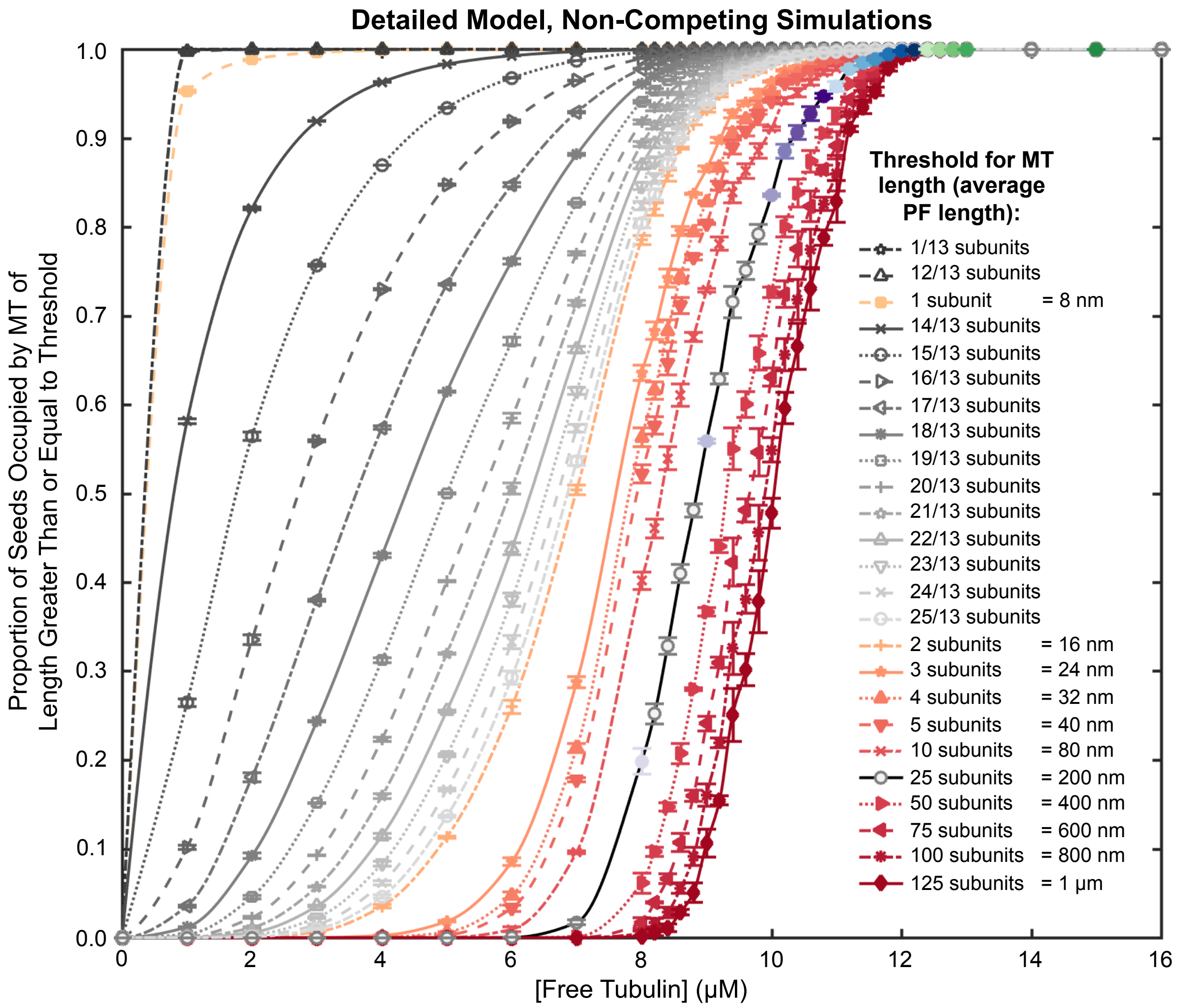


**Figure S7: Detailed model $P_{occ}$ (proportion of occupied seeds) with additional thresholds.** As in **Figure 10**, $P_{occ}$ is the fraction of the stable seeds bearing a MT with length greater than or equal to the given threshold. In the detailed model, the MT length is the average of the 13 protofilament lengths and can therefore have non-integer values. The thresholds with integer values (in units of subunit lengths) are re-plotted from **Figure 10D**. ***Interpretation***: The fractional thresholds from 14/13 subunits to 25/13 subunits shown here demonstrate how the $P_{occ}$ curve varies as the threshold changes from 1 subunit to 2 subunits. These data show that the sigmoidal shape of the $P_{occ}$ curve emerges as the detection threshold is increased. ***Methods***: All data points represent the mean +/- one standard deviation of the values obtained in three independent runs of the simulations with constant [free tubulin]. The values from each run are averages from 25 to 30 minutes.

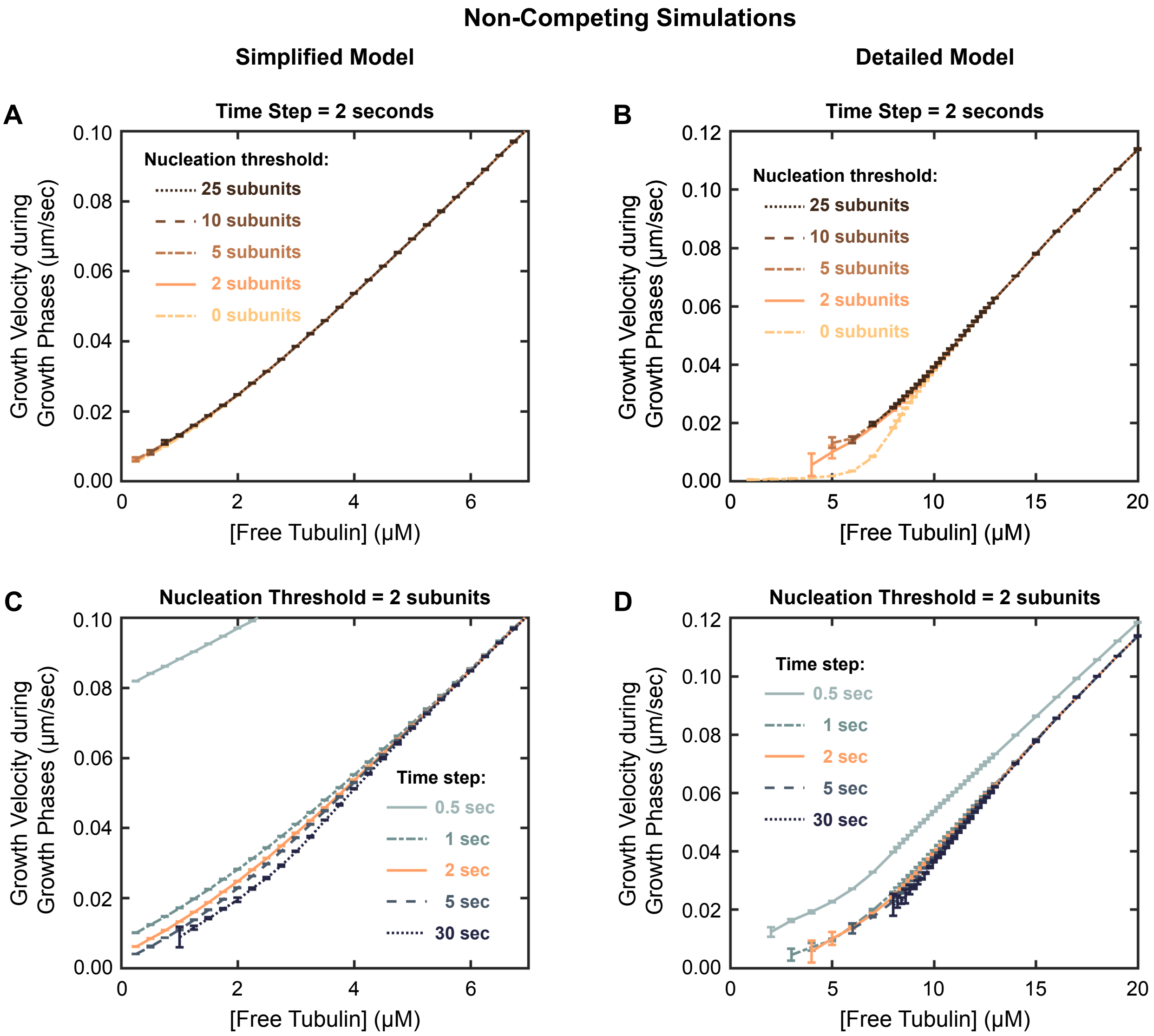


**Figure S8: Effect of time-step duration and "nucleation" threshold in the time-step method for measuring growth velocity ($V_g$) described in the Supplemental Methods.** Left panels: simplified model; right panels: detailed model. **(*A,B*)** Time step = 2 seconds, with varying nucleation thresholds (1 subunit = 8 nm). **(*C,D*)** Nucleation threshold = 2 subunits, with varying time steps. ***Interpretation*:** Based on these results of varying the nucleation threshold and the time step, we chose a time step of 2 seconds and a nucleation threshold of 2 subunits for the analysis in **Figure 7**. *Consequences of changing the nucleation threshold*: Using a lower nucleation threshold underestimates $V_g$, particularly noticeable in the detailed model (panel **B**). Using a higher nucleation threshold increases the minimum concentration at which data are obtained (panels **A-B**). *Consequences of changing the time step*: Using a shorter time step overestimates $V_g$. Using a longer time step underestimates $V_g$ and increases the minimum concentration at which data are obtained (panels **C-D**). See Supplemental Methods for additional information and interpretations.

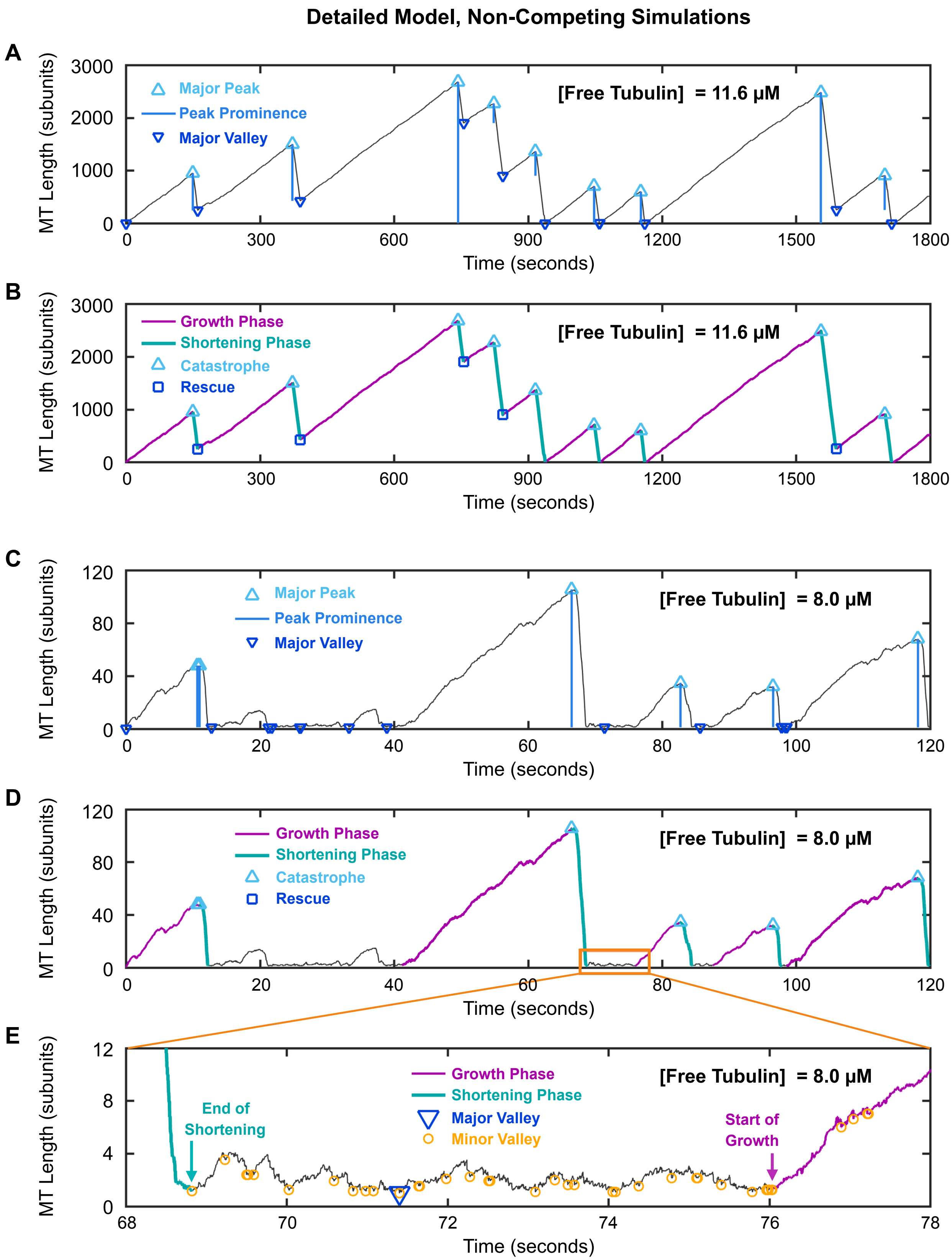


**Figure S9** ***(legend on next page)***

**Figure S9: Length histories illustrating the DI analysis method described in the Supplemental Methods.** Representative length histories from the detailed model under constant [free tubulin] conditions at 11.6 µM (panels **A**-**B**) and at 8 µM (panels **C**-**E**), chosen to highlight different aspects of DI. **(*A,C*)** Identification of major peaks and valleys (those with prominence ≥ 25 subunits = 200 nm) in length history data. The peak prominence is the vertical distance between the peak and the valley that is nearest to the peak, without a larger intervening peak (the nearest valley can be either before or after the peak). **(*B,D*)** Identification of the growth phases, shortening phases, catastrophes, and rescues. The ascent to each major peak is classified as a growth phase (purple), and the descent from each major peak is classified as a shortening phase (teal). Transitions from growth to shortening are called catastrophes (light blue triangles) and occur at major peaks. Transitions from shortening to growth, without depolymerization back to the seed, are called rescues (dark blue squares, panel **B**); we classify major valleys as rescues only if the MT length at the major valley is greater than or equal to the rescue threshold (25 subunits = 200 nm). **(*E*)** Determining the end point of a shortening segment (teal) and the start point of a growth segment (purple) when a major valley occurs below the rescue threshold (25 subunits = 200 nm). The end of a shortening phase is identified as the *first minor* valley that occurs after the MT length has shortened to within 1 subunit of the *first major* valley after the shortening phase. The start of a growth phase is identified as the *last minor* valley that is within one subunit of the major valley and before the next peak. **All panels:** The vertical axes show the MT length (average of the 13 protofilament lengths) above the stable MT seed in units of subunit length (1 subunit = 8 nm).

### SUPPLEMENTAL METHODS

#### Measuring drift coefficient from time-step analysis

The drift coefficient is a measure of the rate of displacement of the MT ends. If one end of a MT is fixed (as in our simulations), then the rate of displacement of the free end is equal to the rate of change in the MT's length. We calculated drift coefficients for our simulation data (**Figures 5E-F**, **S3G-H**) using the drift coefficient formula of (Komarova et al., 2002):

$$v_{\mathrm{d}} = \frac{\sum s_i}{\sum t_i}$$

where $v_{\mathrm{d}}$ is the drift coefficient, $s_i$ is the displacement of a MT end, and $t_i$ is the corresponding time between sequential frames in a time-lapse movie. In (Komarova et al., 2002), where this formula was applied to experimental data analyzed by subtraction of image intensities from sequential frames, the time between successive frames was 3 to 5 seconds.

To apply this approach to our simulation data, we calculated the displacements ($s_i$) of each MT end over time steps ($t_i$) of 2 seconds. For each [free tubulin] separately, we then summed these displacements of the MT ends ($\sum s_i$) over the population and over time for either 30 time steps (from minute 10 to minute 11 in the simulation; **Figure S3G-H**, o symbols) or 450 time steps (from minute 15 to minute 30 in the simulation; **Figures 5E-F** and **S3G-H**, x symbols). The sum of the displacements over the population and over time ($\sum s_i$) and the sum of the corresponding time changes ($\sum t_i$) were plugged into the above formula for $v_d$ to obtain the drift coefficient for each free tubulin concentration (**Figures 5E-F**, **S3G-H**). For example, in simulation data, if 100 MTs were measured over 30 time steps, then $\sum s_i$ and $\sum t_i$ would each be the sum of 100*30 = 3000 values. For experimental data, the same number of MTs would not necessarily be observed in every frame, so the sums would just include all displacements that are measured.

When applying the time-step analysis method to experimental data, the displacements that can be detected would depend on the experimental imaging resolution. For our simulated data, we did not impose any minimum on the size of displacement (the magnitude of $s_i$) during each time step. Also, for the simulated data, the size of the time step does *not* affect the resulting value of drift coefficient, because the length of every MT is known at every time step. In contrast, for experimental data, the size of the time step could have some effect on the results, because it may affect whether a MT is detected in two consecutive frames.

To identify Q5c (**Figures 5E-F**, **S3G-H**), the measurements should be taken during a time period when the drift coefficients have reached their steady-state values (this will occur at polymer-mass steady state for [free tubulin] below Q5 and at polymer-growth steady state for [free tubulin] above Q5). The time-step method described here could also be used to measure drift coefficients at earlier times if one wishes to examine the approach to steady state.

#### Measuring $V_g$ from time-step analysis

The time-step analysis described above can also be used to obtain measurements of $V_g$ (**Figure 7**, square symbols). To do so, the sums are restricted to include only positive displacements:

$$V_g = \frac{\sum_{pos} s_i}{\sum_{pos} t_i}$$

where $\sum_{pos} s_i$ is the sum of all displacements satisfying $s_i > 0$ and $\sum_{pos} t_i$ is the sum of the corresponding time changes. (When applying this method to experiments with physical detection limits, the sum would include only displacements satisfying $s_i \geq$ detection limit.)

When applying the time-step analysis to the simulation data, we observed that $V_g$ could be underestimated if there were time steps during which the length of a MT remained close to zero but displayed a small positive increase in length. We therefore imposed a “nucleation” threshold and counted a displacement only if the length of the MT above the seed was greater than or equal to the threshold for the entire time step (**Figure S8A-B**). Note, this is not the same as imposing a threshold on the size of the displacement itself; whenever the MT length was above the nucleation threshold, all positive displacements ($s_i > 0$) were counted.

Additionally, the value of $V_g$ from the time-step analysis *does* depend on the size of the time step (**Figure S8C-D**). If the time step is too small, the results will be affected by the velocity of upward fluctuations that are small and rapid, and the method will therefore overestimate the actual velocity of the extended growth phases of DI. If the time step is too large, there will be individual displacements that include some shortening (despite being net positive), and the method will therefore underestimate $V_g$. Our results indicate that time steps within an intermediate range produce similar results to each other (e.g., time steps ~2-5 seconds in simplified model and ~1-5 seconds in detailed model; **Figure S8C-D**).

The time-step method could also be used to estimate $V_s$ by restricting to only negative displacements. However, accurate measurement of $V_s$ would require using a smaller time step than accurate measurement of $V_g$, because shortening phases tend to have faster velocities and last for shorter amounts of time than growth phases.

#### Automated quantitative analysis of dynamic instability

For the analyses in **Figures 7, 9,** and **S5**, it is necessary to identify periods of growth and shortening in DI data and to measure the DI parameters ($V_g$ = growth velocity during growth phases, $V_s$ = shortening velocity during shortening phases, $F_{cat}$ = frequency of catastrophe, and $F_{res}$ = frequency of rescue). To obtain these data, we developed an automated MATLAB program (code available upon request). This automated program was applied here to simulated data, but can be applied to any length history data.

*Overview of the DI analysis method:* The analysis program uses the MATLAB function 'findpeaks' to identify major peaks in the length history data with a user-defined threshold for peak prominence (**Figure S9A,C**). The peak prominence is the height of the peak relative to the valley that is nearest to the peak, without a larger intervening peak (the nearest valley can be either before or after the peak)**.** The ascent to each major peak is classified as a growth phase, and the descent from each major peak is classified as a shortening phase. The point of the transition from growth to shortening at a major peak is identified as a catastrophe (**Figure S9B,D**). A transition from shortening to growth is considered to be a rescue *only if* the MT length at the transition is greater than a user-defined threshold (**Figure S9B**).

The DI parameters are calculated as follows, where the totals are over all detected phases (of growth or shortening, as indicated) for all individuals in the population:
$V_g$ = total length change during growth phases / total time spent in growth phases,
$V_s$ = total length change during shortening phases / total time spent in shortening phases,
$F_{cat}$ = total number of catastrophes / total time spent in growth phases,
$F_{res}$ = total number of rescues / total time spent in shortening phases.

*Finding major peaks, major valleys, and minor valleys*: The first step in the analysis uses the MATLAB function 'findpeaks' to identify major peaks in the length history data that have a peak prominence of at least 25 subunits (200 nm) (**Figure S9A,C**). Next, the function 'findpeaks' is applied to the negative of the length data to identify *major* valleys, with prominence of at least 25 subunits, and *minor* valleys, with prominence of at least 0.5 subunits. (In the detailed model, the outputted MT length is the average of the lengths of the 13 protofilaments, so the MT length can have non-integer values).

*Determining the start and end points of each growth/shortening segment*: At a catastrophe event, the end of the growth phase and the start of the shortening phase are chosen to be the same point in time as each other (at a major peak) (**Figure S9A-D**). Similarly, at a rescue event, the end of shortening and the start of growth are identified as occurring at the same time point (at a major valley) (**Figure S9A-B**). However, if the MT length at a major valley is below the rescue threshold (25 subunits), then the start of the growth phase may be at a later time than the end of the preceding shortening phase (**Figure S9C-E**). In this case, the procedure described in the figure legend is used to choose the points to be identified as the end of the shortening phase and the start of the growth phase.
